# DCAF16-Based Covalent Handle for the Rational Design of Monovalent Degraders

**DOI:** 10.1101/2024.02.20.580683

**Authors:** Melissa Lim, Thang Do Cong, Lauren M. Orr, Ethan S. Toriki, Andrew C. Kile, James W. Papatzimas, Elijah Lee, Daniel K. Nomura

## Abstract

Targeted protein degradation with monovalent molecular glue degraders is a powerful therapeutic modality for eliminating disease causing proteins. However, rational design of molecular glue degraders remains challenging. In this study, we sought to identify a transplantable and linker-less covalent handle that could be appended onto the exit vector of various protein-targeting ligands to induce the degradation of their respective targets. Using the BET family inhibitor JQ1 as a testbed, we synthesized and screened a series of covalent JQ1 analogs and identified a vinylsulfonyl piperazine handle that led to the potent and selective degradation of BRD4 in cells. Through chemoproteomic profiling, we identified DCAF16 as the E3 ligase responsible for BRD4 degradation—an E3 ligase substrate receptor that has been previously covalently targeted for molecular glue-based degradation of BRD4. Interestingly, we demonstrated that this covalent handle can be transplanted across a diverse array of protein-targeting ligands spanning many different protein classes to induce the degradation of CDK4, the androgen receptor, BTK, SMARCA2/4, and BCR-ABL/c-ABL. Our study reveals a DCAF16-based covalent degradative and linker-less chemical handle that can be attached to protein-targeting ligands to induce the degradation of several different classes of protein targets.

## Introduction

Monovalent molecular glue degraders have arisen as a powerful therapeutic modality for degrading therapeutic targets of interest through inducing the proximity of an E3 ubiquitin ligase with a neo-substrate protein to ubiquitinate and degrade the target through the proteasome ^1,2^. Molecular glue degraders are potentially more promising compared to heterobifunctional Proteolysis Targeting Chimeras (PROTACs) because of their lower molecular weights and associated drug-like properties, as well as their potential to exploit shallow protein-protein interfaces between an E3 ligase and less tractable therapeutic proteins that may not possess deep binding pockets ^1^. However, most molecular glue degraders have either been discovered fortuitously or through phenotypic screens ^1,3–7^. Rational chemical design of molecular glue or monovalent degraders in a target-based manner remains challenging.

Many recent studies have reported how subtle chemical alterations to otherwise non-degradative small-molecule inhibitors converted them into molecular glue degraders of their respective targets ^6–9^. These studies gave rise to the exciting possibility of transplantable chemical handles that could be appended onto the exit vector of protein-targeting ligands to convert these compounds into molecular glue degraders of their targets. E3 ligases have been shown to be ligandable with covalent small-molecules and chemoproteomic approaches ^10–16^. Covalent handles have also been successfully used in heterobifunctional PROTACs to identify permissive chemical handle and ligandable E3 ligase pairs that can be exploited for targeted protein degradation applications. These studies have identified various covalent handles targeting cysteines in E3 ligase substrate receptors DCAF16 and DCAF11 ^13,17,18^. Covalent ligand screens against specific ubiquitin proteasome system components have also yielded new E3 ligase, E2 ubiquitin conjugating enzyme, or Cullin adaptor recruiters against RNF114, RNF4, FEM1B, UBE2D, DDB1, and SKP1 that can be used for PROTACs ^12,19–24^.

Recent studies have also revealed that covalent chemistry can be used to identify potential chemical handles that enable the rational design of monovalent or molecular glue degraders. We previously discovered a covalent chemical handle that targets a cysteine in the quality control E3 ligase RNF126 that could be appended to the exit vector of a diverse range of protein-targeting ligands without the necessity for a linker to enable degradation of their respective targets ^25^. Covalent molecular glue degraders have also been discovered that enhance weak existing interactions between DCAF16 and BRD4 to degrade BRD4 in a template-assisted covalent modification approach ^26,27^.

In this study, we sought to identify additional transplantable covalent chemical handles that can convert non-degradative inhibitors into molecular glue or monovalent degraders of their respective targets. We have identified a vinylsulfonyl piperazine handle that acts through targeting a cysteine within DCAF16 to not only enable the degradation of BRD4, but also several additional neo-substrates.

## Results

### Identifying Covalent Handles that Enable the Degradation of BRD4

To identify covalent chemical handles that could convert non-degradative inhibitors into molecular glue or monovalent degraders of their targets, we used the BET family inhibitor JQ1 as a testbed to generate a series of covalent JQ1 analogs bearing various electrophilic handles that could react with cysteines, lysines, or other nucleophilic amino acids on E3 ligases to degrade BRD4 **(Figure 1a)**. Among the 18 derivatives tested, we only identified one compound, ML 1-50, that led to the loss of both the long and short isoforms of BRD4 in HEK293T cells **(Figure 1b-1c; Figure 2a)**. ML 1-50 reduced BRD4 levels in a dose-dependent manner with preferential degradation of the short BRD4 isoform with nanomolar potency in HEK293T cells **(Figure 2b-2c)**. We observed hook effects with degradation of the long BRD4 isoform. ML 1-50 only showed modest cell viability impairments in HEK293T cells at the highest concentration of 10 μM tested **(Figure S1a)**. This BRD4 degradation was attenuated by pre-treatment with either proteasome inhibitor or NEDD8-activating enzyme inhibitor MLN4924 **(Figure 2d-2g)**. We had previously observed preference for degradation of the long versus short isoforms of BRD4 with covalent PROTACs that appeared be specific to HEK293T cells compared to other cell lines ^23,24^. Similarly, we found that ML 1-50 potently degraded both the long and short BRD4 isoforms in the MDA-MB-231 breast cancer cell line, with no hook effects observed **(Figure 2h, Figure S1b)**. Quantitative proteomic profiling of ML 1-50 in MDA-MB-231 cells showed relatively selective BRD4 degradation with only 11 other proteins that were significantly reduced in levels by greater than 2-fold **(Figure 2i; Table S1)**.

**Figure 1.**
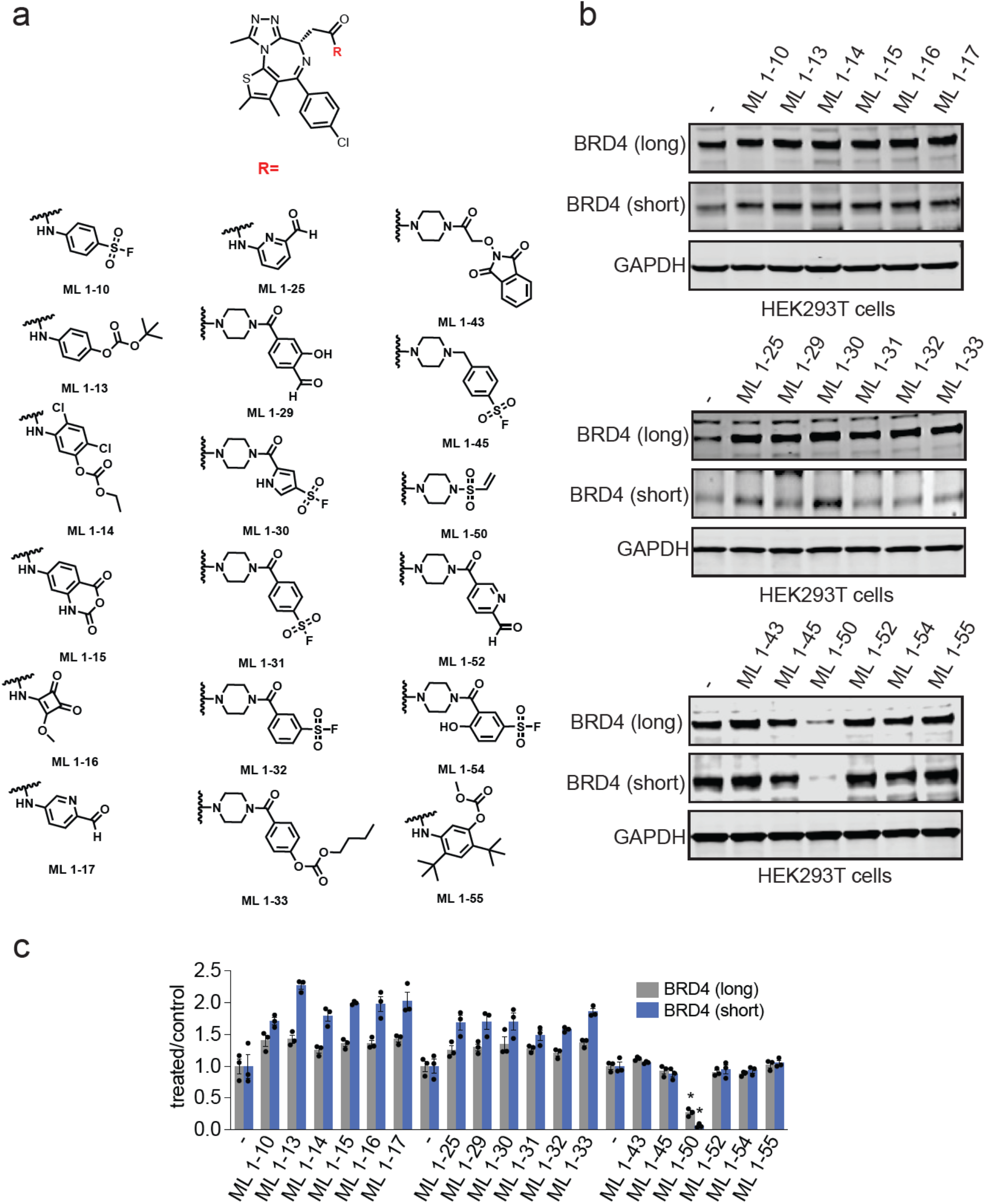
Identifying Covalent Handles that Enable the Degradation of BRD4. **(a)** Series of analogs of the BET family inhibitor JQ1 bearing various electrophilic handles. **(b)** Testing for BRD4 degradation with covalent JQ1 derivatives. HEK293T cells were treated with DMSO vehicle or covalent JQ1 derivatives (10 µM) for 24 h and BRD4 long and short isoforms and GAPDH loading control levels were assessed by Western blotting. Shown are gels that are representative of n=3 biologically independent replicates per group. **(c)** Quantitation of BRD4 long and short isoforms from experiment described in **(b)** showing individual replicate values and average ± sem. Significance is expressed as ^*^p<0.001 compared to vehicle-treated controls.

**Figure 2.**
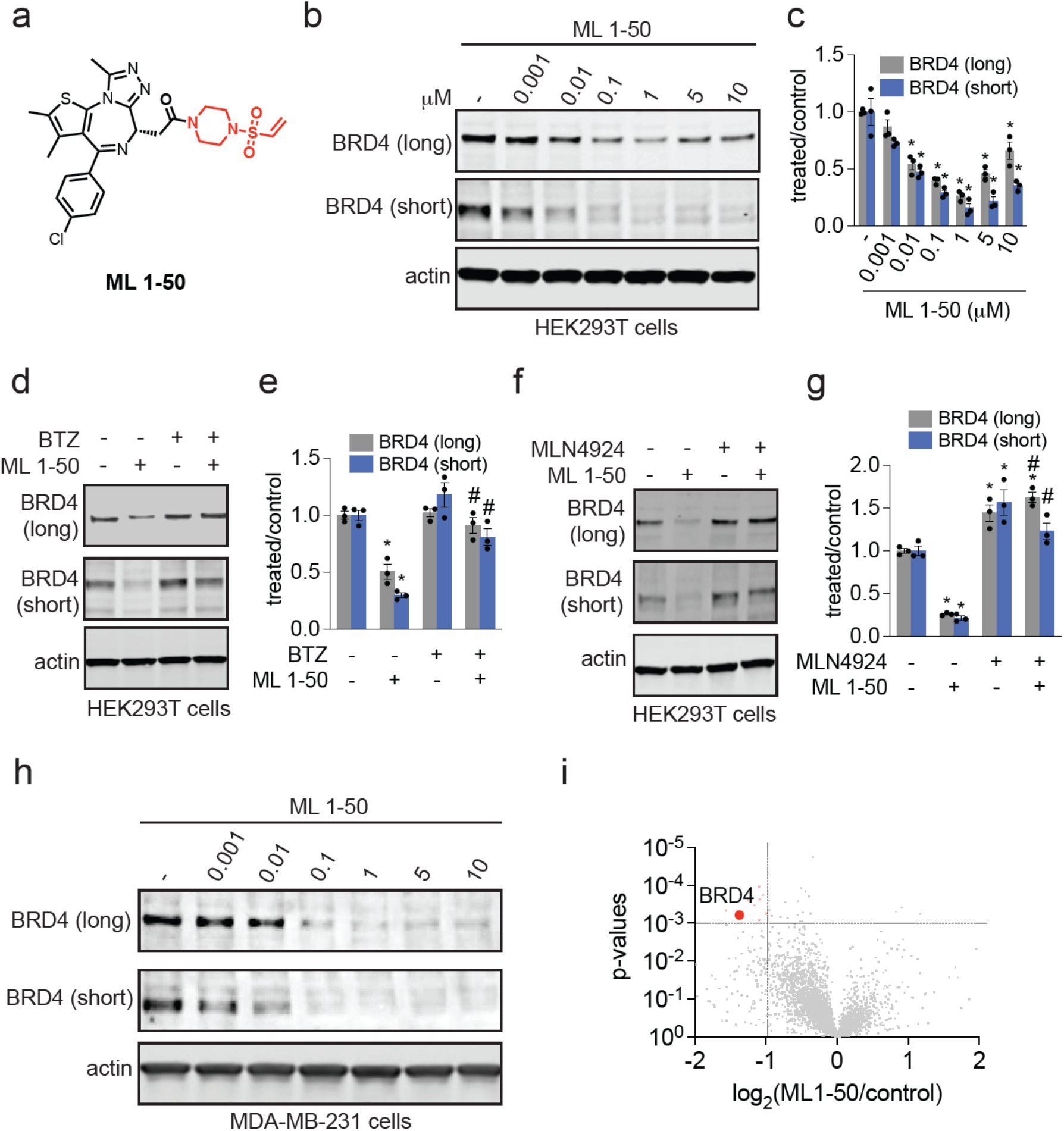
Characterization of the Monovalent and Covalent BRD4 Degrader ML 1-50. **(a)** Structure of ML 1-50 with the vinylsulfonyl piperazine covalent chemical handle in red. **(b, c)** Dose-response of BRD4 degradation. HEK293T cells were treated with DMSO vehicle or ML 1-50 for 24 h. BRD4 and actin loading control levels were assessed by Western blotting and quantified in **(c). (d, e)** Proteasome inhibitor attenuation of BRD4 degradation. HEK293T cells were pre-treated with DMSO vehicle or bortezomib (1 µM) 1 h prior to DMSO vehicle or ML 1-50 (1 µM) treatment for 24 h. BRD4 and actin loading control levels were assessed by Western blotting and quantified in **(e). (f, g)** NEDD8 activating enzyme inhibitor attenuation of BRD4 degradation. HEK293T cells were pre-treated with DMSO vehicle or MLN4924 (1 µM) 1 h prior to DMSO vehicle or ML 1-50 (1 µM) treatment for 24 h. BRD4 and actin loading control levels were assessed by Western blotting and quantified in **(g). (h)** BRD4 degradation in MDA-MB-231 cells. MDA-MB-231 cells were treated with DMSO vehicle or ML 1-50 for 24 h and BRD4 and actin loading control levels were assessed by Western blotting. **(i)** Tandem mass tagging (TMT)-based quantitative proteomic profiling of ML 1-50 in MDA-MB-231 cells. MDA-MB-231 cells were treated with DMSO vehicle or ML 1-50 (1µM) for 24 h. Proteins that were lowered in levels by >2-fold with p<0.001 are highlighted in red with BRD4 specifically labeled. Data are from n-3 biologically independent replicates per group. Blots shown in **(b, d, f, h)** are representative of n=3 biologically independent replicates per group. Bar graphs in **(c, e, g)** show average ± sem. Significance is expressed as ^*^p<0.05 compared to vehicle-treated controls and #p<0.05 compared to ML 1-50 treatment alone.

### Mapping the E3 Ligase Responsible for BRD4 Degradation

We next sought to identify the E3 ligase responsible for the BRD4 degradation observed with ML 1-50. We synthesized an alkyne-functionalized probe based on the vinylsulfonyl piperazine handle, ML 2-33 **(Figure 3a)**. Given the simplicity of this handle without a more elaborated ligand attached, we surmised that the handle would likely be more promiscuous compared to ML 1-50. We thus performed a competitive pulldown chemoproteomic experiment from ML 2-33-treated cells searching for proteins that were significantly outcompeted by ML 1-50 pre-treatment. Among these outcompeted targets, there was only one E3 ligase that was part of the Cullin E3 ligase family that could be regulated by NEDD8—DCAF16 **(Figure 3b; Table S2)**. Using competitive activity-based protein profiling (ABPP), we showed significant engagement of C119 of DCAF16 in cells by ∼28 % **(Figure S1c; Table S3)**. This modest degree of E3 ligase engagement is consistent with previous reports with covalent PROTACs showing that only a small fraction of the E3 ligase needs to be engaged to enable degradation of target proteins ^12,13,17^. Site of modification analysis by liquid chromatography-mass spectrometry (LC-MS/MS) on tryptic digests of pure DCAF16 labeled with ML 1-50 also showed a single labeled site on C119 **(Figure S1d)**.

**Figure 3.**
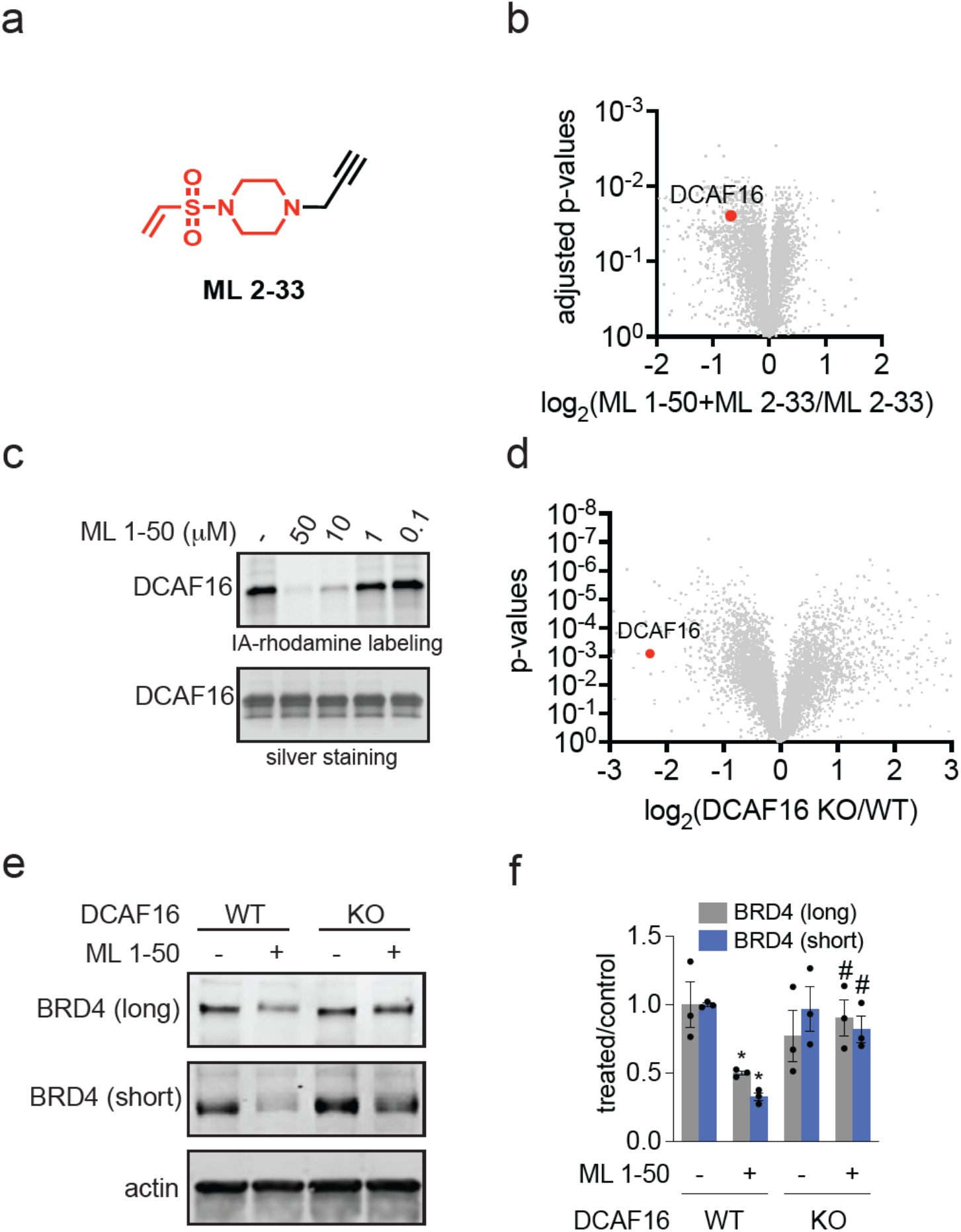
Identifying the E3 Ligase Responsible for ML 1-50-Mediated BRD4 Degradation. **(a)** Structure of alkyne-functionalized probe of the vinylsulfonyl piperazine handle (highlighted in red). **(b)** ML 1-50-outcompeted targets enriched by ML 2-33. HEK293T cell lysate were pre-treated with DMSO vehicle or ML 1-50 (200 µM) 1 h prior to treatment with the ML 2-33 probe (20 µM). Probe-modified proteins were subjected to copper-catalyzed azide alkyne cycloaddition (CuAAC) with an azide-functionalized biotin enrichment handle. Probe-modified proteins were avidin-enriched, tryptically digested, and analyzed by TMT-based proteomics. Among the significantly outcompeted targets, DCAF16 highlighted in red was the only Cullin E3 ligase substrate receptor identified. **(c)** Gel-based ABPP of ML 1-50 against pure DCAF16. Pure DCAF16 protein was pre-incubated with DMSO vehicle or ML 1-50 for 30 min prior to addition of a rhodamine-functionalized cysteine-reactive iodoacetamide probe (IA-rhodamine) (250 nM) for 1 h. Proteins were resolved by SDS/PAGE and assessed by in-gel fluorescence and protein loading was assessed by silver staining. **(d)** TMT-based quantitative proteomic analysis of DCAF16 wild-type (WT) versus knockout (KO) cells. Because there was no commercial DCAF16 antibody, proteomic methods were used to confirm DCAF16 knockout. DCAF16 is labeled in red. **(e, f)** BRD4 degradation in DCAF16 WT and KO cells. DCAF16 WT and KO cells were treated with DMSO vehicle or ML 1-50 (1 µM) for 24 h and BRD4 and actin loading control levels were assessed by Western blotting and quantified in **(f)**. Proteomics experiments and blots in **(b, c, d, e)** are from n=3 biologically independent replicates per group and blots are representative. Bar graph in **(f)** shows individual replicate values average ± sem. Significance is expressed as ^*^p<0.05 compared to vehicle-treated controls and #p<0.05 compared to ML 1-50 treated DCAF16 WT cells.

To further confirm direct interaction of ML 1-50 with DCAF16, we showed that ML 1-50 displaced cysteine-reactive probe labeling of pure DCAF16 protein by gel-based ABPP approaches without causing any precipitation of the protein **(Figure 3c)**. Consistent with the role of DCAF16 in our observed effects, we showed that the BRD4 degradation from ML 1-50 treatment was significantly attenuated in DCAF16 knockout cells compared to wild-type cells **(Figure 3d-3f; Table S4)**.

Given that several previous studies have identified DCAF16 as the E3 ligase substrate receptor responsible for the degradation of covalent BRD4 degraders bearing various different types of electrophilic handles, in part because of the native weak affinity between BRD4 and DCAF16 ^13,18,26,27^, we next determined whether our previously discovered covalent monovalent BRD4 degrader, JP-2-197, bearing a but-2-ene, 1,4-dione “fumarate” covalent degrader handle that targets RNF126 instead acts through DCAF16 ^25^. We showed that the BRD4 degradation observed by JP-2-197 was not mitigated at all in DCAF16 knockout cells **(Figure 4a-4b)**. In contrast, we observed complete and significant attenuation of BRD4 degradation in RNF126 knockout cells **(Figure 4c-4d)**. Furthermore, we demonstrated that ML 1-50-mediated degradation of BRD4 is not diminished in RNF126 KO cells compared to WT cells **(Figure 4e-4f)**. Our data thus indicate that different electrophilic degraders against the same target can act through distinct E3 ligases.

**Figure 4.**
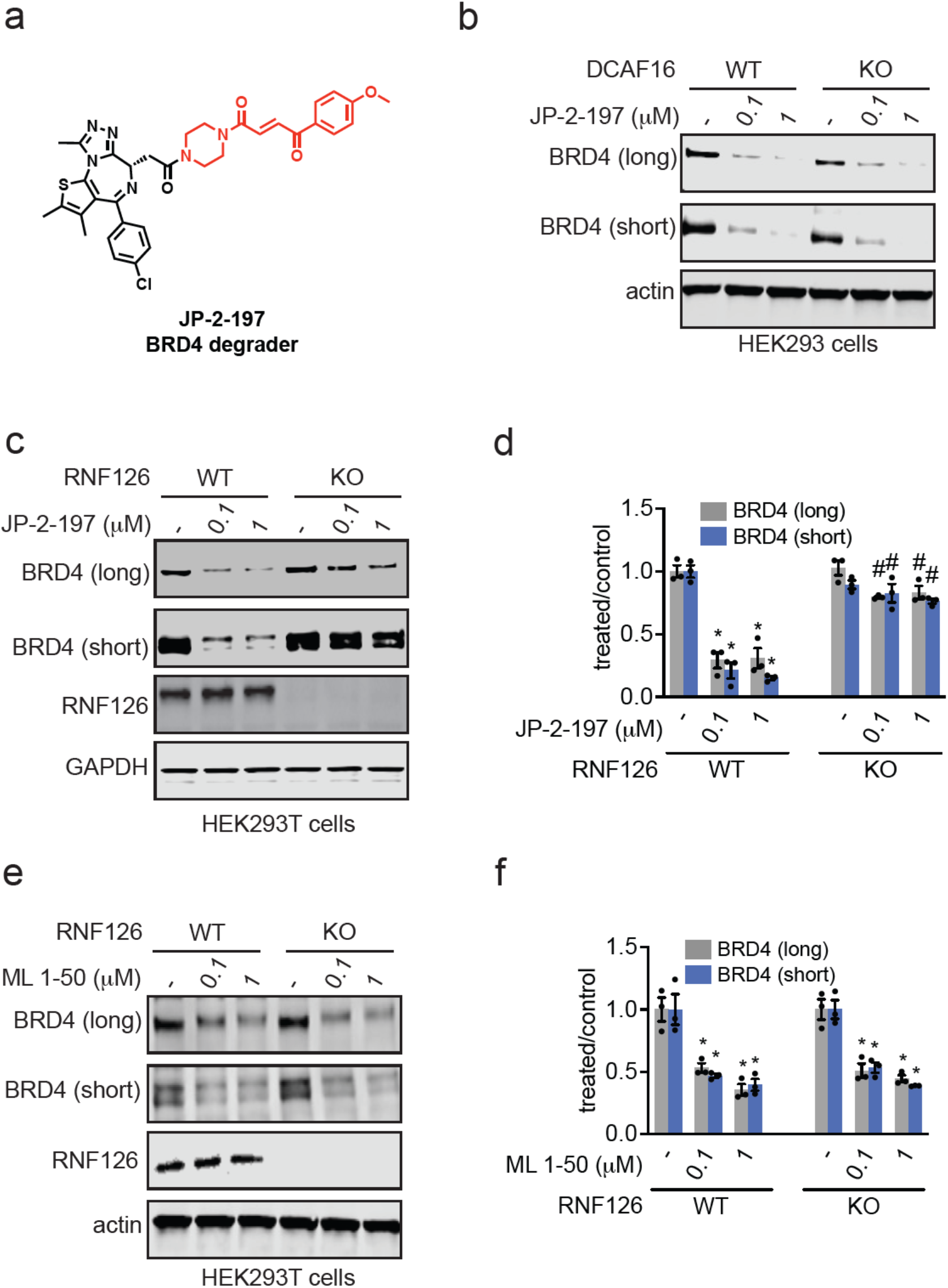
Testing the Dependence of Covalent Monovalent BRD4 Degraders on DCAF16 versus RNF126. **(a)** Structure of our previously published covalent monovalent BRD4 degrader JP-2-197 bearing the covalent “fumarate” handle shown in red. **(b)** BRD4 degradation in DCAF16 WT and KO cells. DCAF16 WT and KO HEK293 cells were treated with DMSO vehicle or JP-2-197 for 24 h and BRD4 and actin loading control levels were assessed by Western blotting. **(c, d)** BRD4 degradation in RNF126 WT and KO cells. RNF126 WT and KO HEK293T cells were treated with JP-2-197 for 10 h and BRD4, RNF126, and GAPDH loading control levels were assessed by Western blotting and quantified in **(d). (e, f)** BRD4 degradation in RNF126 KO cells. RNF126 WT and KO HEK293T cells were treated with ML 1-50 for 16 h and BRD4, RNF126, and actin loading control levels were assessed by Western blotting and quantified in **(f)**. Blots in **(b, c, e)** are representative of n=3 biologically independent replicates per group. Bar graph in **(d, f)** shows individual replicate values and average ± sem. Significance is expressed as ^*^p<0.05 compared to vehicle-treated controls and #p<0.05 compared to JP-2-197 or ML 1-50 treated RNF126 WT cells.

### Exploring the Applicability of the Covalent Handle Against Other Target Proteins

While we identified a covalent handle that can convert the non-degradative JQ1 into a degrader of BRD4, BRD4 is one of the easiest proteins to degrade and thus demonstrating proof-of-concept of a covalent handle that can degrade BRD4 does not speak to the broader applicability of this handle for other targets. Furthermore, previous studies have already demonstrated covalent and noncovalent BRD4 molecular glue degraders that act through DCAF16, through strengthening already existing weak interactions between DCAF16 and BRD4 ^8,9,26,28^. Previous studies have also used covalent DCAF16 recruiters to generate heterobifunctional PROTACs against other targets beyond BRD4 ^13^. Whether a DCAF16-targeting covalent handle could be broadly transplanted across other protein-targeting ligands to generate linker-less monovalent degraders of neo-substrate proteins beyond BRD4 is unknown. We first appended the vinylsulfonyl piperazine handle onto the clinically approved CDK4/6 inhibitor ribociclib to generate ML 1-71 **(Figure 5a)**. ML 1-71 significantly degraded CDK4 in cells in a dose-dependent manner, albeit less potently compared to ML 1-50 and BRD4 **(Figure 5b-5c)**. Despite the modest potency of this degrader, we still observed significant attenuation of CDK4 degradation in DCAF16 knockout cells **(Figure 5d-5e)**. Quantitative proteomic profiling of ML 1-71 in C33A cervical cancer cells also demonstrated relatively selective CDK4 degradation with only 16 other targets reduced in levels that may arise from transcriptional effects downstream of CDK4 inhibition or off-target effects **(Figure 5f; Table S5)**. We next generated a monovalent degrader for SMARCA2/4 bearing the vinylsulfonyl piperazine handle—ML 1-96 **(Figure S2a)** ^29^. ML 1-96 dose-responsively and significantly degraded both SMARCA2 and SMARCA4 in MV-4-11 leukemia cancer cells **(Figure S2b-S2c)**. This SMARCA2/4 loss was once again attenuated in DCAF16 knockout cells **(Figure S2d-S2e)**.

**Figure 5.**
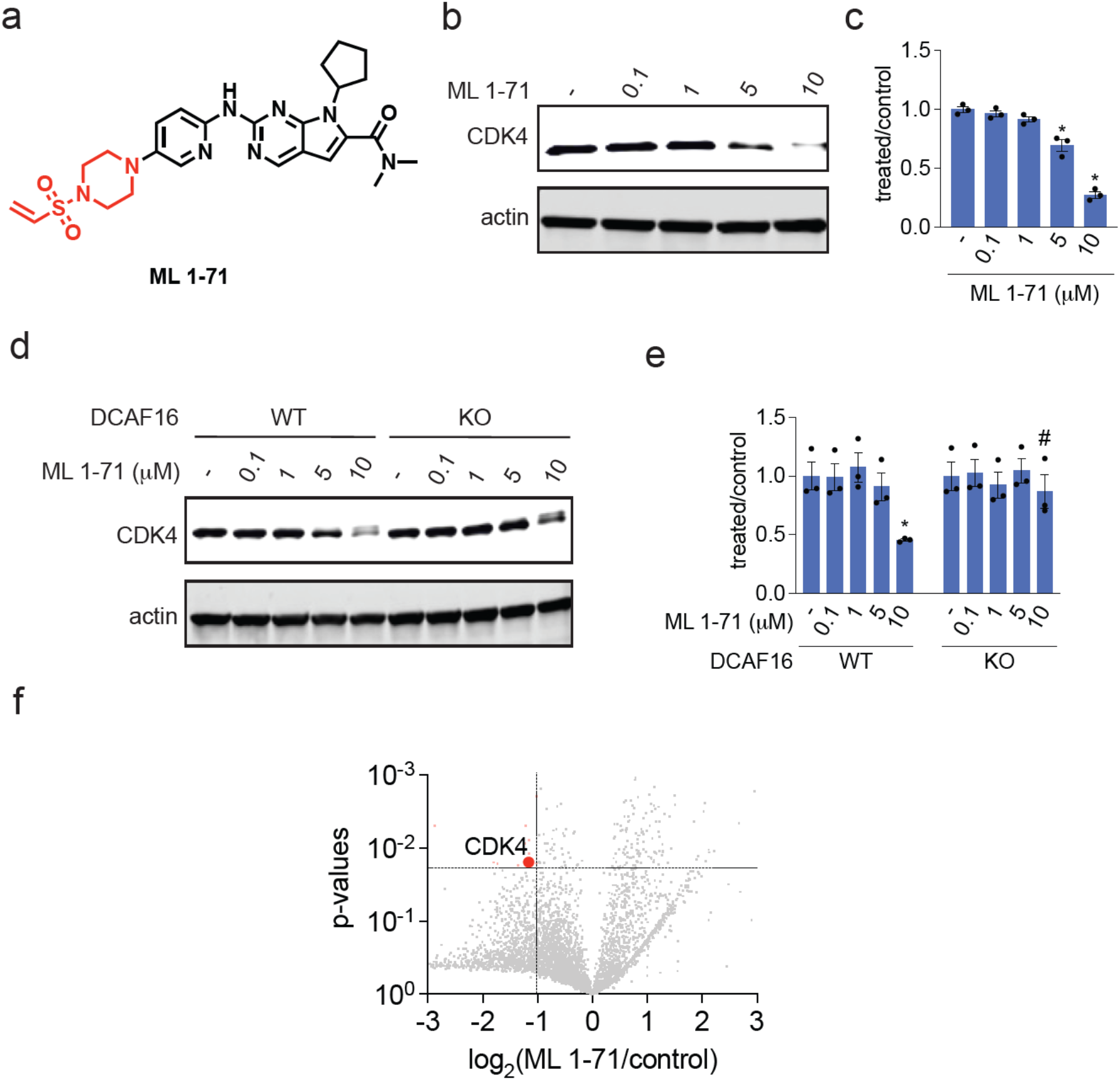
Characterization of CDK4 Monovalent Degrader. **(a)** Structure of ML 1-71, a CDK4 inhibitor ribociclib bearing a vinylsulfonyl piperazine handle highlighted in red. **(b, c)** CDK4 degradation in C33A cervical cancer cells. C33A cells were treated with DMSO vehicle or ML 1-71 for 24 h and CDK4 and actin loading control levels were assessed by Western blotting and quantified in **(c). (d, e)** CDK4 degradation in DCAF16 WT and KO cells. DCAF16 WT and KO HEK293 cells were treated with DMSO vehicle or ML 1-71 for 24 h and CDK4 and actin loading control levels were assessed by Western blotting and quantified in **(e). (f)** TMT-based quantitative proteomic profiling of ML 1-71 in C33A cells. C33A cells were treated with DMSO vehicle or ML 1-71 (10 µM) for 24 h. Proteins that were reduced in levels by >2-fold with p<0.05 are designated in red with CDK4 labeled. Data are from n=3 biologically independent replicates per group. Blots in **(b, d)** are representative of n=3 biologically independent replicates per group. Bar graph in **(e)** shows individual replicate values and average ± sem. Significance is expressed as ^*^p<0.05 compared to vehicle-treated controls and #p<0.05 compared to ML 1-71 treated DCAF16 WT cells.

To further explore the substrate scope of our covalent degradative handle, we next generated an androgen receptor (AR) monovalent degrader consisting of the AR-targeting ligand from the AR PROTAC ARV-110 and the vinylsulfonyl piperazine handle—ML 2-9 **(Figure 6a)** ^30^. ML 2-9 significantly degraded AR in LNCaP prostate cancer cells **(Figure 6b-6c)**. Interestingly, a hook effect was observed with this degrader. Proteomic data showed selective AR degradation with only 7 other proteins reduced in levels **(Figure 6d; Table S6)**. We also generated a vinylsulfonyl derivative of the BTK inhibitor ibrutinib, TH 1-9, and showed that this compound also degrades BTK in dose-dependent manner in MINO lymphoma cancer cells **(Figure 6e-6g)**. Proteomic analyses revealed a higher number of proteins beyond BTK that were lowered in levels compared to the other degraders **(Figure 6h)**. This may be due to other off-targets of ibrutinib that are also being degraded or downstream transcriptional changes resulting from BTK inhibition and degradation. We also made a vinylsulfonyl piperazine bearing derivative of the BCR-ABL and c-ABL kinase inhibitor dasatinib, ML 2-5, and demonstrated the degradation of both the fusion oncogene and the parent kinase in K562 leukemia cancer cells **(Figure S3a-S3c)**. This degradative covalent handle thus enabled the degradation of not only BRD4, but also several other proteins, including CDK4, SMARCA2/4, AR, BTK, and BCR-ABL/c-ABL.

**Figure 6.**
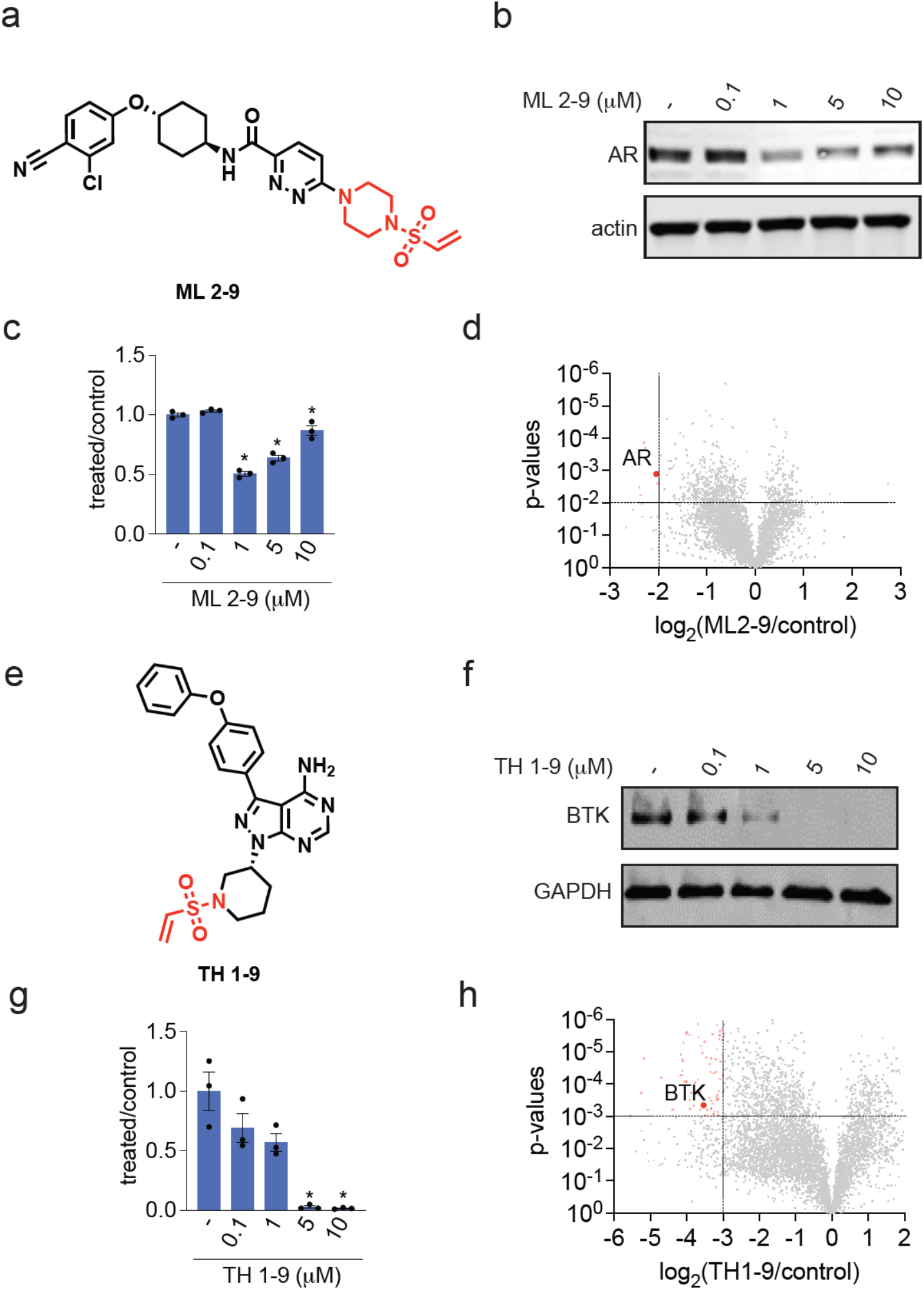
Characterization of AR and BTK Monovalent Degraders. **(a)** Structure of AR monovalent degrader ML 2-9 with AR-targeting ligand derived from the ARV-110 PROTAC bearing the covalent vinylsulfonyl piperazine handle highlighted in red. **(b, c)** AR degradation in LNCaP prostate cancer cells. LNCaP cells were treated with DMSO vehicle or ML 2-9 for 24 h and AR and actin loading control levels were assessed by Western blotting and quantified in **(c). (d)** TMT-based quantitative proteomic profiling of ML 2-9 in LNCaP cells. LNCaP cells were treated with DMSO vehicle or ML 2-9 (1 µM) for 24 h. Proteins that were reduced in levels by >4-fold with p<0.01 are designated in red with AR labeled. Data are from n=3 biologically independent replicates per group. **(e)** Structure of BTK monovalent degrader TH 1-9 with BTK inhibitor derived from ibrutinib bearing the covalent vinylsulfonyl piperazine handle highlighted in red. **(f, g)** BTK degradation in MINO lymphoma cancer cells. MINO cells were treated with DMSO vehicle or TH 1-9 for 24 h and BTK and GAPDH loading control levels were assessed by Western blotting and quantified in **(g). (h)** TMT-based quantitative proteomic profiling of TH 1-9 in MINO cells. MINO cells were treated with DMSO vehicle or TH 1-9 (5 µM) for 24 h. Proteins that were reduced in levels by >8-fold with p<0.001 are designated in red with BTK labeled. Data are from n=3 biologically independent replicates per group. Blots in **(b, f)** are representative of n=3 biologically independent replicates per group. Bar graphs in **(c, g)** show individual replicate values and average ± sem. Significance is expressed as ^*^p<0.05 compared to vehicle-treated controls.

## Discussion

Overall, we have identified a covalent DCAF16 handle that can be transplanted across a diverse range of protein targeting ligands, without the need for a linker, to enable the degradation of several different protein classes. We note that this is a proof-of-concept study further demonstrating the possibility of covalent chemical handle and E3 ligase pairs that can potentially be exploited towards rationally designing monovalent or molecular glue degraders. However, the potency, selectivity, and pharmacokinetic properties of our covalent handle would need to be substantially improved for future translational efforts. Our study, coupled with several other prior studies, also demonstrates the unique susceptibility of DCAF16 to covalent targeting for monovalent or heterobifunctional targeted protein degradation applications ^13,26,28^. This study also points to the potential permissiveness of DCAF16 in enabling the degradation of many different neo-substrate proteins, compared to other E3 ligases such as KEAP1 that may be much more restrictive in its substrate scope ^31^. However, we also demonstrate that not every electrophilic monovalent degrader acts through DCAF16. We show that our previously discovered covalent BRD4 degrader bearing a “fumarate” handle acts through RNF126, and not through DCAF16. Our results are analogous to recent findings with covalent heterobifunctional BRD4 degraders bearing two different electrophilic handles that act through DCAF16 and DCAF11 ^18^. Our study also suggests that pre-existing weak interactions between the E3 ligase and protein target may not be necessary to develop monovalent degraders. Overall, our study underscores the utility of covalent chemoproteomic approaches in identifying chemical starting points to expand the scope of targeted protein degradation applications.

## Acknowledgement

We thank the members of the Nomura Research Group and Novartis BioMedical Research for critical reading of the manuscript. This work was supported by Novartis BioMedical Research and the Novartis-Berkeley Translational Chemical Biology Institute (NB-TCBI) for all listed authors. This work was also supported by the Nomura Research Group and the Mark Foundation for Cancer Research ASPIRE Award. This work was also supported by grants from the National Institutes of Health (R01CA240981 and R35CA263814) and the National Science Foundation Molecular Foundations for Biotechnology Award (2127788). We also thank H. Celik, A. Lund, and UC Berkeley’s NMR facility in the College of Chemistry (CoC-NMR) for spectroscopic assistance. Instruments in the College of Chemistry NMR facility are supported in part by NIH S10OD024998.

## Author Contributions

ML, TDC, DKN conceived of the project idea and wrote the paper. ML, TDC, ET, LO, EL, DKN performed experiments, analyzed and interpreted data, and provided intellectual contributions. ACK generated the RNF126 knockout cell line.

## Methods

### Cell Culture

HEK293T and HEK293 cells were obtained from the UC Berkeley Cell Culture Facility and were cultured in Dulbecco’s Modified Eagle Medium (DMEM) containing 10% (v/v) fetal bovine serum (FBS) and maintained at 37 °C with 5% CO_2_. C33A cells were purchased from the American Type Culture Collection (ATCC) and were cultured in DMEM containing 10% (v/v) FBS and maintained at 37 °C with 5% CO_2_. K562 cells were obtained from the UC Berkeley Cell Culture Facility and were cultured in Iscove’s Modified Dulbecco’s Medium (IMDM) containing 10% (v/v) FBS and maintained at 37 °C with 5% CO_2_. MV-4-11 cells were obtained from the ATCC and were cultured in IMDM containing 10% (v/v) FBS and maintained at 37 °C with 5% CO_2_. Mino cells were obtained from the ATCC and were cultured in RPMI-1640 Medium containing 10% (v/v) FBS and maintained at 37 °C with 5% CO_2_. LNCaP cells were obtained from the UC Berkeley Cell Culture Facility and were cultured in DMEM containing 10% (v/v) FBS and maintained at 37 °C with 5% CO_2_. HEK293 DCAF16 knockout cells were purchased from Ubigene Biosciences and were cultured in DMEM containing 10% (v/v) FBS and maintained at 37 °C with 5% CO_2_. Unless otherwise specified, all cell culture materials were purchased from Gibco. It is not known whether HEK293T cells are from male or female origin.

### Western Blotting

Cells were washed twice with cold PBS, scraped, and pelleted by centrifugation (1,200 g, 5 min, 4 °C). Pellets were resuspended in PBS, lysed by sonication or RIPA lysis buffer (Thermo Scientific), clarified by centrifugation (12,000 g, 10 min, 4 °C), and lysate was transferred to new low-adhesion microcentrifuge tubes. Proteome concentrations were determined using the BCA assay and lysate was diluted to appropriate working concentrations. Proteins were resolved by SDS/PAGE and transferred to nitrocellulose membranes using the Trans-Blot Turbo transfer system (Bio-Rad). Membranes were blocked with 5% BSA in Tris-buffered saline containing Tween 20 (TBS-T) solution for 1 hr at RT, washed in TBS-T, and probed with primary antibody diluted in recommended diluent per manufacturer overnight at 4 °C. After 3 washes with TBS-T, the membranes were incubated in the dark with IR680- or IR800-conjugated secondary antibodies at 1:10,000 dilution in 5 % BSA in TBS-T at RT for 1 h. After 3 additional washes with TBST, blots were visualized using an Odyssey Li-Cor fluorescent scanner. The membranes were stripped using ReBlot Plus Strong Antibody Stripping Solution (EMD Millipore, 2504) when additional primary antibody incubations were performed. Antibodies used in this study were BRD4 (Abcam ab128874), CDK4 (Abcam ab108357), GAPDH (Cell Signaling Technology 14C10), Beta Actin (Cell Signaling Technology 13E5), c-Abl (Santa Cruz Biotechnology sc-23), SMARCA2 (Abcam ab240648), BRG1 (SMARCA4) (Cell Signaling Technology D1Q7F), BTK (Cell Signaling Technology D3H5), Androgen Receptor (Cell Signaling Technology D6F11).

### Bortezomib or MLN4924 Rescue Studies

2E6 of HEK293T cells per 3 mL of media were plated in 6-cm plates and left overnight to adhere. Cells were pretreated for 1 h with either Bortezomib (Cayman, C835F70) or MLN4924 (Tocris Bioscience, 649910) at a final concentration of 1 μM. Cells were then treated with ML1-50 until desired time point. Cells from both the supernatant and on the plate were harvested and assessed via western blot.

### Cell Viability Assay

HEK293T cells were seeded at a density of 20,000/well (100 µL) in 96-well white plates overnight. Cells were then treated with DMSO vehicle control or ML1-50 and incubated at 37 °C for 24 h. Cell viability assay was performed using CellTiter-Glo® 2.0 reagent (Promega, G9241) according to manufacturer’s protocol. Luminescent signals were measured using the Tecan Spark Plate reader (30086376).

### Isotopic desthiobiotin (isoDTB)-ABPP Cysteine Chemoproteomic Profiling of ML1-50

HEK293T cells were treated with either ML1-50 (10 µM) or DMSO for 2 h before cell collection and lysis. The proteome concentrations were determined using BCA assay and adjusted to 2 mg/mL. For each biological replicate, 2 aliquots of 1 mL of 2 mg/mL were used (i.e. 4 mg per condition). Each aliquot was treated with 20 µL of IA-alkyne (26.6 mg/mL in DMSO, 200 µM final concentration) for 1 h at RT. Two master mixes of the click reagents were prepared in the meanwhile, each containing 510 µL TBTA (0.9 mg/mL in 4:1 tBuOH/DMSO), 165 µL CuSO4 (12.5 mg/mL in H_2_O), 165 µL TCEP (14.0 mg/mL in H_2_O) and 160 µL of either heavy or light isoDTB tags (4 mg in DMSO, Click Chemistry Tools, 1565). The samples were then treated with 120 µL of the heavy (DMSO treated) or light (compound treated) master mix for 1 h at RT. After incubation, one light and one heavy labeled samples were combined and acetone-precipitated overnight at -20 °C. The samples were then centrifuged at 3,500 rpm for 10 min, acetone was removed, and the protein pellets resuspended in cold MeOH by sonication. The samples were centrifuged at 3,500 rpm for 10 min and MeOH was removed (repeated 3× in total). The pellets were dissolved in 600 µL urea (8 M in 0.1 M TEAB) by sonication and the urea concentration was then adjusted to 2 M by adding 1800 µL of TEAB (0.1 M). Two tubes containing solubilized proteins were combined, further diluted with 2400 µL 0.2% NP40 in PBS, and bound to high-capacity streptavidin agarose beads (200 µL/sample, ThermoFisher, 20357) for 1 h at RT with mixing. The beads were then centrifuged for 1 min at 1,000 g, the supernatant was removed, and the beads were washed 3 times with 0.1% NP40 in PBS, 3 times with PBS and 3 times with H_2_O. The samples were then resuspended in 8 M urea (600 µL in 0.1 M TEAB) and treated with DTT (30 µL, 31 mg/mL in H_2_O) for 45 min at 37 °C. They were then reacted with iodoacetamide (30 µL, 74 mg/mL in H_2_O) for 30 min at RT, followed by DTT (30 µL, 31 mg/mL in H_2_O) for 30 min at RT. The samples were diluted with 1800 µL TEAB (0.1 M), centrifuged for 1 min at 1,000 g, and the supernatant was removed. The beads were resuspended in 400 µL urea (2M in 0.1 M TEAB), and trypsin (8 µL, 0.5 mg/mL) was added and incubated for 20 h at 37 °C. The samples were then diluted with 800 µL 0.1% NP40 in PBS and the beads were washed 3 times with 0.1% NP40 in PBS, 3 times with PBS, and 3 times with H_2_O. Peptides were then eluted with 0.1% formic acid in 50% acetonitrile (3 × 400 µL). The samples were then dried using a vacuum concentrator at 30 °C, resuspended in 300 µL 0.1% TFA in H_2_O, and fractionated using high pH reversed-phase peptide fractionation kits (ThermoFisher, 84868) according to the manufacturer’s protocol.

### IsoDTB-ABPP Mass Spectrometry Analysis

Mass spectrometry analysis was performed on an Orbitrap Eclipse Tribrid Mass Spectrometer with a High Field Asymmetric Waveform Ion Mobility (FAIMS Pro) Interface (Thermo Scientific) with an UltiMate 3000 Nano Flow Rapid Separation LCnano System (Thermo Scientific). Off-line fractionated samples (5 μl aliquot of 15 μl sample) were injected via an autosampler (Thermo Scientific) onto a 5 μl sample loop which was subsequently eluted onto an Acclaim PepMap 100 C18 HPLC column (75 μm x 50 cm, nanoViper). Peptides were separated at a flow rate of 0.3 μl/min using the following gradient: 2 % buffer B (100 % acetonitrile with 0.1 % formic acid) in buffer A (95:5 water:acetonitrile, 0.1 % formic acid) for 5 min, followed by a gradient from 2 to 40 % buffer B from 5 to 159 min, 40 to 95 % buffer B from 159 to 160 minutes, holding at 95 % B from 160-179 min, 95 % to 2 % buffer B from 179 to 180 min, and then 2 % buffer B from 180 to 200 min. Voltage applied to the nano-LC electrospray ionization source was 2.1 kV. Data was acquired through an MS1 master scan (Orbitrap analysis, resolution 120,000, 400-1800 m/z, RF lens 30 %, heated capillary temperature 250 °C) with dynamic exclusion enabled (repeat count 1, duration 60 s). Data-dependent data acquisition comprised a full MS1 scan followed by sequential MS2 scans based on 2 s cycle times. FAIMS compensation voltages (CV) of -35, -45, and -55 were applied. MS2 analysis consisted of: quadrupole isolation window of 0.7 m/z of precursor ion followed by higher energy collision dissociation (HCD) energy of 38 % with an orbitrap resolution of 50,000.

Data was extracted in the form of MS1 and MS2 files using Raw Converter (Scripps Research Institute) and searched against the Uniprot human database using ProLuCID search methodology in IP2 v.3-v.5 (Integrated Proteomics Applications, Inc.) ^32^. Cysteine residues were searched with a static modification for carboxyaminomethylation (+57.02146) and up to two differential modifications for methionine oxidation and either the light or heavy isoDTB tags (+561.33872 or +567.34621, respectively). Peptides were required to be fully tryptic peptides. ProLuCID data were filtered through DTASelect to achieve a peptide false-positive rate below 5%. Only those probe-modified peptides that were evident across two out of three biological replicates were interpreted for their isotopic light to heavy ratios. Light versus heavy isotopic probe-modified peptide ratios are calculated by taking the mean of the ratios of each replicate paired light versus heavy precursor abundance for all peptide-spectral matches associated with a peptide. The paired abundances were also used to calculate a paired sample t-test P value in an effort to estimate constancy in paired abundances and significance in change between treatment and control. P values were corrected using the Benjamini–Hochberg method.

### Gel-Based ABPP

Recombinant DCAF16 (MyBioSource.com, MBS1375983) (0.1μg/sample) was pre-treated with either DMSO vehicle or covalent ligand at 37 °C for 30 min in 25 μL of PBS, and subsequently treated with of IA-Rhodamine (concentrations designated in figure legends) (Setareh Biotech) at room temperature for 1 h in the dark. The reaction was stopped by addition of 4×reducing Laemmli SDS sample loading buffer (Alfa Aesar). After boiling at 95 °C for 5 min, the samples were separated on precast 4−20% Criterion TGX gels (Bio-Rad). Probe-labeled proteins were analyzed by in-gel fluorescence using a ChemiDoc MP (Bio-Rad). Imaged gels were stained using Pierce™ Silver Stain Kit (Thermo Scientific™, 24612) following manufacturer’s instructions.

### ML1-50-Competed Targets from ML2-33 Probe Pulldown Proteomics

HEK293T cells were harvested, lysed, and the proteome concentration was adjusted to 5 mg/mL in 500 μL of PBS using the BCA assay. HEK293T cell lysate were pre-treated with DMSO vehicle or ML1-50 (200 μM) for 1 h at room temperature prior to ML2-33 probe labeling (20 μM) at room temperature for 1 h. To each tube containing cell lysate, the following reagents were added: 10 μL of 10 mM biotin picolyl azide (Sigma Aldrich, 900912) in DMSO, 10 μL of 50 mM TCEP in H_2_O, 10 μL of 50 mM CuSO_4_ in H_2_O, and 30 μL of TBTA ligand (1.7 mM in 1:4 DMSO/tBuOH, Cayman Chemical, 18816). The reaction mixture was incubated at room temperature for 60 minutes, and the reaction was quenched by protein precipitation. Precipitated pellets were washed using 500 μL of MeOH and centrifuged again to yield white pellets. Samples were resuspended in 1.2% SDS-PBS (1 mL), completely dissolved, and heated to 90 °C for 5 minutes. The soluble proteome was then diluted with 5 mL of PBS and further incubated with high-capacity streptavidin-agarose beads (100 μL/sample, ThermoFisher Scientific, 20357). Beads and lysates were incubated overnight at 4 °C with rotation. On the following day, beads were suspended and washed three times with 0.1% SDS-PBS, PBS, and H_2_O. Washed beads were resuspended in 6 M Urea/PBS (500 μL), and the samples were further treated with DTT and iodoacetamide. After removing the supernatant, beads were resuspended in 100 μL of 50 mM TEAB and enzymatically digested overnight using sequencing-grade trypsin (Promega, V5111). Digested peptides were eluted through centrifugation and labeled using commercially available TMTsixplex tags (ThermoFisher, P/N 90061). After labeling, 35 μg of each labeled sample was combined and dried using a vacufuge. Dried samples were redissolved with 300 μL of 0.1% TFA in H_2_O and further fractionated using high-pH reversed-phase peptide fractionation kits (ThermoFisher, P/N 84868) following the manufacturer’s protocol. Dried fractions were then resuspended in 25 μL of 0.1 % Formic acid/H_2_O (w/v) to be analyzed by LC-MS/MS.

### Quantitative TMT Proteomics Analysis

Cells were treated with either DMSO vehicle or compound (ML1-50 (1 µM, 24 h), ML1-71 (10µM, 16 h), ML1-96 (10 µM, 16 h), ML2-5 (10 µM, 16 h), TH1-9 (5 µM, 16 h), ML2-9 (1 µM, 24 h)) and lysate was prepared as described above. Briefly, 25-100 μg protein from each sample was reduced, alkylated and tryptically digested overnight. Individual samples were then labeled with isobaric tags using commercially available TMTsixplex (Thermo Fisher Scientific, P/N 90061) kits, in accordance with the manufacturer’s protocols. Tagged samples (20 μg per sample) were combined, dried using a vacuum concentrator at 30 °C, resuspended with 300 μL 0.1% TFA in H_2_O, and fractionated using high pH reversed-phase peptide fractionation kits (Thermo Fisher Scientific, P/N 84868) according to the manufacturer’s protocol. Fractions were dried using a vacuum concentrator at 30 °C, resuspended with 50 μL 0.1% FA in H_2_O, and analyzed by LC-MS/MS as described below.

Quantitative TMT-based proteomic analysis was performed as previously described using a Thermo Eclipse with FAIMS LC-MS/MS ^5^. Acquired MS data was processed using ProLuCID search methodology in IP2 v.3-v.5 (Integrated Proteomics Applications, Inc.) ^32^. Trypsin cleavage specificity (cleavage at K, R except if followed by P) allowed for up to 2 missed cleavages. Carbamidomethylation of cysteine was set as a fixed modification, methionine oxidation, and TMT-modification of N-termini and lysine residues were set as variable modifications. Reporter ion ratio calculations were performed using summed abundances with the most confident centroid selected from the 20 ppm window. Only peptide-to-spectrum matches that are unique assignments to a given identified protein within the total dataset are considered for protein quantitation. High confidence protein identifications were reported with a <1% false discovery rate (FDR) cut-off. Differential abundance significance was estimated using ANOVA with Benjamini-Hochberg correction to determine p-values.

### Knock Out Cell Line Generation

To generate a RNF126 knock-out pool in HEK293T, we introduced Cas9 ribonucleoproteins (RNPs) complexed with a custom Alt-R sgRNA synthesized by IDT targeting exon 2 of the RNF126 genomic locus (guide sequence ATGCGAGTCTGGTTTTATCG). spCas9 and sgRNA were introduced into cells by nucleofection. Briefly, 1.6 µL of 62.5 µM Cas9 (IDT, #1081058), 2.88 µL of 50 µM sgRNA (Alt-R from IDT), and 0.52 µL of 1X phosphate-buffered saline were mixed and the RNPs were incubated at room temperature for 30 minutes. Subsequently, the RNPs were added to 200,000 HEK293T cells resuspended in 16.4 µL Nucleofector solution SF plus 3.6 µL of supplement. To this suspension, 1.2 µL of 100 µM Alt-R electroporation enhancer (IDT, #1075916) and 4.32 µL H_2_O were added for a final volume of 30 µL. This nucleofection mix was electroporated using a 4D Nucleofector X Unit with program DG-130 in a nucleofector strip. After 10 minutes of recovery, nucleofected cells were grown in a 6-well dish for 7 days. This RNF126 knock-out pool was expanded and aliquoted for storage.

To isolate isogenic RNF126 knock-out clones, the knock-out pool was subjected to single cell sorting into 96-well plates using a WOLF microfluidic cell sorter (Nanocellect). Single cells were allowed to grow into colonies for two weeks, expanded further, and frozen into aliquots for storage. During cell expansion a sample of each clone was processed into lysate and RNF126 knock-out clones were identified by anti-RNF126 immunoblotting (ProteinTech, #66647-1-Ig). Clone 2B8 was designated as the RNF126 knock-out.

The DCAF16 knock-out cell line was purchased from Ubigene with guide sequences AGAGGGGGCCATTCAGGAAT TGG and TTCTGACAAGTGGTCAGGAG AGG (catalog number YKO-H721).

## Data Availability Statement

The datasets generated during and/or analyzed during the current study are available from the corresponding author on reasonable request.

## Code Availability Statement

Data processing and statistical analysis algorithms from our lab can be found on our lab’s Github site: https://github.com/NomuraRG, and we can make any further code from this study available at reasonable request.

## Safety Statement

No unexpected or unusually high safety hazards were encountered.

