## Supporting Information for "DCAF16-Based Covalent Handle for the Rational Design of Monovalent Degraders"

**Supporting Table Legends**

**Table S1. Quantitative proteomics of BRD4 degrader ML 1-50.** Tandem mass tagging (TMT)-based quantitative proteomic profiling of ML 1-50 in MDA-MB-231 cells. MDA-MB-231 cells were treated with DMSO vehicle or ML 1-50 (1μM) for 24 h. Data are from n=3 biologically independent replicates per group.

**Table S2. Chemoproteomic profiling of ML 1-50 competition against ML 2-33 probe.** ML 1-50-outcompeted targets enriched by ML 2-33. HEK293T cell lysate were pre-treated with DMSO vehicle or ML 1-50 (200 μM) 1 h prior to treatment with the ML 2-33 probe (20 μM). Probe-modified proteins were subjected to copper-catalyzed azide alkyne cycloaddition (CuAAC) with an azide-functionalized biotin enrichment handle. Probe-modified proteins were avidin-enriched, tryptically digested, and analyzed by TMT-based proteomics. Data are from n=3 biologically independent replicates per group.

**Table S3. isoDTB-ABPP analysis of ML 1-50.** isoDTB-ABPP profiling of ML 1-50. HEK293T cells were treated with DMSO vehicle or ML 1-50 (10 μM) for 2 h. Subsequent lysates were labeled with an alkyne-functionalized iodoacetamide probe (200 μM) for 1 h, followed by CuAAC-mediated attachment of an isotopically light (for control) or heavy (for treated) azide-functionalized desthiobiotin handle, after which probe-modified proteins were streptavidin-enriched, tryptically digested, eluted from beads, and analyzed by LC-MS/MS. Data are from n=3 biologically independent replicates per group.

**Table S4. Quantitative proteomics of DCAF16 WT and KO cells.** TMT-based quantitative proteomic analysis of DCAF16 wild-type (WT) versus knockout (KO) cells. Data are from n=3 biologically independent replicates per group.

**Table S5. Quantitative proteomics of CDK4 degrader ML 1-71.** TMT-based quantitative proteomic profiling of ML 1-71 in C33A cells. C33A cells were treated with DMSO vehicle or ML 1-71 (10 μM) for 24 h. Data are from n=3 biologically independent replicates per group.

**Table S6. Quantitative proteomics of AR degrader ML 2-9.** TMT-based quantitative proteomic profiling of ML 2-9 in LNCaP cells. LNCaP cells were treated with DMSO vehicle or ML 2-9 (1 μM) for 24 h. Data are from n=3 biologically independent replicates per group.

**Table S7. Quantitative proteomics of BTK degrader TH 1-9.** TMT-based quantitative proteomic profiling of TH 1-9 in MINO cells. MINO cells were treated with DMSO vehicle or TH 1-9 (5 μM) for 24 h. Data are from n=3 biologically independent replicates per group.

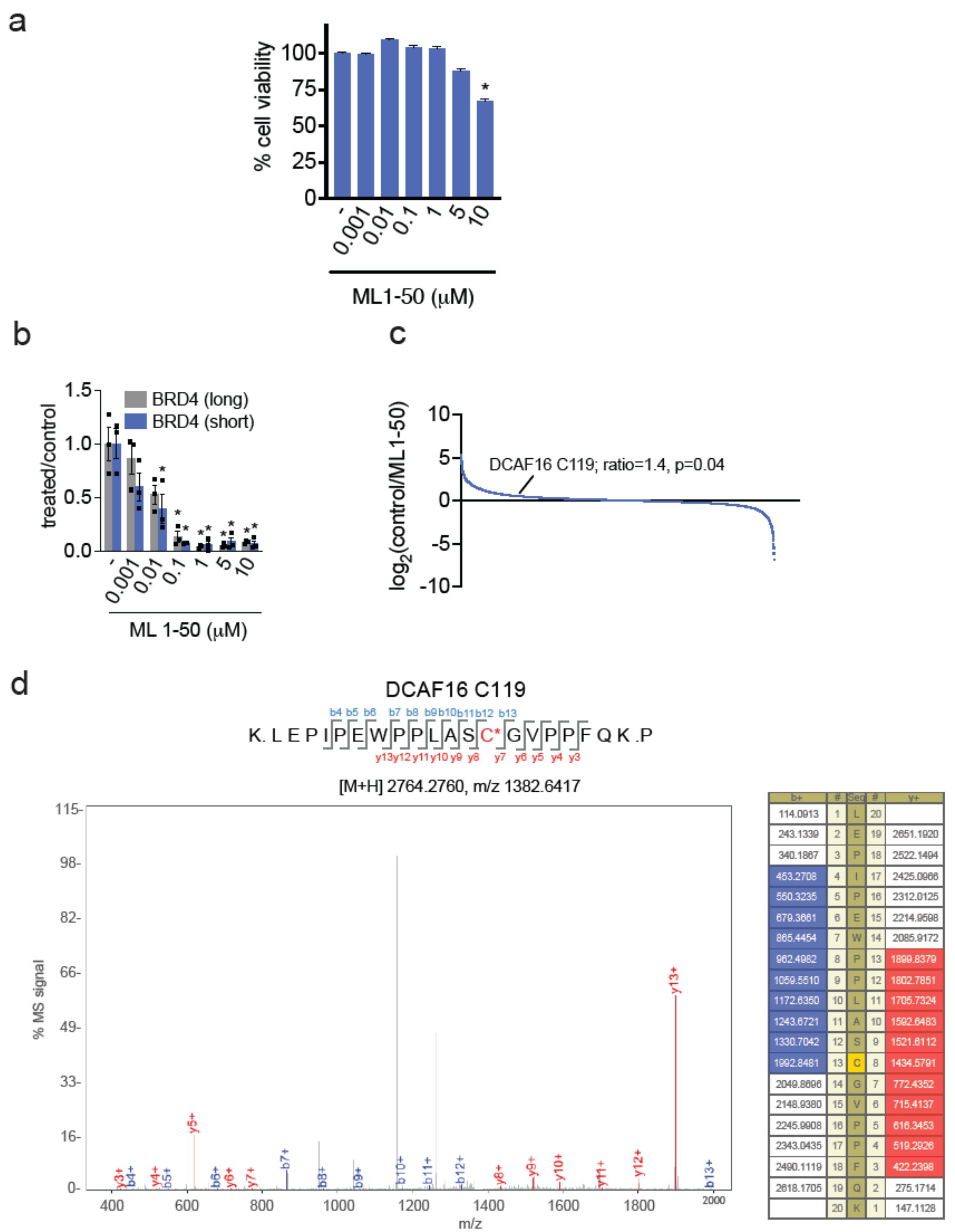


**Figure S1. Characterization of ML 1-50. (a)** Cell viability of HEK293T cells. HEK293T cells were treated with DMSO vehicle or ML 1-50 for 24 h and cell viability was assessed by CellTiter-Glo. **(b)** Quantification of BRD4 degradation in MDA-MB-231 cells from experiment described in **Figure 2h. (c)** IsoDTB-ABPP profiling of ML 1-50. HEK293T cells were treated with DMSO vehicle or ML 1-50 (10 μM) for 2h. Subsequent lysates were labeled with an alkyne-functionalized iodoacetamide probe (200 μM) for 1 h, followed by CuAAC-mediated attachment of an isotopically light (for control) or heavy (for treated) azide-functionalized desthiobiotin handle, after which probe-modified proteins were streptavidin-enriched, tryptically digested, eluted from beads, and analyzed by LC-MS/MS. The light versus heavy probe-modified peptide ratio for C119 of DCAF16 is noted with associated p-value. Each blue dot corresponds to the average light/heavy ratio of each probe-modified peptide. Data are from n=3 biologically independent replicates per group. **(d)** Site of modification analysis of ML 1-50 on pure DCAF16 protein. Pure DCAF16 protein was labeled with ML 1-50 (50 μM) for 1 h. Protein was tryptically digested and analyzed for the ML 1-50 adduct by LC-MS/MS.

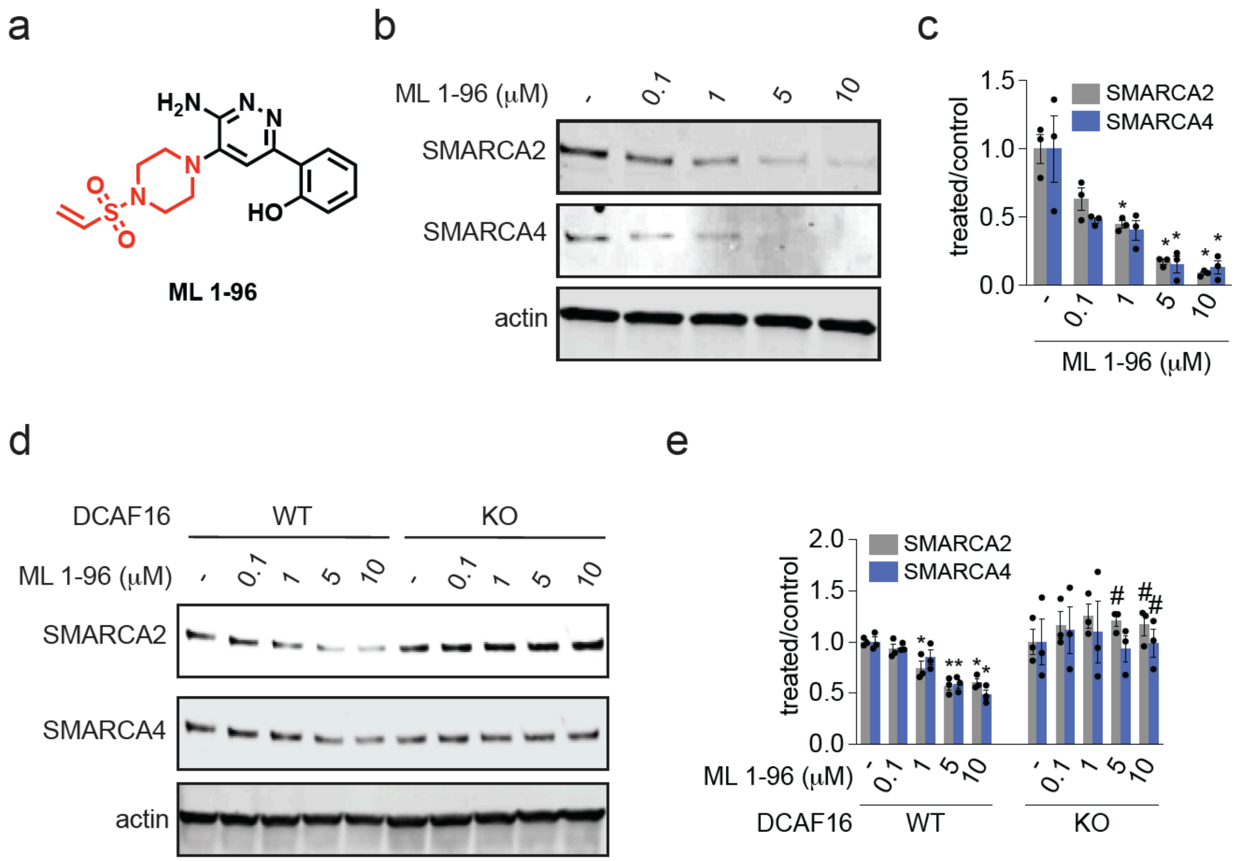


**Figure S2. Characterization of SMARCA2 Monovalent Degrader. (a)** Structure of ML 1-96 with the vinylsulfonyl piperazine covalent chemical handle in red. **(b,c)** SMARCA2 degradation in MV-4-11 leukemia cancer cells. MV-4-11 cells were treated with DMSO vehicle or ML 1-96 for 24 h and SMARCA2/4 and actin loading control levels were assessed by Western blotting and quantified in **(c)**. **(d, e)** BRD4 degradation in DCAF16 WT and KO cells. DCAF16 WT and KO HEK293 cells were treated with ML 1-96 for 24 h and SMARCA2, SMARCA4, and actin loading control levels were assessed by Western blotting and quantified in **(d).** Blots in **(b, d)** are representative of n=3 biologically independent replicates per group. Bar graphs in **(b, d)** show individual replicate values and average ± sem. Significance is expressed as *p<0.05 compared to vehicle-treated controls and #p<0.05 compared to ML 1-96 treated DCAF16 WT cells.

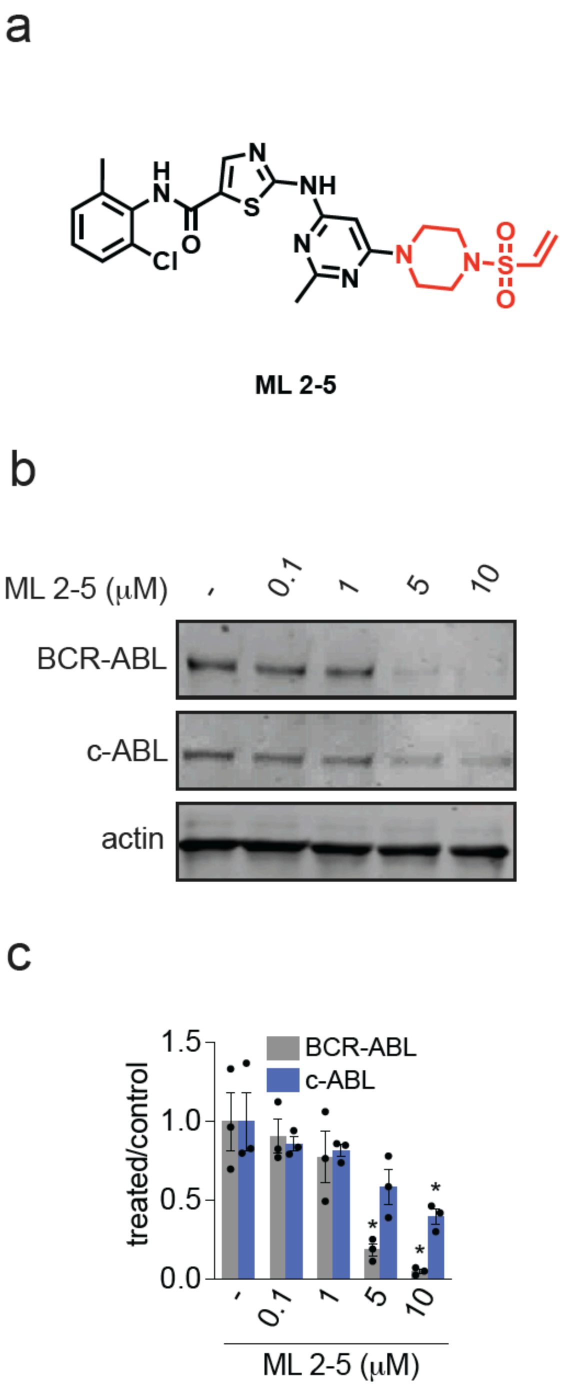


**Figure S3. Characterization of a BCR-ABL/c-ABL Monovalent Degrader. (a)** Structure of ML 2-5 with a dasatinib derivative bearing a vinylsulfonyl piperazine covalent chemical handle in red. **(b,c)** BCR-ABL and c-ABL degradation in K562 leukemia cancer cells. K562 cells were treated with DMSO vehicle or ML 2-5 for 24 h and SMARCA2/4 and actin loading control levels were assessed by Western blotting and quantified in **(c)**. Blot is representative is n=3 biologically independent replicates per group. Bar graph in **(c)** shows individual replicate values and average ± sem. Significance is expressed as *p<0.05 compared to vehicle-treated controls.

### Synthetic Methods and Characterization

All chemical reactions were carried out under a nitrogen atmosphere with dry solvents under anhydrous conditions, unless otherwise noted. Reagents were purchased at the highest commercial quality and used without further purification, unless otherwise stated. Room temperature is defined as between 21-25 °C. Reactions were stirred magnetically and monitored by thin layer chromatography (TLC) using TLC plates precoated with silica gel 60 F254 on aluminium (Merck KGaA). Detection was by UV (254 nm and 365 nm) or chemical stain (KMnO4, ninhydrin, iodine). Solvents were removed *in vacuo* using either a Buchi R-300 Rotavapor (equipped with an I-300 Pro Interface, B-300 Base Heating Bath, Welch 2037B-01 DryFast pump, and VWR AD15R-40-V11B Circulating Bath). Solvents for silica gel chromatography were used as supplied by Sigma-Aldrich. Automated flash chromatography was performed on a Biotage Isolera instrument, equipped with a UV detector. Chromatograms were recorded at 254 and 280 nm. If additional purification is needed, compound was further purified using Thermo Scientific's semi-prep reversed phase high-performance liquid chromatography equipped (RP-HPLC: Ultimate 3000 HPLC) equipped with C18 column (Luna® 10 µm c18(2), 100 Å, Serial #:5293-0084). Elution of the sample was monitored using DIONEX UltiMate 3000. Eluting buffer A: 100% distilled water + 0.1% trifluoracetic acid (TFA). Eluting buffer B: 95% acetonitrile + 5% distilled water + 0.1% TFA. High-resolution mass spectra (HRMS) were obtained using Q Exactive™ Plus Hybrid Quadrupole-Orbitrap™ Mass Spectrometer. $^{1}$H and $^{13}$C Nuclear Magnetic Resonance (NMR) spectra were recorded on BRUKER AV (600 MHz and 700 MHz), AVB (400 MHz), AVQ (400 MHz) and NEO (500 MHz) spectrometers. Measurements were carried out at ambient temperature. Chemical shifts (δ) are reported in ppm with the residual solvent signal as internal standard (chloroform at 7.26 and 77.2 ppm for $^{1}$H NMR and $^{13}$C NMR, respectively, methanol at 3.31 and 49.0, respectively and DMSO at 2.50 and 39.5, respectively). Multiplicity is reported as follows: singlet (s), doublet (d), doublet of doublet (dd) doublet of triplet (dt), triplet (t), triplet of doublet (td), quartet (q), and multiplet (m). Coupling constants (J) are reported in Hertz (Hz). $^{13}$C NMR spectra were recorded with broadband 1 H decoupling.

### General Procedures

#### Amide Couplings

#### General Procedure A

The corresponding carboxylic acid (1.0 equiv.) was added to a vessel and purged with $N_2$ for 5 minutes. The acid was dissolved in N,N-dimethylformamide (DMF) (0.1 M) and N,N-diisopropylethylamine (DIPEA) (3 equiv.) was added. A >50% wt. solution of propylphosphonic anhydride (T3P) in EtOAc (1.5 equiv.) was added dropwise, and the reaction mixture was stirred at ambient temperature for 30 minutes. The corresponding amine (1.2 equiv.) was dissolved in DMF (0.1 M) then added dropwise and the reaction mixture was stirred at ambient temperature overnight. The reaction was quenched with 5 times the reaction volume of 5% LiCl(aq) and extracted 3 times with ethyl acetate (EtOAc). The organic extracts were washed once with brine, dried over $Na_2SO_4$, vacuum filtered, and concentrated *in vacuo*. The resultant residue was purified by silica gel flash chromatography to afford the title compound.

#### General Procedure B

A mixture of the corresponding carboxylic acid (1.1 equiv.) and HATU (1.2 equiv.) was purged with $N_2$ for 5 minutes. The mixture was dissolved in DMF (0.1 M), DIPEA (3 equiv.) was added and the reaction mixture was allowed to stir at ambient temperature for 30 minutes. The corresponding amine (1 equiv.) was dissolved in DMF (0.1 M) then added dropwise and the reaction mixture was stirred at ambient temperature overnight. The reaction was quenched with 5 times the reaction volume of 5% LiCl(aq) and extracted 3 times with EtOAc. The organic extracts were dried over $Na_2SO_4$, vacuum filtered, and concentrated *in vacuo*. The resultant residue was purified by silica gel flash chromatography to afford the title compound.

#### Tert-butyloxycarbonyl Deprotection

#### General Procedure C

The corresponding tert-butyloxycarbonyl protected amine (1 equiv.) was dissolved in DCM (0.1 M). Trifluoroacetic acid (32 equiv.) was added, and the reaction mixture was stirred at ambient temperature for 30 minutes to overnight. The volatiles were removed *in vacuo* and the crude residue was used without further purification, unless otherwise noted.

#### Synthesis of Vinyl Sulfonamides

#### General Procedure D

The corresponding amine (1.0 equiv.) was dissolved in dichloromethane (DCM) (0.1 M) and triethylamine (3.0 equiv.) was added at 0 °C. 2-chloroethanesulfonyl chloride (0.8-1.2 equiv.) in DCM (0.1 M) was added dropwise and the resulting reaction mixture was stirred at ambient temperature overnight. The reaction was quenched with 5 times the reaction volume of water and extracted 3 times with DCM. The organic extracts were washed once with brine, dried over $Na_2SO_4$, vacuum filtered, and concentrated *in vacuo*. The resultant residue was purified by silica gel flash chromatography to afford the title compound.

**(S)-4-(2-(4-(4-chlorophenyl)-2,3,9-trimethyl-6H-thieno[3,2-f][1,2,4]triazolo[4,3-a][1,4]diazepin-6-yl)acetamido)benzenesulfonyl fluoride (ML1-10)**

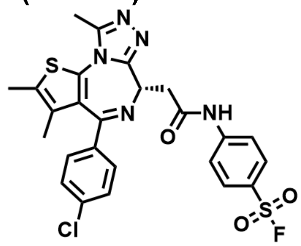

**General Procedure A** was followed with (S)-2-(4-(4-chlorophenyl)-2,3,9-trimethyl-6H-thieno [3,2-f][1,2,4]triazolo[4,3-a][1,4]diazepin-6-yl)acetic acid (JQ1-Acid) (50.0 mg, 0.12 mmol), T3P (0.1 mL, 0.19 mmol), DIPEA (0.07 mL, 0.37 mmol), and 4-aminobenzenesulfonyl fluoride (24.0 mg, 0.14 mmol). The crude residue was purified by silica gel chromatography (0-7% MeOH in DCM) to afford 16.6 mg (24%) of the title compound as a yellow-white powder.

**$^1$H NMR** (600 MHz, $CDCl_3$) δ 10.54 (s, 1H), 7.76 (d, *J* = 9.0 Hz, 2H), 7.72 (d, *J* = 9.0 Hz, 2H), 7.40 (d, *J* = 8.5 Hz, 2H), 7.31 (d, *J* = 8.8 Hz, 2H), 4.75 (dd, *J* = 9.7, 4.5 Hz, 1H), 3.99 (dd, *J* = 15.0, 9.7 Hz, 1H), 3.59 (dd, *J* = 15.0, 4.5 Hz, 1H), 2.72 (s, 3H), 2.44 (s, 2H), 1.71 (s, 2H).

**$^{13}$C NMR** (151 MHz, $CDCl_3$) δ 169.72, 164.43, 155.93, 150.35, 145.15, 137.14, 136.24, 131.89, 131.42, 131.03, 130.60, 129.90, 129.56, 128.78, 126.48, 126.32, 119.48, 54.24, 40.36, 14.42, 13.14, 11.86.

**HRMS (ESI)** m/z calcd for $C_{25}H_{21}ClFN_5NaO_3S_2^+$ $[M+Na]^+$: 580.0651; found: 580.0651

**(S)-tert-butyl (4-(2-(4-(4-chlorophenyl)-2,3,9-trimethyl-6H-thieno[3,2-f][1,2,4]triazolo[4,3-a] [1,4]diazepin-6-yl)acetamido)phenyl) carbonate (ML1-13)**

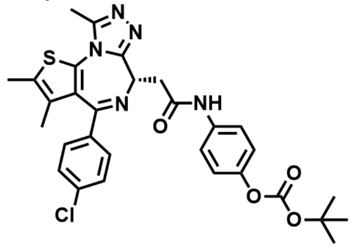

**General Procedure A** was followed with (S)-2-(4-(4-chlorophenyl)-2,3,9-trimethyl-6H-thieno [3,2-f][1,2,4]triazolo[4,3-a][1,4]diazepin-6-yl)acetic acid (JQ1-Acid) (30.0 mg, 0.07 mmol), T3P (0.07 mL, 0.11 mmol), DIPEA (0.04 mL, 0.22 mmol), and 4-aminophenyl tert-butyl carbonate (17.0 mg, 0.08 mmol). The crude residue was purified by silica gel chromatography (0-7% MeOH in DCM) to afford 30.0 mg (68%) of the title compound as a yellow-white powder.

**$^1$H NMR** (600 MHz, $CDCl_3$) δ 9.36 (s, 1H), 7.57 (d, *J* = 9.0 Hz, 2H), 7.39 (d, *J* = 8.3 Hz, 2H), 7.30 (d, *J* = 8.8 Hz, 2H), 7.05 (d, *J* = 9.0 Hz, 2H), 4.68 (dd, *J* = 8.1, 5.9 Hz, 1H), 3.81 (dd, *J* = 14.4, 8.2 Hz, 1H), 3.58 (dd, *J* = 14.3, 5.9 Hz, 1H), 2.67 (s, 3H), 2.39 (s, 3H), 1.67 (s, 3H), 1.53 (s, 9H).

**$^{13}$C NMR** (151 MHz, $CDCl_3$) δ 168.91, 164.05, 155.79, 151.92, 149.99, 147.08, 136.86, 136.51, 136.00, 132.08, 130.99, 130.96, 130.52, 129.88, 128.72, 121.46, 120.78, 83.35, 54.59, 40.36, 27.71, 14.38, 13.09, 11.82.

**HRMS (ESI)** m/z calcd for $C_{30}H_{30}ClN_5NaO_4S^+$ $[M+Na]^+$: 614.1599; found:614.1600

**(S)-2,4-dichloro-5-(2-(4-(4-chlorophenyl)-2,3,9-trimethyl-6H-thieno[3,2-f][1,2,4]triazolo[4,3-a][1,4]diazepin-6-yl)acetamido)phenyl ethyl carbonate (ML1-14)**

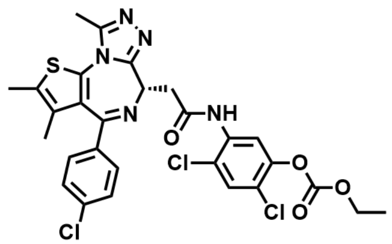

**General Procedure A** was followed with (S)-2-(4-(4-chlorophenyl)-2,3,9-trimethyl-6H-thieno [3,2-f][1,2,4]triazolo[4,3-a][1,4]diazepin-6-yl)acetic acid (JQ1-Acid) (30.0 mg, 0.07 mmol), T3P (0.07 mL, 0.11 mmol), DIPEA (0.04 mL, 0.22 mmol), and 5-amino-2,4-dichlorophenyl ethyl carbonate (20.6 mg, 0.08 mmol). The crude residue was purified by silica gel chromatography (0-8% MeOH in DCM) to afford 5.0 mg (10.6%) of the title compound as a yellow-white powder.

**$^1$H NMR** (600 MHz, $CDCl_3$) δ 8.98 (s, 1H), 8.50 (s, 1H), 7.44 (s, 1H), 7.43 (d, *J* = 8.5 Hz, 2H), 7.34 (d, *J* = 8.8 Hz, 2H), 4.60 (t, *J* = 6.5 Hz, 1H), 4.33 (q, *J* = 7.2 Hz, 2H), 3.77 (dd, *J* = 14.6, 6.9 Hz, 1H), 3.58 (dd, *J* = 14.5, 6.1 Hz, 1H), 2.68 (s, 3H), 2.41 (d, *J* = 0.8 Hz, 3H), 1.71 – 1.68 (m, 3H), 1.38 (t, *J* = 7.1 Hz, 3H).

**$^{13}$C NMR** (151 MHz, $CDCl_3$) δ 169.09, 164.38, 152.30, 150.09, 146.02, 137.08, 134.61, 132.26, 130.26, 129.94, 129.68, 128.76, 121.58, 120.49, 116.07, 65.52, 54.22, 40.80, 14.43, 14.16, 13.11, 11.84.

**HRMS (ESI)** m/z calcd for $C_{28}H_{24}Cl_3N_5NaO_4S^+$ $[M+Na]^+$: 654.0507; found: 654.0512

**(S)-2-(4-(4-chlorophenyl)-2,3,9-trimethyl-6H-thieno[3,2-f][1,2,4]triazolo[4,3-a][1,4]diazepin-6-yl)-N-(2,4-dioxo-1,4-dihydro-2H-benzo[d][1,3]oxazin-7-yl)acetamide (ML1-15)**

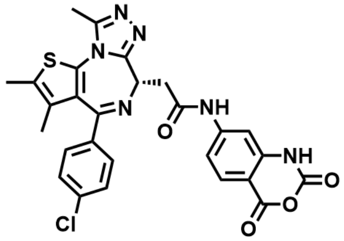

**General Procedure A** was followed with (S)-2-(4-(4-chlorophenyl)-2,3,9-trimethyl-6H-thieno [3,2-f][1,2,4]triazolo[4,3-a][1,4]diazepin-6-yl)acetic acid (JQ1-Acid) (30.0 mg, 0.07 mmol), T3P (0.07 mL, 0.11 mmol), DIPEA (0.04 mL, 0.22 mmol), and 7-amino-2H-benzo[d][1,3]oxazine-2,4 (1H)-dione (14.7 mg, 0.08 mmol). The crude residue was purified by silica gel chromatography (0-8% MeOH in DCM) to afford 8.6 mg (20.5%) of the title compound as a yellow-white powder.

**$^1$H NMR** (600 MHz, $CDCl_3$) δ 11.23 (s, 1H), 10.42 (s, 1H), 7.47 (s, 1H), 7.38 (d, *J* = 8.2 Hz, 2H), 7.21 (d, *J* = 8.2 Hz, 2H), 5.03 – 5.00 (m, 1H), 4.24 – 4.21 (m, 1H), 3.76 – 3.73 (m, 1H), 2.72 (s, 3H), 2.38 (s, 3H), 1.72 (s, 3H).

**$^{13}$C NMR** (151 MHz, $CDCl_3$) δ 164.37, 156.26, 155.10, 153.41, 150.58, 136.64, 136.56, 131.85, 131.03, 130.95, 130.43, 130.00, 129.78, 129.04, 128.53, 115.66, 54.18, 40.37, 14.38, 13.11, 11.93.

**HRMS (ESI)** m/z calcd for $C_{27}H_{21}ClN_6NaO_4S^+$ $[M+Na]^+$: 583.0926; found: 583.0934

**(S)-2-(4-(4-chlorophenyl)-2,3,9-trimethyl-6H-thieno[3,2-f][1,2,4]triazolo[4,3-a][1,4]diazepin-6-yl)-N-(2-methoxy-3,4-dioxocyclobut-1-en-1-yl)acetamide (ML1-16)**

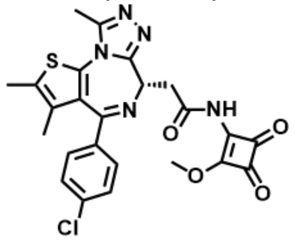

**General Procedure A** was followed with (S)-2-(4-(4-chlorophenyl)-2,3,9-trimethyl-6H-thieno [3,2-f][1,2,4]triazolo[4,3-a][1,4]diazepin-6-yl)acetic acid (JQ1-Acid) (30.0 mg, 0.07 mmol), T3P (0.07 mL, 0.11 mmol), DIPEA (0.04 mL, 0.22 mmol), and 3-amino-4-methoxycyclobut-3-ene- 1,2-dione (10.5 mg, 0.08 mmol). The crude residue was purified by silica gel chromatography (0-7% MeOH in DCM) to afford 8.0 mg (21.0%) of the title compound as a yellow-white powder.

**$^1$H NMR** (600 MHz, $CDCl_3$) δ 10.97 (s, 1H), 7.42 (d, *J* = 8.3 Hz, 2H), 7.35 (d, *J* = 8.8 Hz, 2H), 4.65 (t, *J* = 6.5 Hz, 1H), 4.50 (s, 3H), 3.81 (dd, *J* = 8.0, 6.5 Hz, 2H), 2.72 (s, 3H), 2.42 (s, 3H), 1.70 (s, 3H).

**$^{13}$C NMR** (151 MHz, $CDCl_3$) δ 168.01, 150.62, 150.36, 136.05, 135.32, 132.58, 132.13, 132.12, 131.30, 131.03, 129.94, 128.87, 61.21, 53.67, 39.03, 14.47, 13.13, 11.73.
**HRMS (ESI)** m/z calcd for $C_{24}H_{20}ClN_5NaO_4S^+$ $[M+Na]^+$: 532.0817; found: 532.0819

**(S)-2-(4-(4-chlorophenyl)-2,3,9-trimethyl-6H-thieno[3,2-f][1,2,4]triazolo[4,3-a][1,4]diazepin-6-yl)-N-(6-formylpyridin-3-yl)acetamide (ML1-17)**

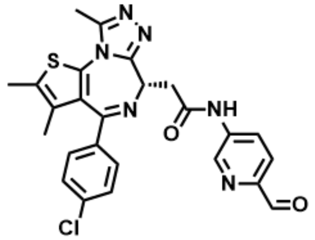

**General Procedure A** was followed with (S)-2-(4-(4-chlorophenyl)-2,3,9-trimethyl-6H-thieno [3,2-f][1,2,4]triazolo[4,3-a][1,4]diazepin-6-yl)acetic acid (JQ1-Acid) (30.0 mg, 0.07 mmol), T3P (0.07 mL, 0.11 mmol), DIPEA (0.04 mL, 0.22 mmol), and 5-aminopicolinaldehyde (10.5 mg, 0.08 mmol). The crude residue was purified by silica gel chromatography (0-7% MeOH in DCM) to afford 4.0 mg (10.5%) of the title compound as a yellow-white powder.
**$^{1}$H NMR** (600 MHz, $CDCl_3$) δ 10.45 (s, 1H), 9.91 (s, 1H), 8.75 (d, *J* = 2.4 Hz, 1H), 8.22 (dd, *J* = 8.5, 2.4 Hz, 1H), 7.78 (d, *J* = 8.5 Hz, 1H), 7.41 (d, *J* = 8.4 Hz, 2H), 7.32 (d, *J* = 8.7 Hz, 2H), 4.76 (dd, *J* = 9.2, 4.6 Hz, 1H), 3.97 (dd, *J* = 14.9, 9.3 Hz, 1H), 3.63 (dd, *J* = 14.9, 4.7 Hz, 1H), 2.72 (s, 3H), 2.44 (s, 3H), 1.71 (s, 3H).
**$^{13}$C NMR** (151 MHz, $CDCl_3$) δ 192.15, 169.82, 164.47, 155.82, 150.34, 147.97, 141.08, 139.10, 137.17, 136.22, 131.94, 131.43, 131.02, 130.56, 129.89, 128.81, 126.17, 122.50, 54.27, 40.19, 14.43, 13.16, 11.88.
**HRMS (ESI)** m/z calcd for $C_{25}H_{21}ClN_6NaO_2S^+$ $[M+Na]^+$: 527.1027; found: 527.1034

**(S)-2-(4-(4-chlorophenyl)-2,3,9-trimethyl-6H-thieno[3,2-f][1,2,4]triazolo[4,3-a][1,4]diazepin-6-yl)-N-(6-formylpyridin-2-yl)acetamide (ML1-25)**

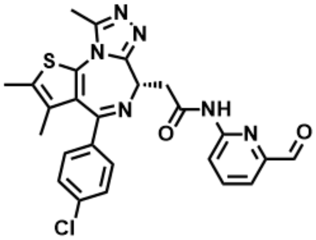

**General Procedure A** was followed with (S)-2-(4-(4-chlorophenyl)-2,3,9-trimethyl-6H-thieno [3,2-f][1,2,4]triazolo[4,3-a][1,4]diazepin-6-yl)acetic acid (JQ1-Acid) (25.0 mg, 0.06 mmol), T3P (0.06 mL, 0.09 mmol), DIPEA (0.03 mL, 0.19 mmol), and 6-aminopicolinaldehyde (9.1 mg, 0.07 mmol). The crude residue was purified by silica gel chromatography (0-7% MeOH in DCM) to afford 6.0 mg (19.0%) of the title compound as a yellow-white powder.
**$^{1}$H NMR** (600 MHz, $CDCl_3$) δ 9.96 (s, 1H), 9.42 (s, 1H), 8.46 (d, *J* = 8.3 Hz, 1H), 7.87 (t, *J* = 7.9 Hz, 1H), 7.69 (d, *J* = 7.4 Hz, 1H), 7.52 (d, *J* = 8.1 Hz, 2H), 7.36 (d, *J* = 8.7 Hz, 2H), 4.66 (t, *J* = 6.8 Hz, 1H), 3.70 (d, *J* = 6.9 Hz, 2H), 2.69 (s, 3H), 2.41 (s, 3H), 1.69 (s, 3H).
**$^{13}$C NMR** (151 MHz, $CDCl_3$) δ 192.68, 169.71, 164.55, 155.29, 151.79, 150.99, 150.14, 139.25, 137.16, 136.39, 132.32, 131.06, 130.98, 130.29, 129.98, 128.82, 118.66, 117.83, 54.13, 41.03, 40.35, 14.47, 13.12, 11.88.
**HRMS (ESI)** m/z calcd for $C_{25}H_{21}ClN_6NaO_2S^+$ $[M+Na]^+$: 527.1027; found: 527.1024

**tert-butyl (S)-4-(2-(4-(4-chlorophenyl)-2,3,9-trimethyl-6H-thieno[3,2-f][1,2,4]triazolo[4,3-a] [1,4]diazepin-6-yl)acetyl)piperazine-1-carboxylate (ML1-26)**

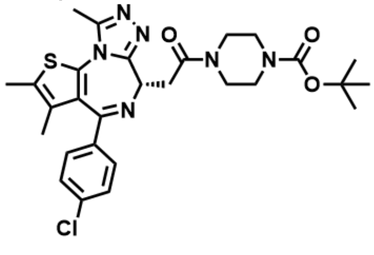

**General Procedure A** was followed with (S)-2-(4-(4-chlorophenyl)-2,3,9-trimethyl-6H-thieno [3,2-f][1,2,4]triazolo[4,3-a][1,4]diazepin-6-yl)acetic acid (JQ1-Acid) (322.0 mg, 0.80 mmol), T3P (0.53 mL, 0.88 mmol), DIPEA (0.56 mL, 3.21 mmol), and 1-boc-piperazine (179.5 mg, 0.96 mmol). The crude residue was purified by silica gel chromatography (0-5% MeOH in DCM) to afford 338.9 mg (74.1%) of the title compound as a yellow oil.
**$^1$H NMR** (600 MHz, $CDCl_3$) δ 7.41 (d, *J* = 8.6 Hz, 2H), 7.34 (d, *J* = 8.7 Hz, 2H), 4.84 – 4.79 (m, 1H), 3.84 (ddd, *J* = 13.3, 6.3, 3.5 Hz, 1H), 3.75 (dd, *J* = 15.9, 6.2 Hz, 2H), 3.72 – 3.65 (m, 1H), 3.58 (ddp, *J* = 15.4, 7.6, 4.0 Hz, 4H), 3.52 (s, 1H), 3.41 (ddd, *J* = 13.2, 7.5, 3.6 Hz, 1H), 2.68 (s, 3H), 2.41 (d, *J* = 0.9 Hz, 3H), 1.69 (s, 3H), 1.50 (s, 9H).
**$^{13}$C NMR** (151 MHz, $CDCl_3$) δ 169.22, 163.71, 155.82, 154.61, 149.83, 136.82, 136.67, 132.24, 130.91, 130.65, 130.52, 129.80, 128.69, 80.25, 54.47, 45.74, 41.70, 35.35, 28.41, 14.35, 13.06, 11.83.
**HRMS (ESI)** m/z calcd for $C_{28}H_{33}ClN_6NaO_3S^+$ $[M+Na]^+$: 591.1916; found: 691.1922

**(S)-4-(4-(2-(4-(4-chlorophenyl)-2,3,9-trimethyl-6H-thieno[3,2-f][1,2,4]triazolo[4,3-a][1,4]diazepin-6-yl)acetyl)piperazine-1-carbonyl)-2-hydroxybenzaldehyde (ML1-29)**

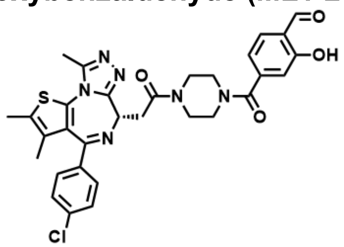

**General Procedure C** was followed with **ML1-26** (35.0 mg, 0.06 mmol), and TFA (0.15 mL, 1.97 mmol) for 2 hours. The crude material was used without further purification.
**General Procedure A** was followed with 4-formyl-3-hydroxybenzoic acid (12.8 mg, 0.08 mmol), T3P (0.06 mL, 0.10 mmol), DIPEA (0.06 mL, 0.32 mmol), and the amine from above (30.0 mg, 0.06 mmol). The crude residue was purified by silica gel chromatography (0-7% MeOH in DCM) to afford 10.0 mg (25.3%) of the title compound as a yellow-white powder.
**$^1$H NMR** (600 MHz, $CDCl_3$) δ 11.12 (s, 1H), 9.95 (s, 1H), 7.67 (d, *J* = 7.8 Hz, 1H), 7.39 (d, *J* = 8.2 Hz, 2H), 7.33 (d, *J* = 8.7 Hz, 2H), 7.06 (d, *J* = 6.4 Hz, 0H), 7.02 (s, 1H), 4.80 (t, *J* = 6.8 Hz, 1H), 4.02 – 3.97 (m, 2H), 3.91 – 3.84 (m, 1H), 3.81 – 3.78 (m, 2H), 3.63 – 3.56 (m, 2H), 3.51 – 3.45 (m, 1H), 3.44 – 3.39 (m, 2H), 2.67 (s, 3H), 2.40 (s, 3H), 1.67 (s, 1H).
**$^{13}$C NMR** (151 MHz, $CDCl_3$) δ 196.18, 168.70, 163.96, 161.66, 149.97, 143.44, 136.81, 136.70, 134.37, 132.20, 130.94, 130.45, 129.79, 128.77, 121.11, 118.20, 116.10, 54.36, 46.34, 41.96, 14.42, 13.14, 11.88.
**HRMS (ESI)** m/z calcd for $C_{31}H_{29}ClN_6NaO_4S^+$ $[M+Na]^+$: 639.1552; found: 639.1562

**(*S*)-5-(4-(2-(4-(4-chlorophenyl)-2,3,9-trimethyl-6*H*-thieno[3,2-*f*][1,2,4]triazolo[4,3-*a*][1,4]diazepin-6-yl)acetyl)piperazine-1-carbonyl)-1*H*-pyrrole-3-sulfonyl fluoride (ML1-30)**

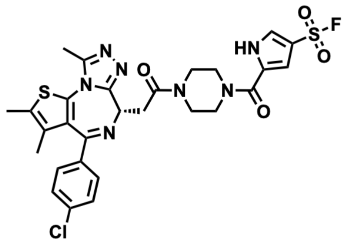

**General Procedure C** was followed with **ML1-26** (35.0 mg, 0.06 mmol), and TFA (0.15 mL, 1.97 mmol) for 2 hours. The crude material was used without further purification.
**General Procedure A** was followed with 4-(fluorosulfonyl)-1H-pyrrole-2-carboxylic acid (14.8 mg, 0.08 mmol), T3P (0.06 mL, 0.10 mmol), DIPEA (0.06 mL, 0.32 mmol), and the amine from above (30.0 mg, 0.06 mmol). The crude residue was purified by silica gel chromatography (0-7% MeOH in DCM) to afford 24.8 mg (60.2%) of the title compound as a yellow-white powder.
**$^1$H NMR** (600 MHz, $CDCl_3$) δ 11.71 (s, 1H), 7.66 (s, 1H), 7.40 (d, *J* = 8.3 Hz, 2H), 7.33 (d, *J* = 8.3 Hz, 2H), 6.93 (d, *J* = 1.5 Hz, 1H), 4.82 (t, *J* = 6.9 Hz, 1H), 4.10 – 4.01 (m, 2H), 3.96 (tt, *J* = 15.2, 6.8 Hz, 3H), 3.89 – 3.75 (m, 3H), 3.67 – 3.60 (m, 1H), 3.52 (dd, *J* = 16.3, 6.4 Hz, 1H), 2.67 (s, 3H), 2.40 (s, 3H), 1.68 (s, 3H).

**$^{13}$C NMR** (151 MHz, $CDCl_3$) δ 169.57, 164.00, 160.46, 155.72, 149.99, 136.82, 136.71, 132.16, 130.94, 130.90, 130.54, 129.81, 128.75, 127.05, 126.70, 116.16, 115.96, 111.68, 54.57, 45.44, 41.37, 35.36, 31.92, 22.68, 14.36, 13.09, 11.81.
**HRMS (ESI)** m/z calcd for $C_{28}H_{27}ClFN_7NaO_4S_2^+$ [M+Na]$^+$: 666.1131; found: 666.1127

**(*S*)-4-(4-(2-(4-(4-chlorophenyl)-2,3,9-trimethyl-6*H*-thieno[3,2-*f*][1,2,4]triazolo[4,3-*a*][1,4]diazepin-6-yl)acetyl)piperazine-1-carbonyl)benzenesulfonyl fluoride (ML1-31)**

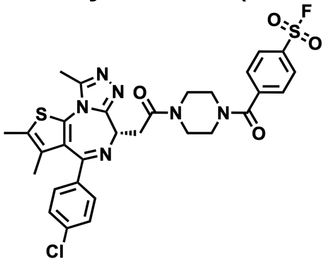

**General Procedure C** was followed with **ML1-26** (35.0 mg, 0.06 mmol), and TFA (0.15 mL, 1.97 mmol) for 2 hours. The crude material was used without further purification.
**General Procedure A** was followed with 4-(fluorosulfonyl)benzoic acid (15.7 mg, 0.08 mmol), T3P (0.06 mL, 0.10 mmol), DIPEA (0.06 mL, 0.32 mmol), and the amine from above (30.0 mg, 0.06 mmol). The crude residue was purified by silica gel chromatography (0-7% MeOH in DCM) to afford 22.7 mg (54.2%) of the title compound as a yellow-white powder.
**$^1$H NMR** (600 MHz, $CDCl_3$) δ 8.10 (d, *J* = 8.2 Hz, 2H), 7.68 (d, *J* = 8.0 Hz, 2H), 7.39 (d, *J* = 8.2 Hz, 2H), 7.33 (d, *J* = 8.6 Hz, 2H), 4.80 (dd, *J* = 7.4, 6.4 Hz, 1H), 4.09 – 4.00 (m, 3H), 3.88 – 3.81 (m, 2H), 3.74 – 3.30 (m, 6H), 2.67 (s, 3H), 2.40 (s, 3H), 1.67 (s, 3H).
**$^{13}$C NMR** (151 MHz, $CDCl_3$) δ 155.77, 150.10, 142.56, 136.96, 136.83, 134.54, 134.37, 132.31, 131.06, 131.00, 130.61, 129.91, 129.12, 128.88, 128.54, 128.46, 32.04, 30.44, 29.47, 22.81, 14.48, 13.21, 11.92.
**HRMS (ESI)** m/z calcd for $C_{30}H_{28}ClFN_6NaO_4S_2^+$ [M+Na]$^+$: 677.1178; found: 677.1175

**(*S*)-3-(4-(2-(4-(4-chlorophenyl)-2,3,9-trimethyl-6*H*-thieno[3,2-*f*][1,2,4]triazolo[4,3-*a*][1,4]diazepin-6-yl)acetyl)piperazine-1-carbonyl)benzenesulfonyl fluoride (ML1-32)**

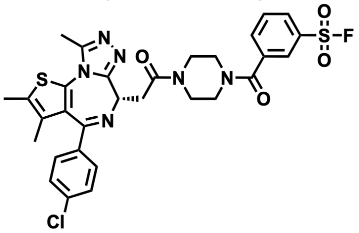

**General Procedure C** was followed with **ML1-26** (35.0 mg, 0.06 mmol), and TFA (0.15 mL, 1.97 mmol) for 2 hours. The crude material was used without further purification.
**General Procedure A** was followed with 3-(fluorosulfonyl)benzoic acid (15.7 mg, 0.08 mmol), T3P (0.06 mL, 0.10 mmol), DIPEA (0.06 mL, 0.32 mmol), and the amine from above (30.0 mg, 0.06 mmol). The crude residue was purified by silica gel chromatography (0-7% MeOH in DCM) to afford 10.0 mg (23.9%) of the title compound as a yellow-white powder.
**$^1$H NMR** (600 MHz, $CDCl_3$) δ 8.10 (dq, *J* = 3.3, 1.7 Hz, 2H), 7.85 (dt, *J* = 7.7, 1.4 Hz, 1H), 7.74 (t, *J* = 8.1 Hz, 1H), 7.40 (d, *J* = 8.3 Hz, 2H), 7.34 (d, *J* = 8.5 Hz, 2H), 4.82 (t, *J* = 6.8 Hz, 1H), 4.11 – 3.95 (m, 2H), 3.84 (dd, *J* = 15.9, 7.1 Hz, 2H), 3.78 – 3.30 (m, 5H), 2.69 (s, 3H), 2.40 (s, 3H), 1.68 (s, 3H).
**$^{13}$C NMR** (151 MHz, $CDCl_3$) δ 169.41, 167.64, 164.04, 155.64, 150.02, 137.17, 136.89, 136.64, 134.05, 132.04, 130.99, 130.62, 130.25, 129.79, 129.72, 128.77, 127.31, 35.26, 31.92, 30.32, 22.68, 14.36, 13.09, 11.75.
**HRMS (ESI)** m/z calcd for $C_{30}H_{28}ClFN_6NaO_4S_2^+$ [M+Na]$^+$: 677.1178; found: 677.1183

**(*S*)-butyl (4-(4-(2-(4-(4-chlorophenyl)-2,3,9-trimethyl-6*H*-thieno[3,2-*f*][1,2,4]triazolo[4,3-*a*][1, 4]diazepin-6-yl)acetyl)piperazine-1-carbonyl)phenyl) carbonate (ML1-33)**

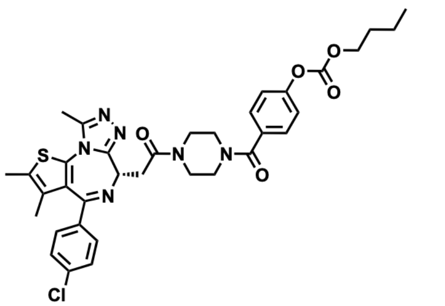

**General Procedure C** was followed with **ML1-26** (35.0 mg, 0.06 mmol), and TFA (0.15 mL, 1.97 mmol) for 2 hours. The crude material was used without further purification.

**General Procedure A** was followed with 4-((butoxycarbonyl)oxy)benzoic acid (18.3 mg, 0.08 mmol), T3P (0.06 mL, 0.10 mmol), DIPEA (0.06 mL, 0.32 mmol), and the amine from above (30.0 mg, 0.06 mmol). The crude residue was purified by silica gel chromatography (0-7% MeOH in DCM) to afford 27.0 mg (61.2%) of the title compound as a yellow-white powder.

**$^{1}$H NMR** (600 MHz, $CDCl_3$) δ 7.48 (d, *J* = 8.6 Hz, 2H), 7.40 (d, *J* = 8.4 Hz, 2H), 7.33 (d, *J* = 8.6 Hz, 2H), 7.27 (d, *J* = 4.3 Hz, 2H), 4.80 (t, *J* = 6.8 Hz, 1H), 4.27 (t, *J* = 6.7 Hz, 2H), 3.93 (s, 2H), 3.80 (dd, *J* = 15.8, 6.9 Hz, 2H), 3.54 (s, 6H), 2.66 (s, 3H), 2.39 (s, 3H), 1.78 – 1.70 (m, 2H), 1.67 (s, 3H), 1.51 – 1.44 (m, 2H), 0.97 (t, *J* = 7.4 Hz, 3H).

**$^{13}$C NMR** (151 MHz, $CDCl_3$) δ 169.74, 169.35, 163.87, 155.72, 153.30, 152.30, 149.89, 136.76, 136.75, 132.83, 132.23, 130.92, 130.74, 130.49, 129.79, 128.74, 128.72, 121.37, 68.97, 54.55, 35.30, 30.57, 29.68, 18.91, 14.35, 13.63, 13.07, 11.81.

**HRMS (ESI)** m/z calcd for $C_{35}H_{37}ClN_6NaO_5S^+$ $[M+Na]^+$: 711.2127; found: 711.2125

**(*S*)-2-(2-(4-(2-(4-(4-chlorophenyl)-2,3,9-trimethyl-6*H*-thieno[3,2-*f*][1,2,4]triazolo[4,3-*a*][1,4]diazepin-6-yl)acetyl)piperazin-1-yl)-2-oxoethoxy)isoindoline-1,3-dione (ML1-43)**

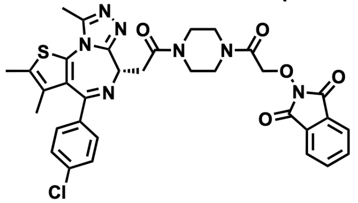

**General Procedure C** was followed with **ML1-26** (35.0 mg, 0.06 mmol), and TFA (0.15 mL, 1.97 mmol) for 2 hours. The crude material was used without further purification.

**General Procedure B** was followed with 2-((1,3-dioxoisoindolin-2-yl)oxy)acetic acid (15.6 mg, 0.07 mmol), HATU (29.3 mg, 0.08 mmol), DIPEA (0.05 mL, 0.26 mmol), and the amine from above (30.0 mg, 0.06 mmol). The crude residue was purified by silica gel chromatography (0-7% MeOH in DCM) to afford 5.0 mg (11.6%) of the title compound as a yellow-white powder.

**$^{1}$H NMR** (500 MHz, $CDCl_3$) δ 7.88 (dd, *J* = 5.4, 3.0 Hz, 2H), 7.76 (dd, *J* = 5.5, 3.1 Hz, 2H), 7.35 (d, *J* = 8.5 Hz, 2H), 7.26 (d, *J* = 8.7 Hz, 2H), 4.74 (dd, *J* = 7.2, 5.9 Hz, 1H), 4.69 (s, 2H), 3.74 (q, *J* = 5.3 Hz, 2H), 3.71 – 3.62 (m, 2H), 3.63 – 3.51 (m, 2H), 2.77 (ddd, *J* = 6.4, 4.0, 2.2 Hz, 2H), 2.70 – 2.65 (m, 2H), 2.63 (s, 3H), 2.37 (s, 3H), 1.64 (s, 3H).

**$^{13}$C NMR** (151 MHz, $CDCl_3$) δ 167.76, 162.97, 149.94, 136.76, 136.72, 134.76, 132.47, 132.06, 131.00, 130.92, 130.88, 130.64, 129.85, 128.85, 128.81, 128.74, 123.81, 77.24, 77.03, 76.82, 68.17, 38.75, 30.37, 28.93, 23.76, 22.98, 14.39, 14.04, 13.10, 11.79, 10.96.

**HRMS (ESI)** m/z calcd for $C_{33}H_{30}ClN_7NaO_5S^+$ $[M+Na]^+$: 694.1610; found: 694.1589

**(*S*)-4-((4-(2-(4-(4-chlorophenyl)-2,3,9-trimethyl-6*H*-thieno[3,2-*f*][1,2,4]triazolo[4,3-*a*][1,4]diazepin-6-yl)acetyl)piperazin-1-yl)methyl)benzenesulfonyl fluoride (ML1-45)**

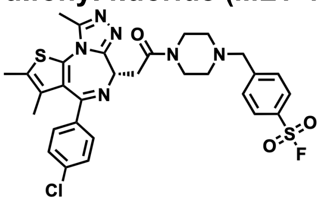

**General Procedure C** was followed with **ML1-26** (35.0 mg, 0.06 mmol), and TFA (0.15 mL, 1.97 mmol) for 2 hours. The crude material was used without further purification.

To a mixture of the above product (30.0 mg, 0.06 mmol) and 4-(bromomethyl)benzenesulfonyl fluoride (17.8 mg, 0.07 mmol) in DMF (1mL) was added DIPEA (0.06 mL, 0.35 mmol). The reaction was stirred at ambient temperature for 20 min and concentrated *in vacuo* before it was redissolved in EtOAc (20 mL) and saturated aqueous sodium bicarbonate (10 mL). The aqueous phase was extracted 3 times with EtOAc, and the organic extracts were washed with brine (10 mL), dried over Na2SO4, vacuum filtered, and concentrated *in vacuo*. The resultant residue was purified by silica gel chromatography (0-5% MeOH in DCM) to afford 19.9 mg (48.5 %) of the title compound as a yellow-white powder.
**$^{1}$H NMR** (600 MHz, $CDCl_3$) δ 7.98 (d, *J* = 8.4 Hz, 2H), 7.64 (d, *J* = 8.2 Hz, 2H), 7.40 (d, *J* = 8.5 Hz, 2H), 7.32 (d, *J* = 8.4 Hz, 2H), 4.83 – 4.77 (m, 1H), 3.90 – 3.84 (m, 1H), 3.80 (ddd, *J* = 13.2, 6.4, 3.3 Hz, 1H), 3.71 (dt, *J* = 15.8, 4.8 Hz, 2H), 3.66 (s, 2H), 3.59 (dtd, *J* = 19.9, 15.2, 9.3 Hz, 2H), 2.66 (s, 3H), 2.63 – 2.54 (m, 2H), 2.50 (ddd, *J* = 10.0, 6.4, 3.4 Hz, 1H), 2.43 (ddd, *J* = 11.1, 7.5, 3.3 Hz, 1H), 2.39 (s, 3H), 1.67 (s, 3H).
**$^{13}$C NMR** (151 MHz, $CDCl_3$) δ 168.99, 163.67, 155.89, 149.83, 147.10, 136.85, 136.66, 132.23, 131.90, 131.74, 130.92, 130.66, 130.56, 129.83, 129.80, 128.68, 128.60, 62.03, 54.44, 53.30, 52.86, 45.78, 41.81, 35.23, 14.35, 13.06, 11.83.
**HRMS (ESI)** m/z calcd for $C_{30}H_{30}ClFN_6NaO_3S_2^+$ $[M+Na]^+$: 663.1386; found: 663.1387

**(*S*)-2-(4-(4-chlorophenyl)-2,3,9-trimethyl-6*H*-thieno[3,2-*f*][1,2,4]triazolo[4,3-*a*][1,4]diazepin-6-yl)-1-(4-(vinylsulfonyl)piperazin-1-yl)ethan-1-one (ML1-50)**

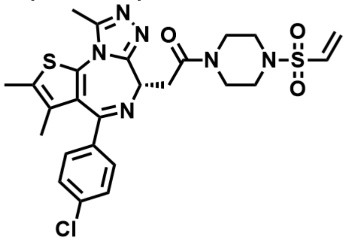

**General Procedure C** was followed with **ML1-26** (200.0 mg, 0.35 mmol), and TFA (0.86 mL, 11.25 mmol) for 2 hours. The crude material was used without further purification.
**General Procedure D** was followed with the above product (164.8 mg, 0.06 mmol), triethylamine (0.25 mL, 1.76 mmol), and 2-chloroethanesulfonyl chloride (0.04 mL, 0.39 mmol). The crude residue was purified by silica gel chromatography (0-5% MeOH in DCM) to afford 103.5mg (52.7%) of the title compound as a yellow-white powder.
**$^{1}$H NMR** (600 MHz, $CDCl_3$) δ 7.38 (d, *J* = 8.5 Hz, 2H), 7.32 (d, *J* = 8.7 Hz, 2H), 6.44 (dd, *J* = 16.6, 10.0 Hz, 1H), 6.28 (d, *J* = 16.6 Hz, 1H), 6.09 (d, *J* = 10.0 Hz, 1H), 4.77 (t, *J* = 6.8 Hz, 1H), 4.03 – 3.94 (m, 2H), 3.79 – 3.71 (m, 2H), 3.59 – 3.46 (m, 2H), 3.38 (ddd, *J* = 11.8, 5.7, 3.2 Hz, 1H), 3.27 (dddt, *J* = 22.7, 11.6, 8.1, 3.4 Hz, 2H), 3.05 (ddd, *J* = 11.7, 8.2, 3.3 Hz, 1H), 2.65 (s, 3H), 2.39 (s, 3H), 1.67 (s, 3H).
**$^{13}$C NMR** (151 MHz, $CDCl_3$) δ 177.74, 172.47, 164.27, 158.48, 145.34, 145.32, 140.79, 140.77, 139.50, 139.33, 139.04, 138.36, 138.06, 137.30, 85.81, 85.60, 85.38, 62.98, 54.25, 54.15, 53.99, 50.01, 43.86, 22.94, 21.66, 20.41.
**HRMS (ESI)** m/z calcd for $C_{25}H_{27}ClN_6NaO_3S_2^+$ $[M+Na]^+$: 581.1167; found: 581.1171

**(*S*)-5-(4-(2-(4-(4-chlorophenyl)-2,3,9-trimethyl-6*H*-thieno[3,2-*f*][1,2,4]triazolo[4,3-*a*][1,4]diazepin-6-yl)acetyl)piperazine-1-carbonyl)picolinaldehyde (ML1-52)**

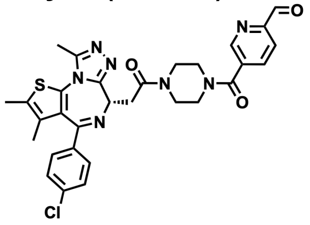

**General Procedure C** was followed with **ML1-26** (35.0 mg, 0.06 mmol), and TFA (0.15 mL, 1.97 mmol) for 2 hours. The crude material was used without further purification.
**General Procedure B** was followed with 6-formylnicotinic acid (10.2 mg, 0.07 mmol), HATU (28.1 mg, 0.07 mmol), DIPEA (0.04 mL, 0.25 mmol), and the amine from above (28.8 mg, 0.06 mmol). The crude residue was purified by silica gel chromatography (0-10% MeOH in DCM) to afford 15.0 mg (40.5%) of the title compound as a yellow-white powder.

**$^1$H NMR** (600 MHz, $CDCl_3$) δ 10.12 (s, 1H), 8.85 (s, 1H), 8.05 (d, *J* = 7.9 Hz, 1H), 7.95 (dd, *J* = 8.0, 2.0 Hz, 1H), 7.40 (d, *J* = 8.3 Hz, 2H), 7.33 (d, 2H), 4.80 (t, *J* = 6.9 Hz, 1H), 4.10 – 4.01 (m, 2H), 3.88 – 3.75 (m, 2H), 3.76 – 3.64 (m, 1H), 3.62 – 3.34 (m, 1H), 2.66 (s, 3H), 2.40 (s, 3H), 1.68 (s, 3H).
**$^{13}$C NMR** (151 MHz, $CDCl_3$) δ 192.43, 171.12, 166.92, 153.35, 148.43, 136.82, 136.74, 136.16, 134.90, 132.26, 130.92, 130.44, 129.77, 128.75, 121.49, 60.38, 29.69, 21.03, 14.36, 13.08, 11.83.
**HRMS (ESI)** m/z calcd for $C_{30}H_{28}ClN_7NaO_3S^+$ $[M+Na]^+$: 624.1555; found: 624.1563

**(*S*)-3-(4-(2-(4-(4-chlorophenyl)-2,3,9-trimethyl-6*H*-thieno[3,2-*f*][1,2,4]triazolo[4,3-*a*][1,4]diazepin-6-yl)acetyl)piperazine-1-carbonyl)-4-hydroxybenzenesulfonyl fluoride (ML1-54)**

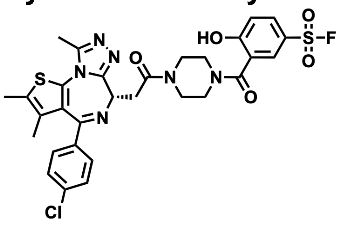

**General Procedure C** was followed with **ML1-26** (30.0 mg, 0.05 mmol), and TFA (0.13 mL, 1.69 mmol) for 2 hours. The crude material was used without further purification.
**General Procedure A** was followed with 5-(fluorosulfonyl)-2-hydroxybenzoic acid (13.9 mg, 0.06 mmol), T3P (0.04 mL, 0.06 mmol), DIPEA (0.05 mL, 0.26 mmol), and the amine from above (24.7 mg, 0.05 mmol). The crude residue was purified by silica gel chromatography (0-8% MeOH in DCM) to afford 5.5 mg (15.6%) of the title compound as a yellow-white powder.
**$^1$H NMR** (600 MHz, $CDCl_3$) δ 7.97 – 7.91 (m, 2H), 7.41 (d, *J* = 8.4 Hz, 2H), 7.34 (d, *J* = 8.7 Hz, 2H), 7.25 (d, *J* = 8.7 Hz, 1H), 4.82 (dd, *J* = 7.7, 6.1 Hz, 1H), 4.09 – 4.04 (m, 2H), 3.97 – 3.89 (m, 2H), 3.87 (dd, *J* = 15.8, 7.9 Hz, 1H), 3.77 (dd, *J* = 13.0, 5.7 Hz, 2H), 3.68 – 3.63 (m, 1H), 3.61 – 3.53 (m, 1H), 3.45 (dd, *J* = 15.8, 6.1 Hz, 1H), 2.68 (s, 3H), 2.42 (s, 3H), 1.69 (s, 3H).
**$^{13}$C NMR** (151 MHz, $CDCl_3$) δ 169.32, 168.37, 155.72, 149.97, 136.93, 136.59, 132.57, 132.02, 131.03, 130.98, 130.64, 129.87, 129.79, 129.59, 128.79, 119.25, 118.87, 54.58, 41.51, 35.35, 14.37, 13.10, 11.81.
**HRMS (ESI)** m/z calcd for $C_{30}H_{28}ClFN_6NaO_5S_2^+$ $[M+Na]^+$: 693.1127; found: 693.1132

**(*S*)-2,4-di-*tert*-butyl-5-(2-(4-(4-chlorophenyl)-2,3,9-trimethyl-6*H*-thieno[3,2-*f*][1,2,4]triazolo[4,3-*a*][1,4]diazepin-6-yl)acetamido)phenyl methyl carbonate (ML1-55)**

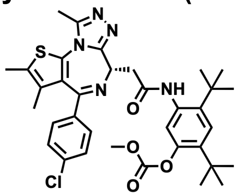

**General Procedure A** was followed with (S)-2-(4-(4-chlorophenyl)-2,3,9-trimethyl-6H-thieno [3,2-f][1,2,4]triazolo[4,3-a][1,4]diazepin-6-yl)acetic acid (JQ1-Acid) (25.0 mg, 0.06 mmol), T3P (0.05 mL, 0.07 mmol), DIPEA (0.04 mL, 0.25 mmol), and 5-amino-2,4-di-tert-butylphenyl methyl carbonate (19.2 mg, 0.07 mmol). The crude residue was purified by silica gel chromatography (0-8% MeOH in DCM) to afford 5.8 mg (14.0%) of the title compound as a yellow-white powder.
**$^1$H NMR** (400 MHz, $CDCl_3$) δ 8.03 (s, 1H), 7.49 (s, 1H), 7.42 (s, 1H), 7.39 (d, *J* = 5.1 Hz, 2H), 7.32 (d, *J* = 8.5 Hz, 2H), 4.72 (t, *J* = 6.7 Hz, 1H), 3.87 (s, 3H), 3.71 (dd, *J* = 14.6, 6.2 Hz, 1H), 3.59 (dd, *J* = 14.6, 7.2 Hz, 1H), 2.68 (s, 3H), 2.41 (s, 3H), 1.68 (s, 3H), 1.46 (s, 9H), 1.34 (s, 9H).
**$^{13}$C NMR** (151 MHz, $CDCl_3$) δ 168.68, 164.01, 155.70, 154.24, 149.99, 147.53, 139.50, 137.85, 136.82, 136.56, 133.56, 132.14, 131.00, 130.85, 130.53, 129.93, 128.68, 125.25, 121.85, 55.31, 54.23, 40.46, 34.78, 34.72, 30.79, 30.22, 14.40, 13.09, 11.81.
**HRMS (ESI)** m/z calcd for $C_{35}H_{40}ClN_5NaO_4S^+$ $[M+Na]^+$: 684.2382; found: 684.2385

**7-cyclopentyl-*N*,*N*-dimethyl-2-((5-(4-(vinylsulfonyl)piperazin-1-yl)pyridin-2-yl)amino)-7*H*-pyrrolo[2,3-*d*]pyrimidine-6-carboxamide (ML1-71)**

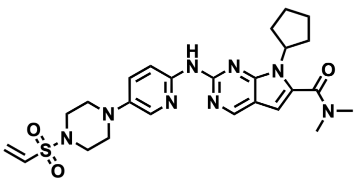

**General Procedure D** was followed with 7-cyclopentyl-N,N-dimethyl-2-((5-(piperazin-1-yl) pyridin-2-yl)amino)-7H-pyrrolo[2,3-d]pyrimidine-6-carboxamide (ribociclib) (50.0 mg, 0.12 mmol), triethylamine (0.05 mL, 0.35 mmol), and 2-chloroethanesulfonyl chloride (0.01 mL, 0.13 mmol). The crude residue was purified by silica gel chromatography (0-8% MeOH in DCM) to afford 21.7 mg (36.0%) of the title compound as a yellow-white powder.
**$^{1}$H NMR** (600 MHz, $CDCl_3$) δ 8.69 (s, 1H), 8.38 (d, J = 8.3 Hz, 1H), 8.00 (d, J = 2.3 Hz, 1H), 7.87 (s, 1H), 7.32 (dd, J = 9.1, 2.9 Hz, 1H), 6.48 (dd, J = 16.6, 10.0 Hz, 1H), 6.44 (s, 1H), 6.30 (d, J = 16.6 Hz, 1H), 6.10 (d, J = 10.0 Hz, 1H), 4.80 (p, J = 8.9 Hz, 1H), 3.38 – 3.34 (m, 4H), 3.25 – 3.20 (m, 5H), 2.58 (qd, J = 10.1, 5.0 Hz, 2H), 2.12 – 2.00 (m, 5H), 1.75 – 1.68 (m, 2H).
**$^{13}$C NMR** (151 MHz, $CDCl_3$) δ 164.06, 154.47, 151.96, 151.77, 147.78, 141.85, 132.32, 132.18, 129.16, 128.34, 127.52, 112.80, 112.37, 100.94, 57.90, 50.22, 45.57, 30.20, 24.73.
**HRMS (ESI)** m/z calcd for $C_{25}H_{33}N_8O_3S^+$ $[M+Na]^+$: 525.2391; found: 525.2382

**2-(6-amino-5-(4-(vinylsulfonyl)piperazin-1-yl)pyridazin-3-yl)phenol (ML1-96)**

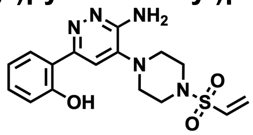

**General Procedure D** was followed with 2-(6-amino-5-(piperazin-1-yl)pyridazin-3-yl)phenol (50.0 mg, 0.18 mmol), triethylamine (0.08 mL, 0.55 mmol), and 2-chloroethanesulfonyl chloride (0.02 mL, 0.15 mmol). The crude residue was purified by silica gel chromatography (0-8% MeOH in DCM) and another round of purification using HPLC (buffer B : buffer A = 25:75 to 60:40) to afford 12.4 mg (18.6%) of the title compound as a yellow-white powder.
**$^{1}$H NMR** (600 MHz, $CDCl_3$) δ 7.61 (dd, *J* = 8.0, 1.6 Hz, 1H), 7.38 (s, 1H), 7.33 (ddd, *J* = 8.5, 7.3, 1.6 Hz, 1H), 7.09 (dd, *J* = 8.2, 1.2 Hz, 1H), 6.99 – 6.92 (m, 1H), 6.52 (dd, *J* = 16.6, 9.9 Hz, 1H), 6.37 (d, *J* = 16.6 Hz, 1H), 6.16 (d, *J* = 9.9 Hz, 1H), 4.81 (s, 2H), 3.43 (t, *J* = 4.9 Hz, 4H), 3.28 (t, *J* = 4.9 Hz, 4H).
**$^{13}$C NMR** (151 MHz, $CDCl_3$) δ 159.23, 155.52, 153.88, 140.25, 132.32, 131.13, 129.58, 125.18, 118.82, 118.62, 117.35, 111.46, 48.91, 45.44.
**HRMS (ESI)** m/z calcd for $C_{16}H_{20}N_5O_3S^+$ $[M+H]^+$: 362.1281; found: 362.1291

**(*R*)-3-(4-phenoxyphenyl)-1-(1-(vinylsulfonyl)piperidin-3-yl)-1*H*-pyrazolo[3,4-*d*]pyrimidin-4-amine (TH1-9)**

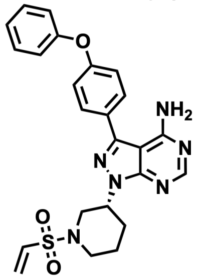

**General Procedure C** was followed with tert-butyl (R)-3-(4-amino-3-(4-phenoxyphenyl)-1H- pyrazolo[3,4-d]pyrimidin-1-yl)piperidine-1-carboxylate (100.0mg, 0.21mmol), and TFA (0.5 mL, 6.58 mmol) overnight. The crude material was used without further purification.
**General Procedure D** was followed with the above product (50.0mg, 0.13mmol), triethylamine (0.08 mL, 0.58 mmol), and 2-chloroethanesulfonyl chloride (0.02 mL, 0.19 mmol). The crude residue was purified by silica gel chromatography (0-10% MeOH in DCM) to afford 18.2 mg (29.5%) of the title compound as a yellow-white powder.
**$^{1}$H NMR** (600 MHz, $CDCl_3$) δ 8.37 (s, 1H), 7.62 (d, *J* = 8.6 Hz, 2H), 7.38 (dd, *J* = 8.6, 7.4 Hz, 2H), 7.17 (t, 1H), 7.15 (d, 2H), 7.07 (d, *J* = 7.6 Hz, 1H), 6.46 (dd, *J* = 16.6, 10.0 Hz, 1H), 6.24 (d, *J* = 16.6 Hz, 1H), 6.02 (d, *J* = 10.0 Hz, 1H), 5.50 (s, 2H), 5.05 – 4.97 (m, 1H), 4.01 – 3.94 (m, 1H), 3.81 (dt, *J* = 12.2, 2.3 Hz, 1H), 3.28 (t, *J* = 11.2 Hz, 1H), 2.72 (td, *J* = 12.2, 2.9 Hz, 1H), 2.24 – 2.18 (m, 2H), 2.02 – 1.95 (m, 1H), 1.92 – 1.83 (m, 1H).
**$^{13}$C NMR** (151 MHz, $CDCl_3$) δ 158.64, 157.76, 156.32, 156.00, 154.38, 144.04, 132.98, 129.99, 129.96, 128.57, 127.65, 124.11, 119.58, 119.13, 98.62, 52.89, 49.19, 45.52, 29.57, 24.37.

**HRMS (ESI)** m/z calcd for $C_{24}H_{24}N_6NaO_3S^+$ $[M+Na]^+$: 499.1523; found: 499.1523

***N*-(2-chloro-6-methylphenyl)-2-((2-methyl-6-(4-(vinylsulfonyl)piperazin-1-yl)pyrimidin-4-yl)amino)thiazole-5-carboxamide (ML2-5)**

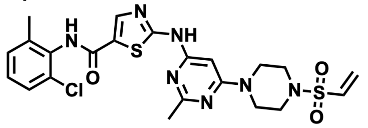

**General Procedure D** was followed with N-(2-chloro-6-methylphenyl)-2-((2-methyl-6-(piperazin- 1-yl)pyrimidin-4-yl)amino)thiazole-5-carboxamide (100.0 mg, 0.23 mmol), triethylamine (0.09 mL, 0.68 mmol), and 2-chloroethanesulfonyl chloride (0.02 mL, 0.23 mmol). The crude residue was purified by silica gel chromatography (50-100% EtOAc in Hexanes) to afford 35.6 mg (29.6%) of the title compound as a yellow-white powder.
**$^1$H NMR** (600 MHz, DMSO) δ 11.52 (s, 1H), 9.87 (s, 1H), 8.23 (s, 1H), 7.40 (dd, *J* = 7.8, 1.7 Hz, 1H), 7.28 (dt, *J* = 15.4, 7.5 Hz, 2H), 6.83 (dd, *J* = 16.5, 10.0 Hz, 1H), 6.20 (d, *J* = 10.0 Hz, 1H), 6.16 (d, *J* = 16.5 Hz, 1H), 6.11 (s, 1H), 3.66 (t, *J* = 5.1 Hz, 4H), 3.31 (s, 3H), 3.14 (t, *J* = 5.1 Hz, 4H), 2.43 (s, 3H), 2.25 (s, 3H).
**$^{13}$C NMR** (151 MHz, DMSO) δ 170.21, 165.22, 162.37, 162.08, 159.78, 157.01, 140.71, 138.71, 133.41, 132.42, 132.33, 129.64, 128.92, 128.07, 126.90, 125.73, 82.99, 59.63, 44.75, 43.02, 25.44, 20.65, 18.18, 13.98.
**HRMS (ESI)** m/z calcd for $C_{22}H_{24}ClN_7NaO_3S_2^+$ $[M+Na]^+$: 556.0963; found: 556.0966

***tert*-butyl 4-(6-(((1*r*,4*r*)-4-(3-chloro-4-cyanophenoxy)cyclohexyl)carbamoyl)pyridazin-3-yl) piperazine-1-carboxylate (ML2-8)**

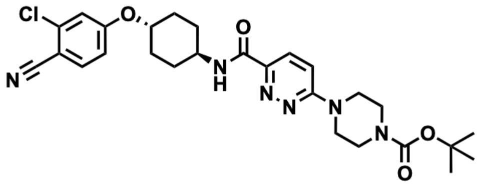

To 6-chloro-N-((1r,4r)-4-(3-chloro-4-cyanophenoxy)cyclohexyl)pyridazine-3-carboxamide (200.0 mg, 0.51 mmol) in DMF (4 mL) was added 1-boc-piperazine (142.8 mg, 0.77 mmol), followed by triethylamine (0.21 mL, 1.53 mmol). The resultant mixture was stirred at 80 °C overnight. It was then cooled to RT and the precipitate was filtered off and dried to afford 156.6mg (56.6%) of the title compound as a white powder.
**$^1$H NMR** (400 MHz, $CDCl_3$) δ 8.01 (d, *J* = 9.5 Hz, 1H), 7.84 (d, *J* = 8.5 Hz, 1H), 7.54 (d, *J* = 8.6 Hz, 1H), 7.01 – 6.94 (m, 2H), 6.84 (dd, *J* = 8.8, 2.5 Hz, 1H), 4.31 (tt, *J* = 9.9, 3.6 Hz, 1H), 4.04 (dtt, *J* = 10.8, 7.4, 3.9 Hz, 1H), 3.78 – 3.71 (m, 4H), 3.62 – 3.54 (m, 4H), 2.16 (tt, *J* = 10.8, 4.2 Hz, 5H), 1.81 – 1.59 (m, 2H), 1.48 (s, 9H), 1.45 – 1.31 (m, 2H).
**$^{13}$C NMR** (151 MHz, $CDCl_3$) δ 162.60, 161.63, 160.20, 154.64, 144.83, 138.35, 135.08, 126.98, 116.94, 116.40, 114.66, 112.28, 104.86, 80.44, 75.68, 47.12, 30.03, 29.67, 28.41.
**HRMS (ESI)** m/z calcd for $C_{27}H_{33}ClN_6NaO_4^+$ $[M+Na]^+$: 563.2144; found: 563.2146

***N*-((1*r*,4*r*)-4-(3-chloro-4-cyanophenoxy)cyclohexyl)-6-(4-(vinylsulfonyl)piperazin-1-yl)pyridazine-3-carboxamide (ML2-9)**

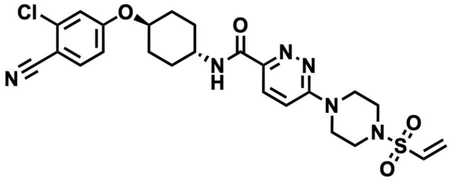

**General Procedure C** was followed with **ML2-8** (70.0 mg, 0.13 mmol), and TFA (0.32 mL, 4.14 mmol) for 2 hours. The crude material was used without further purification.
**General Procedure D** was followed with the above product (57.1 mg, 0.13 mmol), triethylamine (0.08 mL, 0.58 mmol), and 2-chloroethanesulfonyl chloride (0.02 mL, 0.16 mmol). The crude residue was purified by silica gel chromatography (0-100% EtOAc in Hexanes) to afford 27.4mg (39.9%) of the title compound as a yellow-white powder.
**$^1$H NMR** (600 MHz, $CDCl_3$) δ 8.05 (d, *J* = 9.4 Hz, 1H), 7.84 (d, *J* = 8.2 Hz, 1H), 7.55 (d, *J* = 8.7 Hz, 1H), 7.02 (s, 1H), 7.02 – 6.97 (m, 1H), 6.85 (dd, *J* = 8.7, 2.4 Hz, 1H), 6.43 (dd, *J* = 16.6, 9.9 Hz, 1H), 6.30 (d, *J* = 16.6 Hz, 1H), 6.09 (d, *J* = 9.9 Hz, 1H), 4.32 (td, *J* = 10.3, 5.2 Hz, 1H), 4.06 (dtd, *J* = 10.8, 7.4, 4.0 Hz, 1H), 3.89 (t, *J* = 5.1 Hz, 4H), 3.32 (t, *J* = 5.1 Hz, 4H), 2.23 – 2.13 (m, 4H), 1.74 – 1.66 (m, 2H), 1.52 – 1.42 (m, 2H).

**$^{13}$C NMR** (151 MHz, $CDCl_3$) δ 162.38, 161.60, 159.99, 145.30, 138.34, 135.09, 132.10, 129.68, 127.20, 116.91, 116.40, 114.66, 112.65, 104.84, 75.62, 47.18, 45.04, 44.65, 30.02, 29.70, 29.66.
**HRMS (ESI)** m/z calcd for $C_{24}H_{27}ClN_6NaO_4S^+$ $[M+Na]^+$: 553.1395; found: 553.1387

***tert*-butyl 4-(prop-2-yn-1-yl)piperazine-1-carboxylate (ML2-32)**

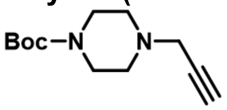


A combination of 1-boc-piperazine (1.50 g, 8.05 mmol) and potassium carbonate (1.67 g, 12.08 mmol) was suspended in acetonitrile (30 mL) and stirred at ambient temperature for 10 minutes. Propargyl bromide (0.84 mL, 8.86 mmol) was added, and the reaction mixture was stirred at 60°C for 2 hours. The reaction mixture was concentrated *in vacuo*, and the crude residue was purified by silica gel chromatography (0-10% MeOH in EtOAc) to afford 1.50 g (82.9%) of the title compound as a yellow oil.
**$^{1}$H NMR** (600 MHz, $CDCl_3$) δ 3.38 (t, *J* = 5.1 Hz, 4H), 3.22 (d, *J* = 2.5 Hz, 2H), 2.42 (t, *J* = 5.1 Hz, 4H), 2.19 (t, *J* = 2.4 Hz, 1H), 1.37 (s, 9H).
**$^{13}$C NMR** (151 MHz, $CDCl_3$) δ 154.55, 79.54, 78.35, 73.41, 51.54, 46.90, 28.35.

**1-(prop-2-yn-1-yl)-4-(vinylsulfonyl)piperazine (ML2-33)**

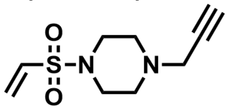


**General Procedure C** was followed with **ML2-32** (500.0 mg, 2.23 mmol), and TFA (5.5 mL, 73.3 mmol) for 2 hours. The crude material was used without further purification.
**General Procedure D** was followed with the above product (28 mg, 0.23 mmol), triethylamine (0.14 mL, 1.01 mmol), and 2-chloroethanesulfonyl chloride (0.03 mL, 0.27 mmol). The reaction mixture was concentrated *in vacuo*, and the crude residue was purified by silica gel chromatography (0-10% MeOH in EtOAc) to afford 13.0 mg (26.9%) of the title compound as a yellow oil.
**$^{1}$H NMR** (600 MHz, $CDCl_3$) δ 6.42 (dd, *J* = 16.6, 10.0 Hz, 1H), 6.24 (d, *J* = 16.6 Hz, 1H), 6.04 (d, *J* = 10.0 Hz, 1H), 3.33 (d, *J* = 2.4 Hz, 2H), 3.22 (t, *J* = 5.0 Hz, 4H), 2.68 – 2.63 (m, 4H), 2.28 (t, *J* = 2.5 Hz, 1H).
**$^{13}$C NMR** (151 MHz, $CDCl_3$) δ 132.32, 128.81, 77.98, 73.72, 51.10, 46.68, 45.45.

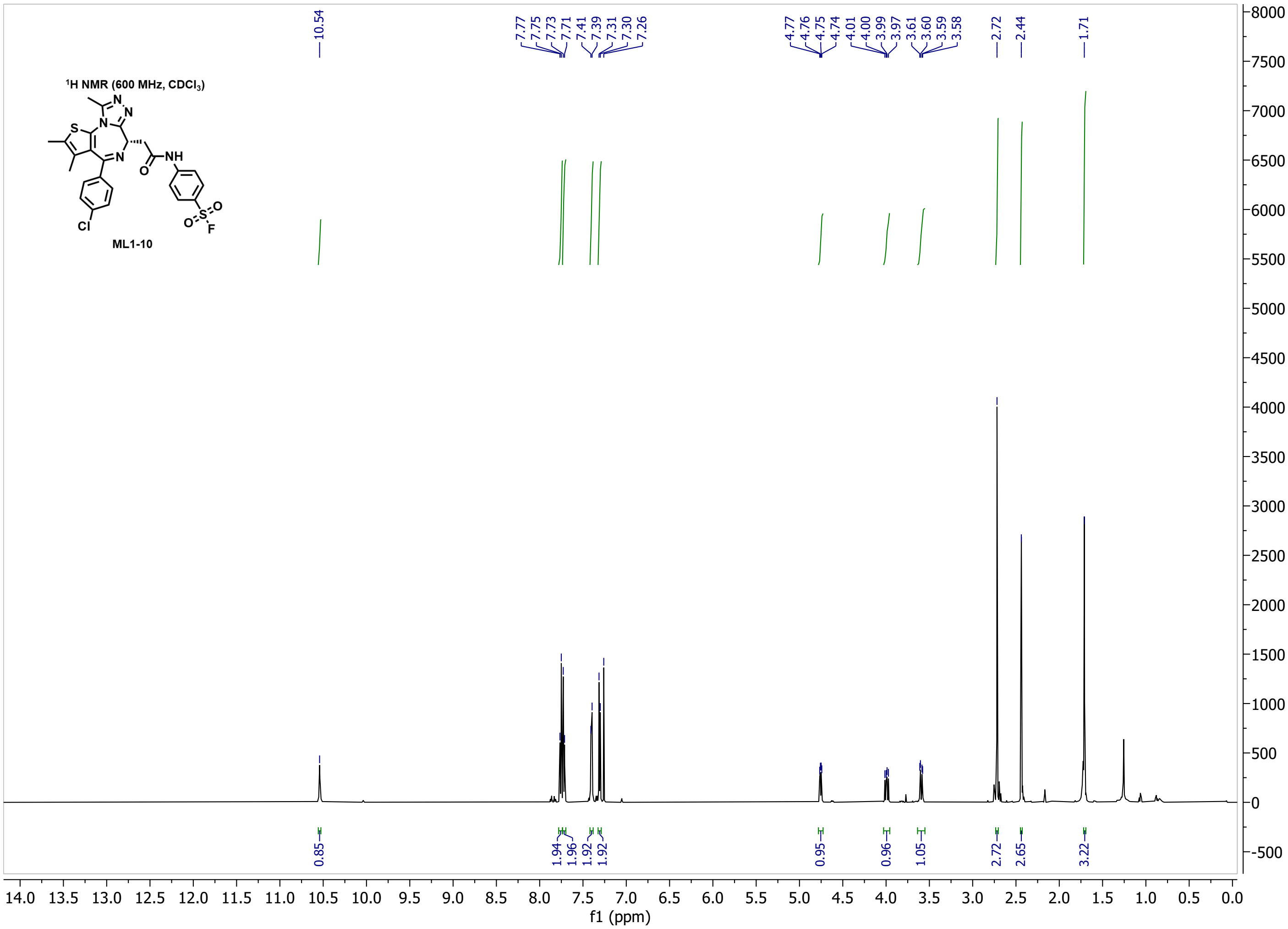

$^1$H NMR (600 MHz, CDCl$_3$)
ML1-10
10.54
7.77 7.75 7.73 7.71 7.41 7.39 7.31 7.30 7.26
4.77 4.76 4.75 4.74 4.01 4.00 3.99 3.97 3.61 3.60 3.59 3.58
2.72
2.44
1.71
0.85 1.94 1.96 1.92 1.92 0.95 0.96 1.05 2.72 2.65 3.22
f1 (ppm)


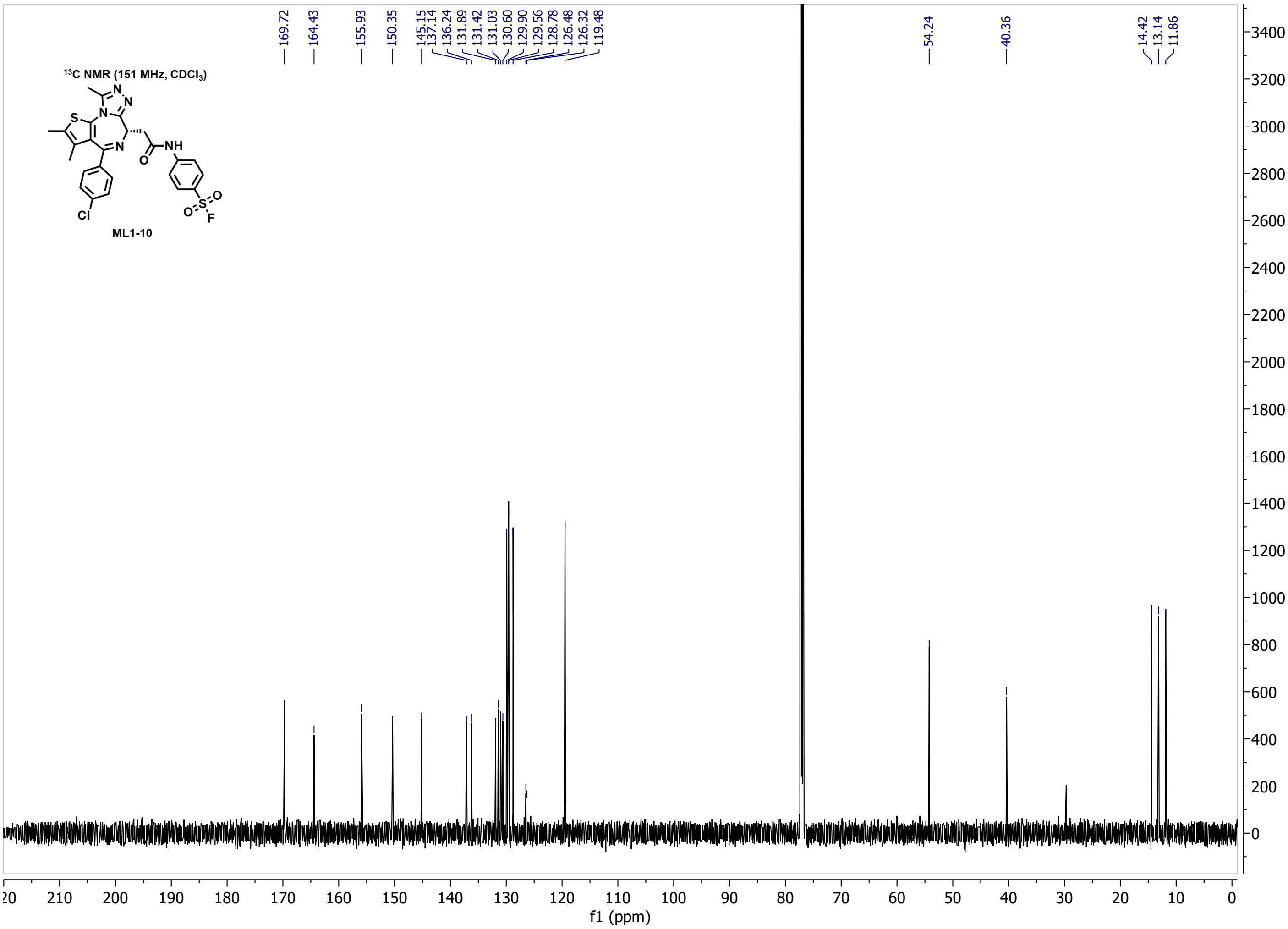

$^{13}$C NMR (151 MHz, CDCl$_3$)
ML1-10
169.72
164.43
155.93
150.35
145.15 137.14 136.24 131.89 131.42 131.03 130.60 129.90 129.56 128.78 126.48 126.32 119.48
54.24
40.36
14.42 13.14 11.86
f1 (ppm)

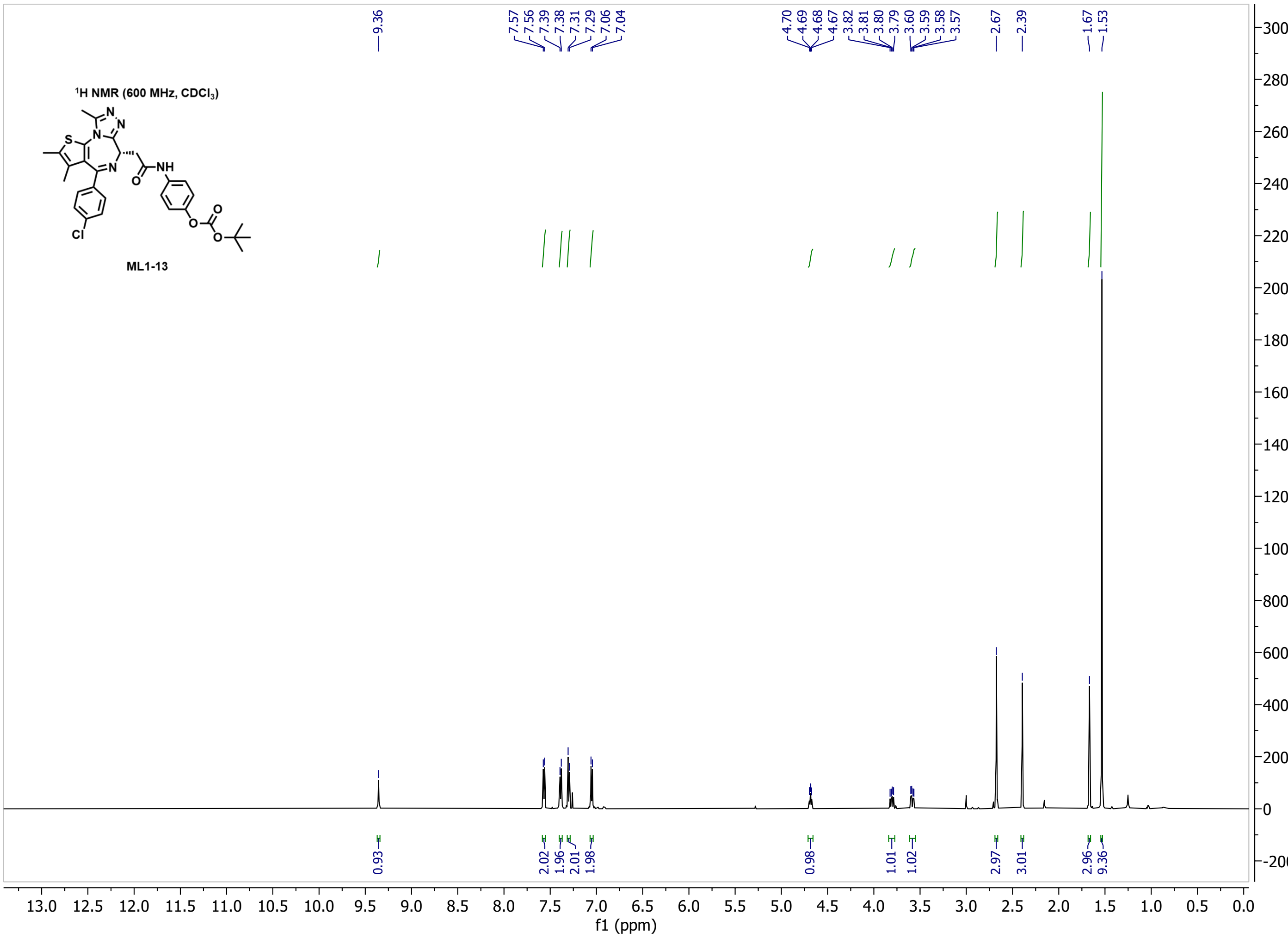
1H NMR (600 MHz, CDCl3)
ML1-13
9.36
7.57 7.56 7.39 7.38 7.31 7.29 7.06 7.04
4.70 4.69 4.68 4.67 3.82 3.81 3.80 3.79 3.60 3.59 3.58 3.57
2.67 2.39 1.67 1.53
0.93 2.02 1.96 2.01 1.98 0.98 1.01 1.02 2.97 3.01 2.96 9.36
f1 (ppm)

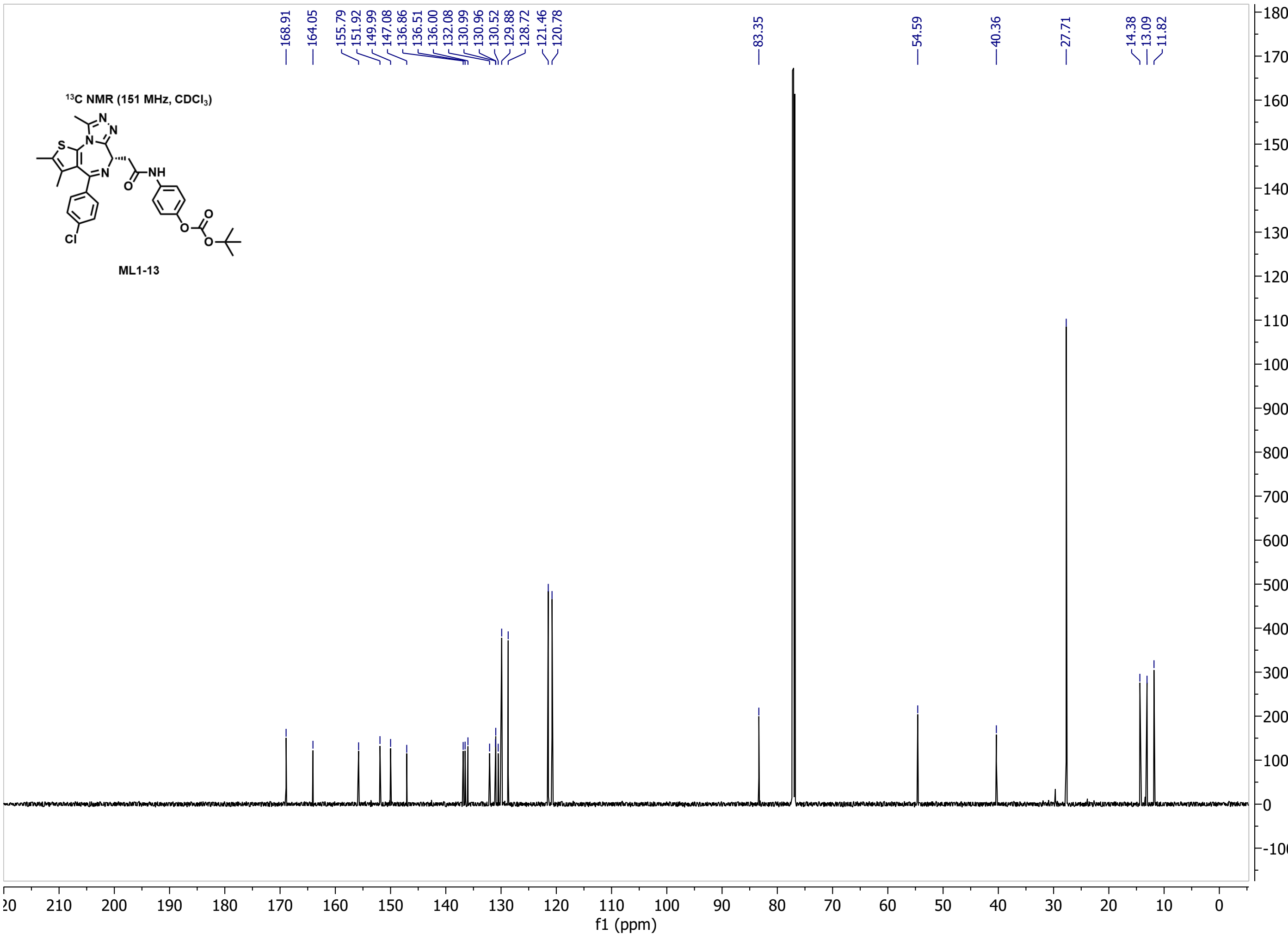
13C NMR (151 MHz, CDCl3)
ML1-13
168.91 164.05 155.79 151.92 149.99 147.08 136.86 136.51 136.00 132.08 130.99 130.96 130.52 129.88 128.72 121.46 120.78
83.35
54.59
40.36
27.71
14.38 13.09 11.82
f1 (ppm)

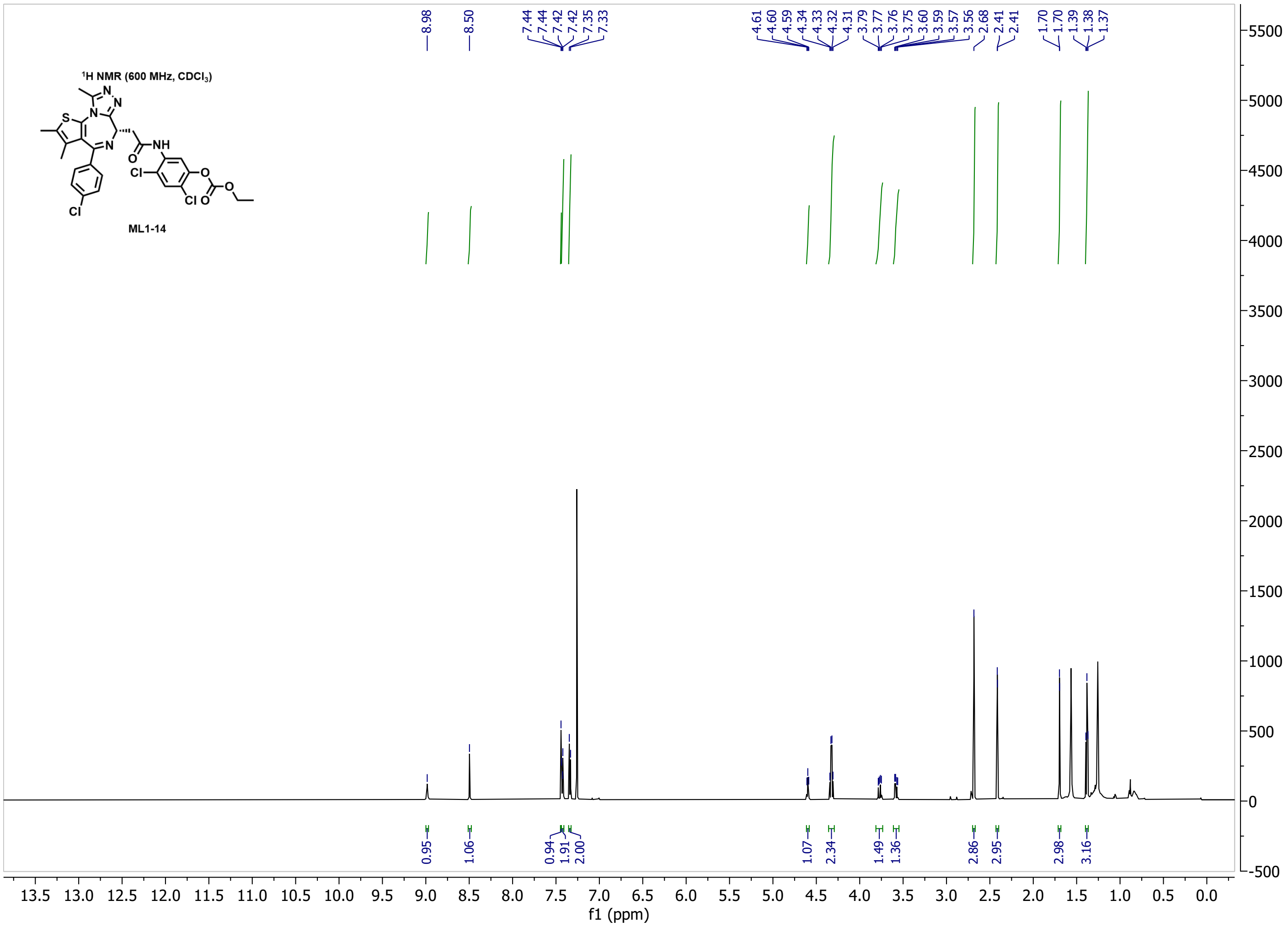

¹H NMR (600 MHz, CDCl3)
ML1-14
f1 (ppm)


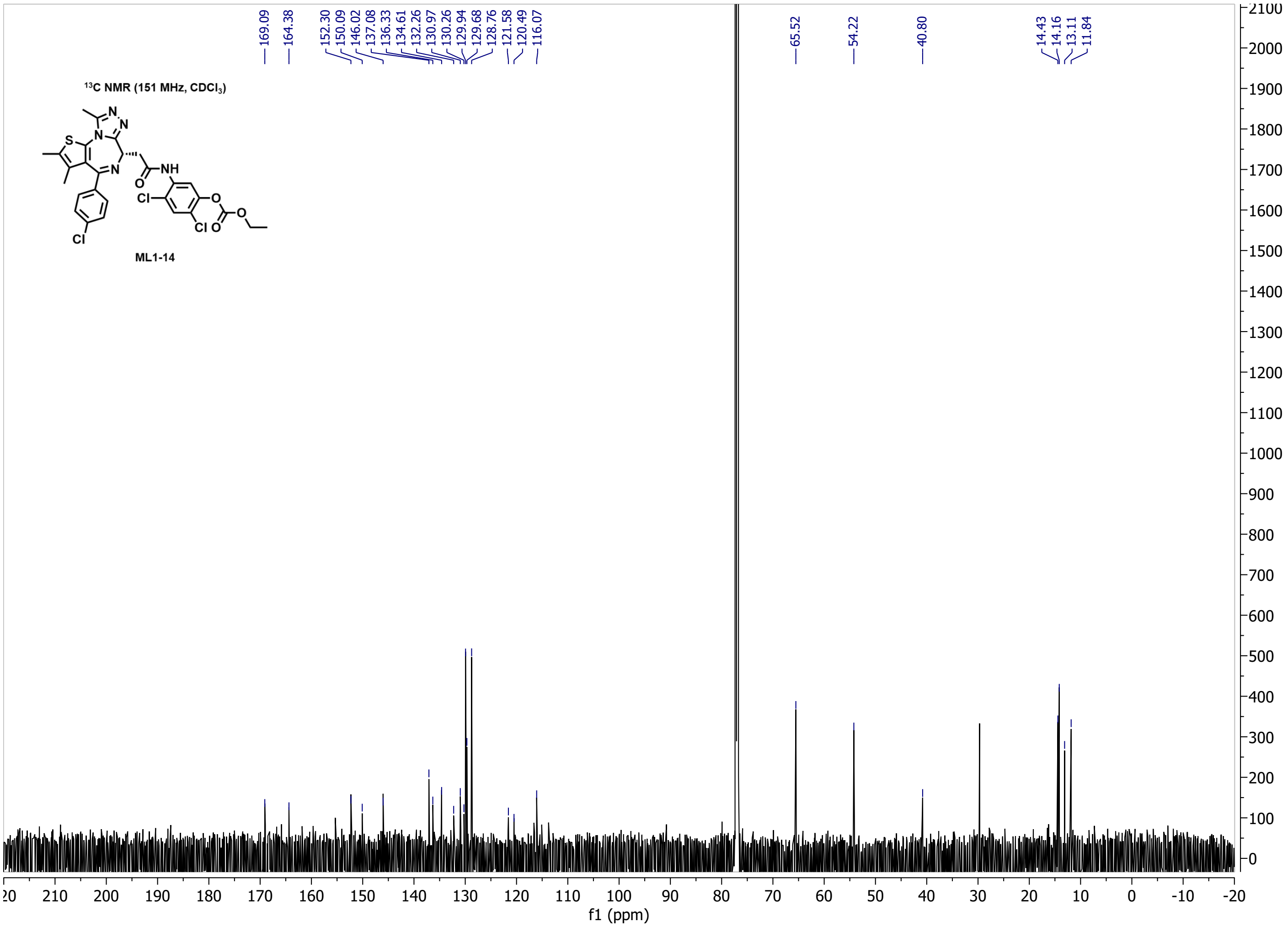

¹³C NMR (151 MHz, CDCl3)
ML1-14
f1 (ppm)

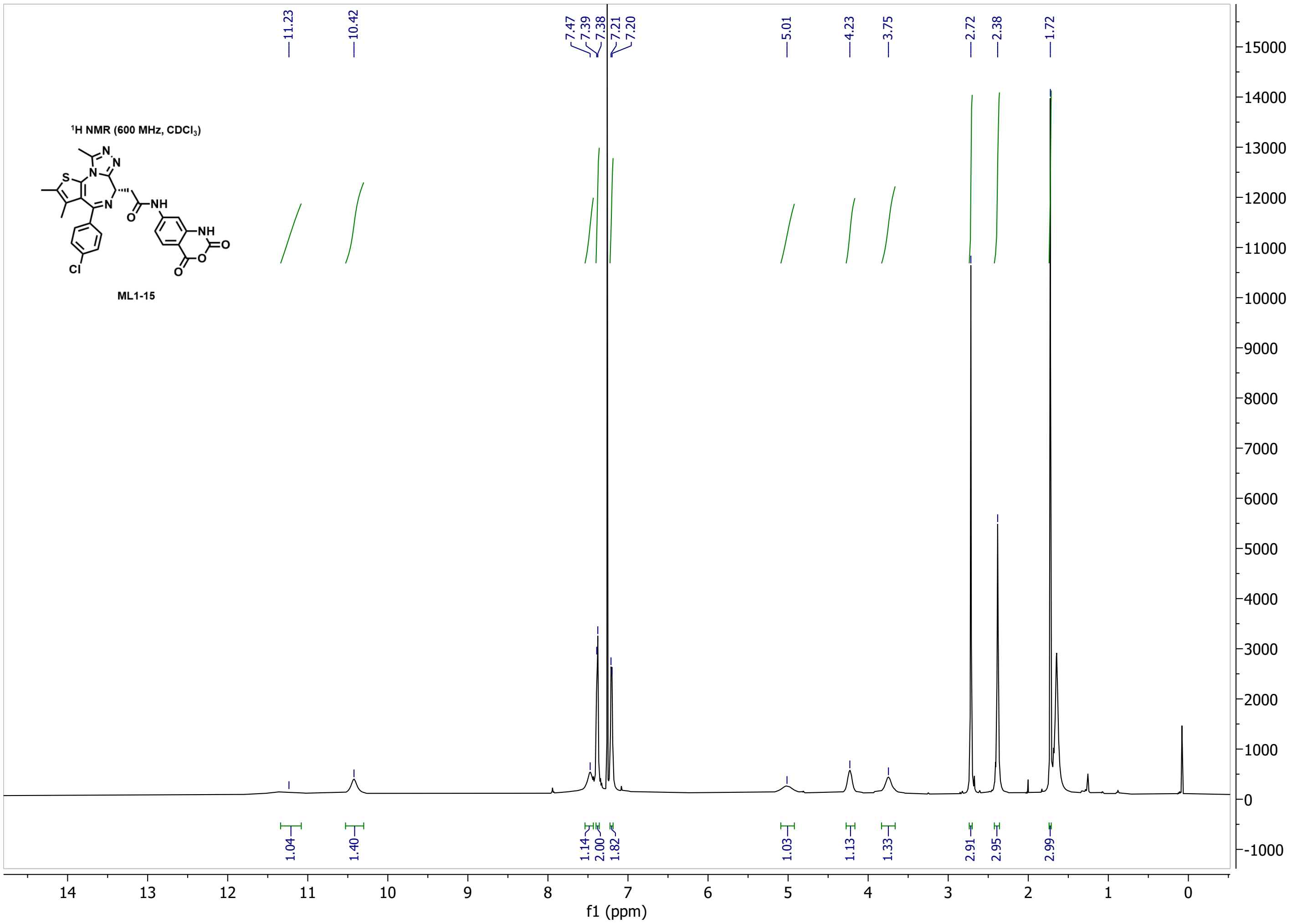
1H NMR (600 MHz, CDCl3)
ML1-15
NH
O
S
N
Cl
11.23
10.42
7.47
7.39
7.38
7.21
7.20
5.01
4.23
3.75
2.72
2.38
1.72
1.04
1.40
1.14
2.00
1.82
1.03
1.13
1.33
2.91
2.95
2.99
f1 (ppm)

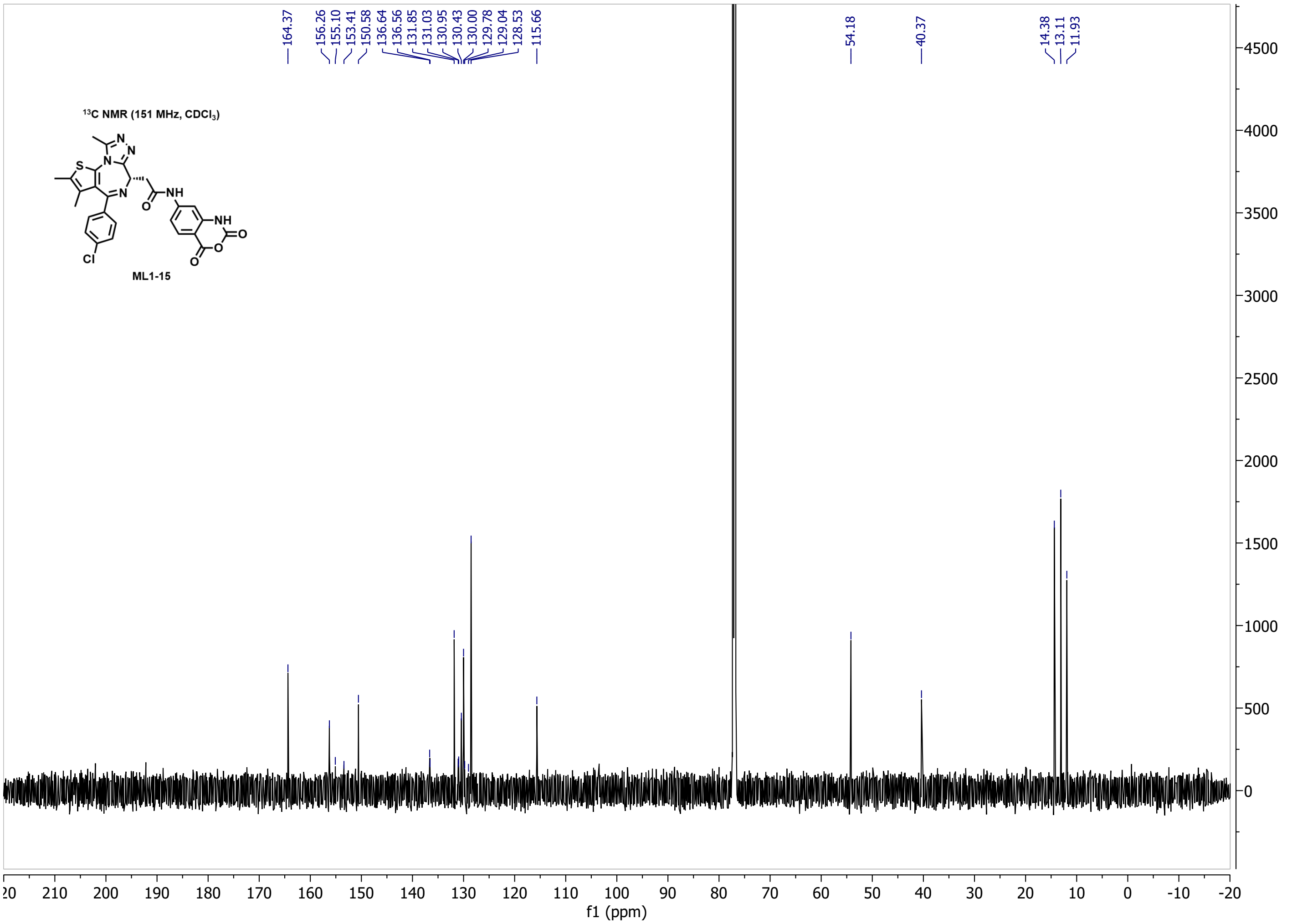
13C NMR (151 MHz, CDCl3)
ML1-15
NH
O
S
N
Cl
164.37
156.26
155.10
153.41
150.58
136.64
136.56
131.85
131.03
130.95
130.43
130.00
129.78
129.04
128.53
115.66
54.18
40.37
14.38
13.11
11.93
f1 (ppm)

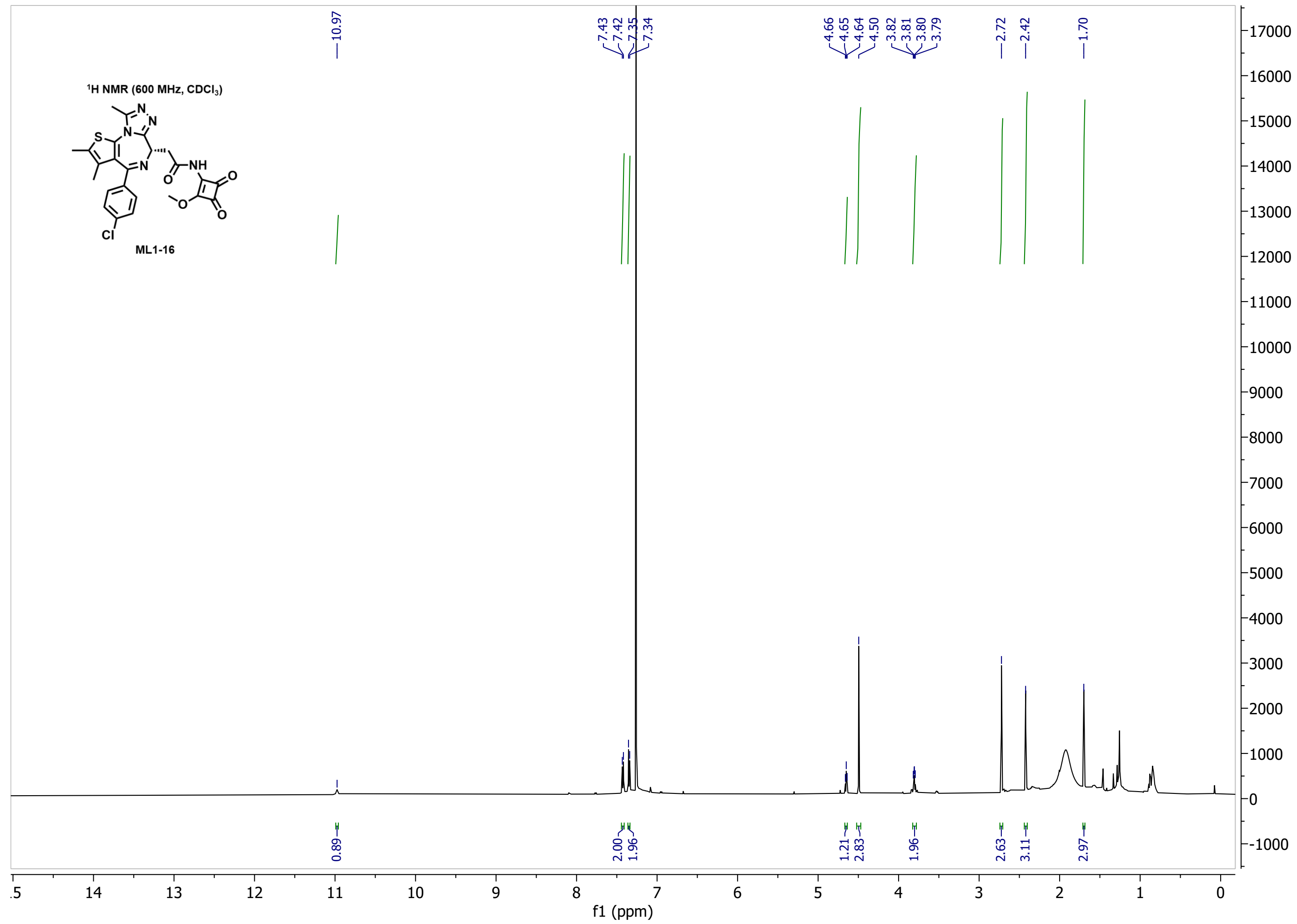

$^{1}$H NMR (600 MHz, CDCl$_3$)
ML1-16
10.97
7.43 7.42 7.35 7.34
4.66 4.65 4.64 4.50
3.82 3.81 3.80 3.79
2.72
2.42
1.70
0.89 2.00 1.96 1.21 2.83 1.96 2.63 3.11 2.97
f1 (ppm)


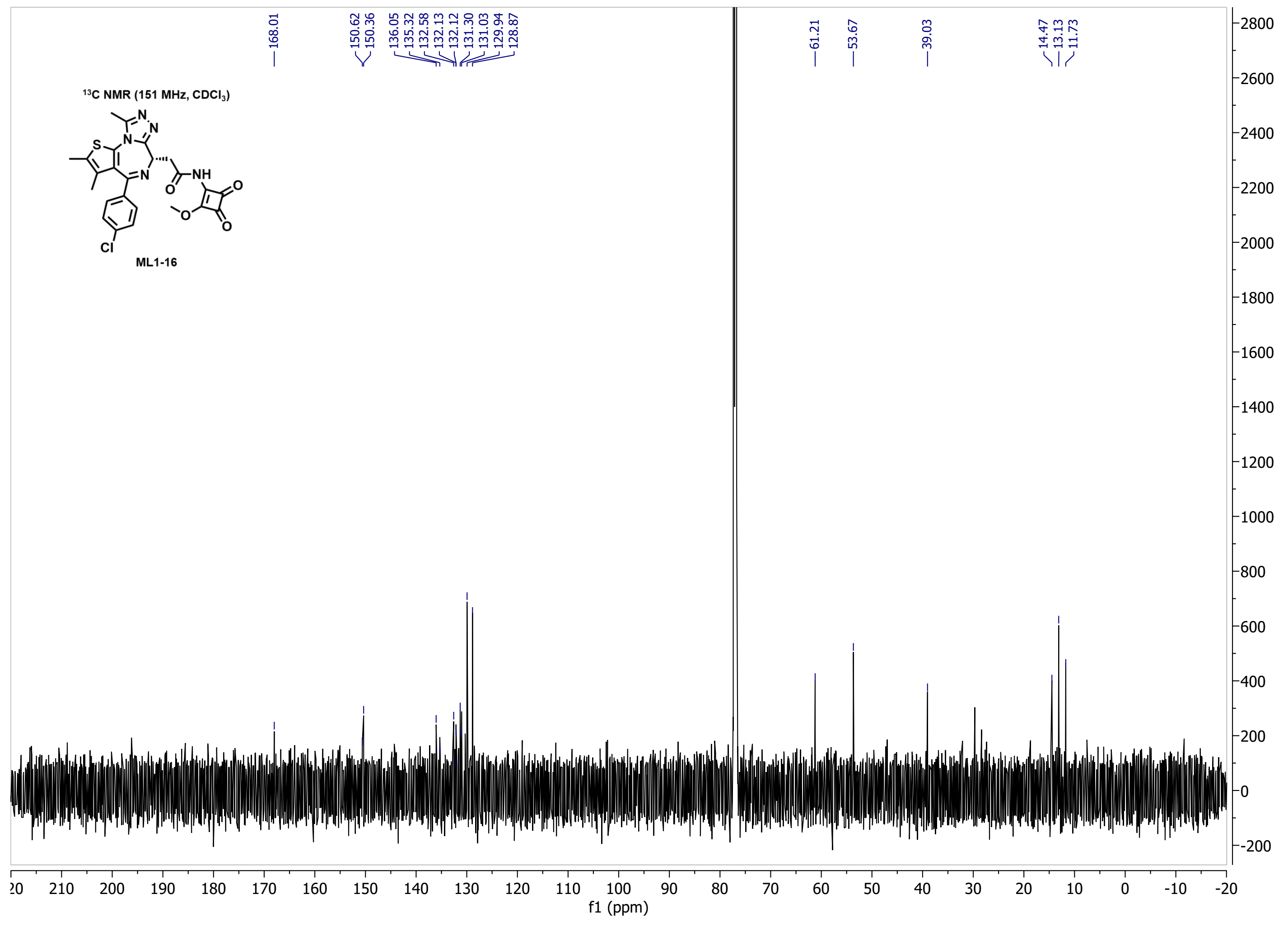

$^{13}$C NMR (151 MHz, CDCl$_3$)
ML1-16
168.01
150.62 150.36
136.05 135.32 132.58 132.13 132.12 131.30 131.03 129.94 128.87
61.21
53.67
39.03
14.47 13.13 11.73
f1 (ppm)

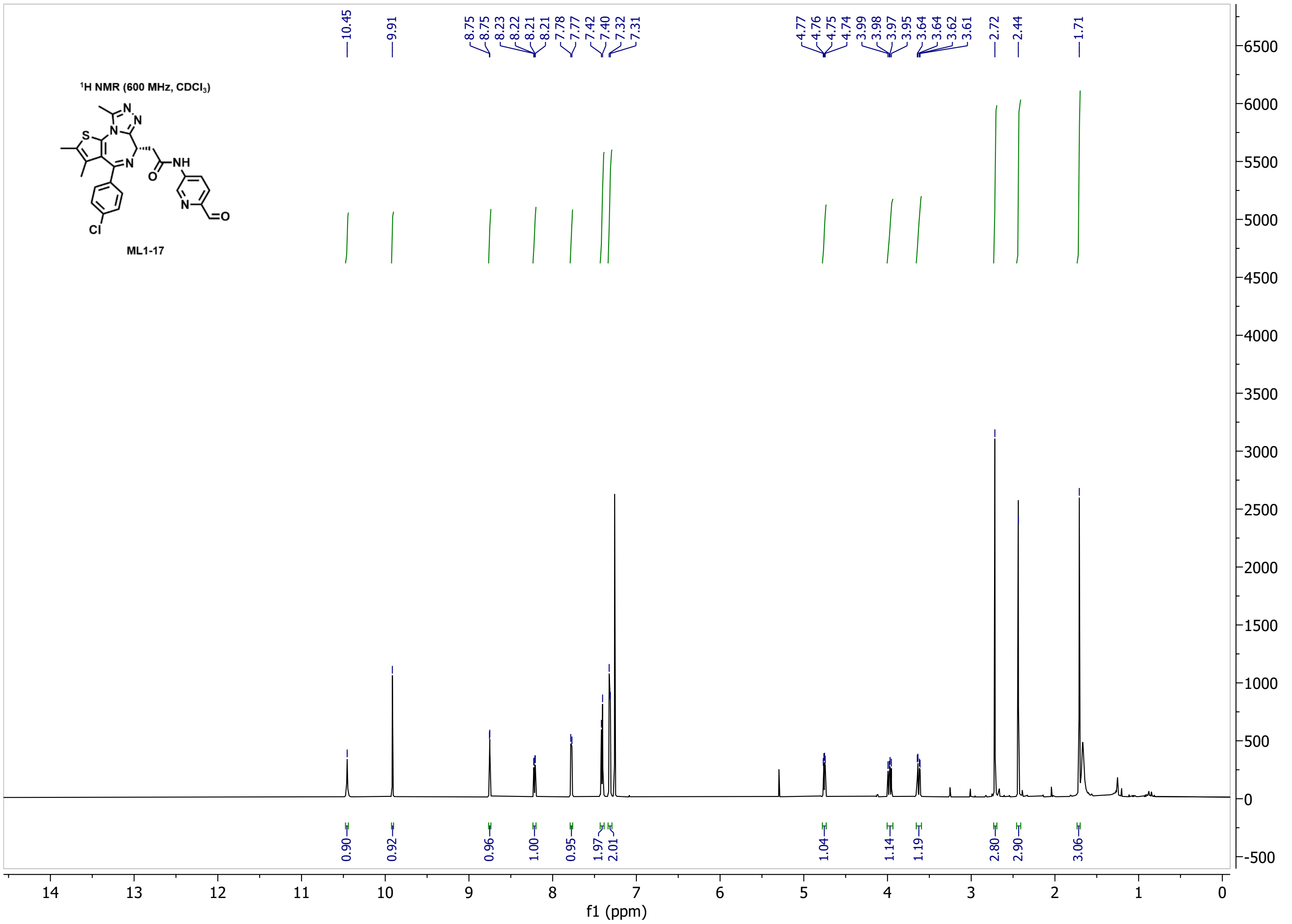
1H NMR (600 MHz, $CDCl_3$)
ML1-17
10.45
9.91
8.75
8.75
8.23
8.22
8.21
8.21
7.78
7.77
7.42
7.40
7.32
7.31
4.77
4.76
4.75
4.74
3.99
3.98
3.97
3.95
3.64
3.64
3.62
3.61
2.72
2.44
1.71
0.90
0.92
0.96
1.00
0.95
1.97
2.01
1.04
1.14
1.19
2.80
2.90
3.06
f1 (ppm)

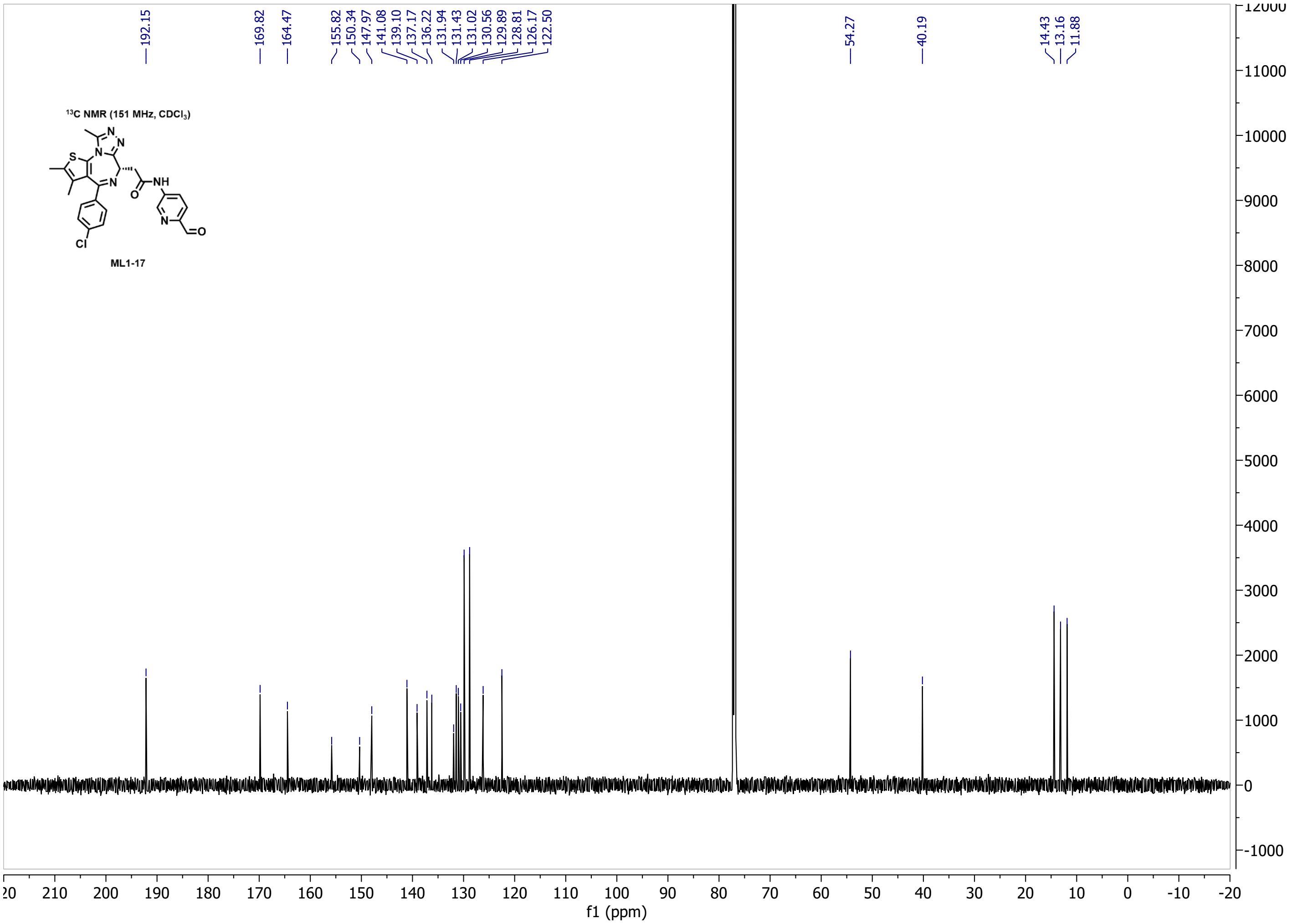
13C NMR (151 MHz, $CDCl_3$)
ML1-17
192.15
169.82
164.47
155.82
150.34
147.97
141.08
139.10
137.17
136.22
131.94
131.43
131.02
130.56
129.89
128.81
126.17
122.50
54.27
40.19
14.43
13.16
11.88
f1 (ppm)

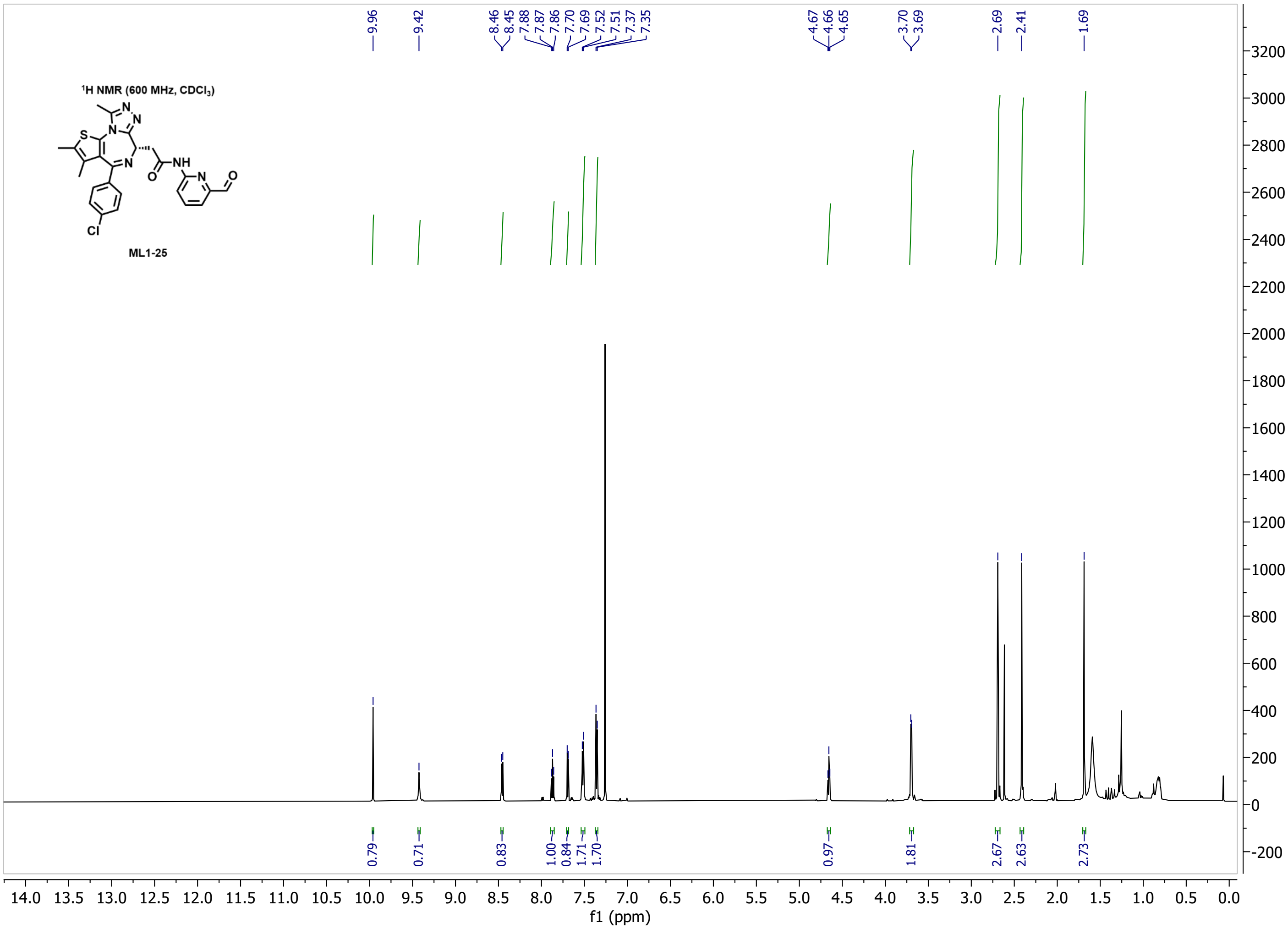

9.96
9.42
8.46 8.45 7.88 7.87 7.86 7.70 7.69 7.52 7.51 7.37 7.35
4.67 4.66 4.65
3.70 3.69
2.69
2.41
1.69
$^{1}$H NMR (600 MHz, CDCl$_3$)
ML1-25
0.79 0.71 0.83 1.00 0.84 1.71 1.70 0.97 1.81 2.67 2.63 2.73
f1 (ppm)


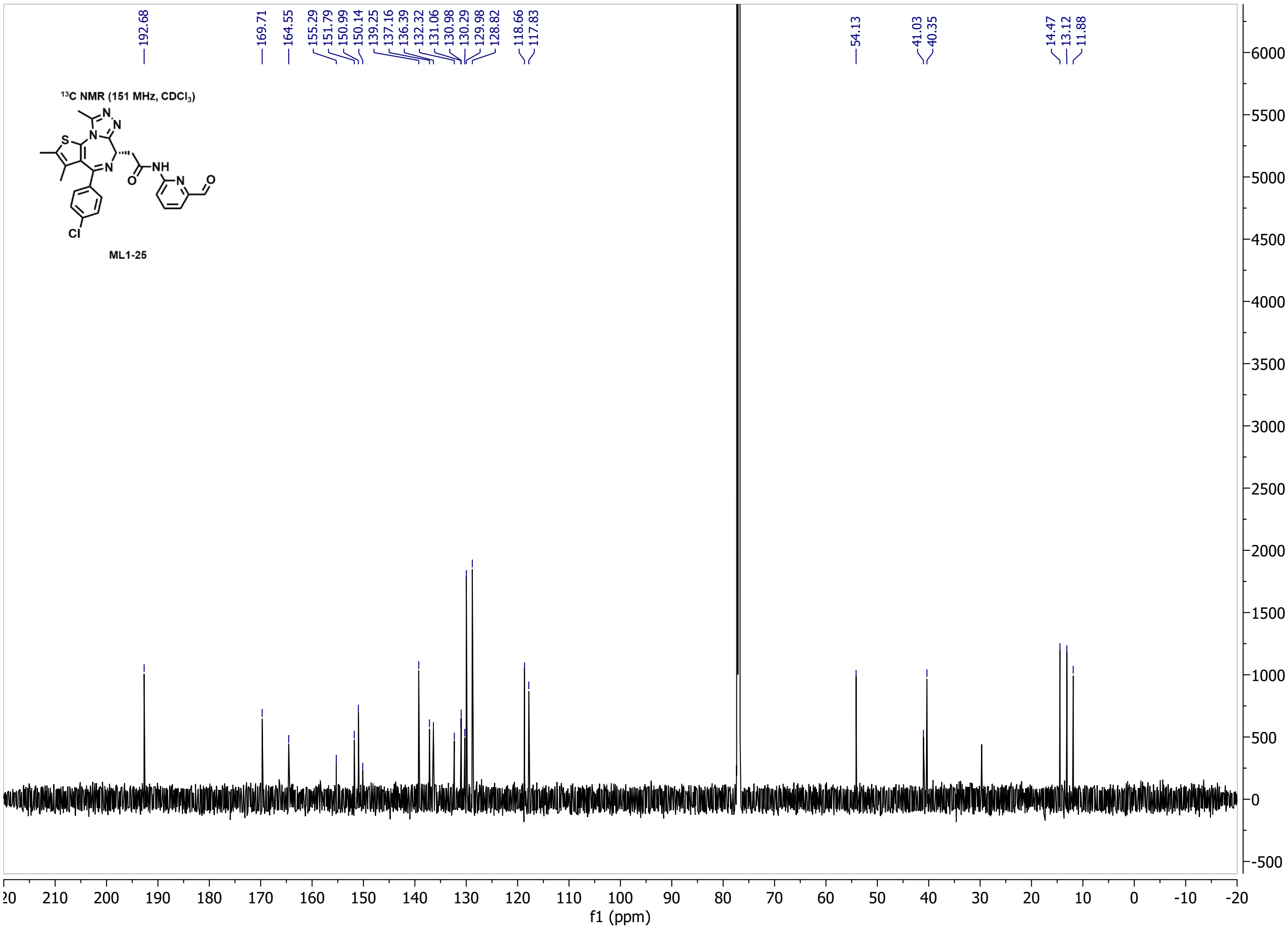

192.68
169.71 164.55 155.29 151.79 150.99 150.14 139.25 137.16 136.39 132.32 131.06 130.98 130.29 129.98 128.82 118.66 117.83
54.13
41.03 40.35
14.47 13.12 11.88
$^{13}$C NMR (151 MHz, CDCl$_3$)
ML1-25
f1 (ppm)

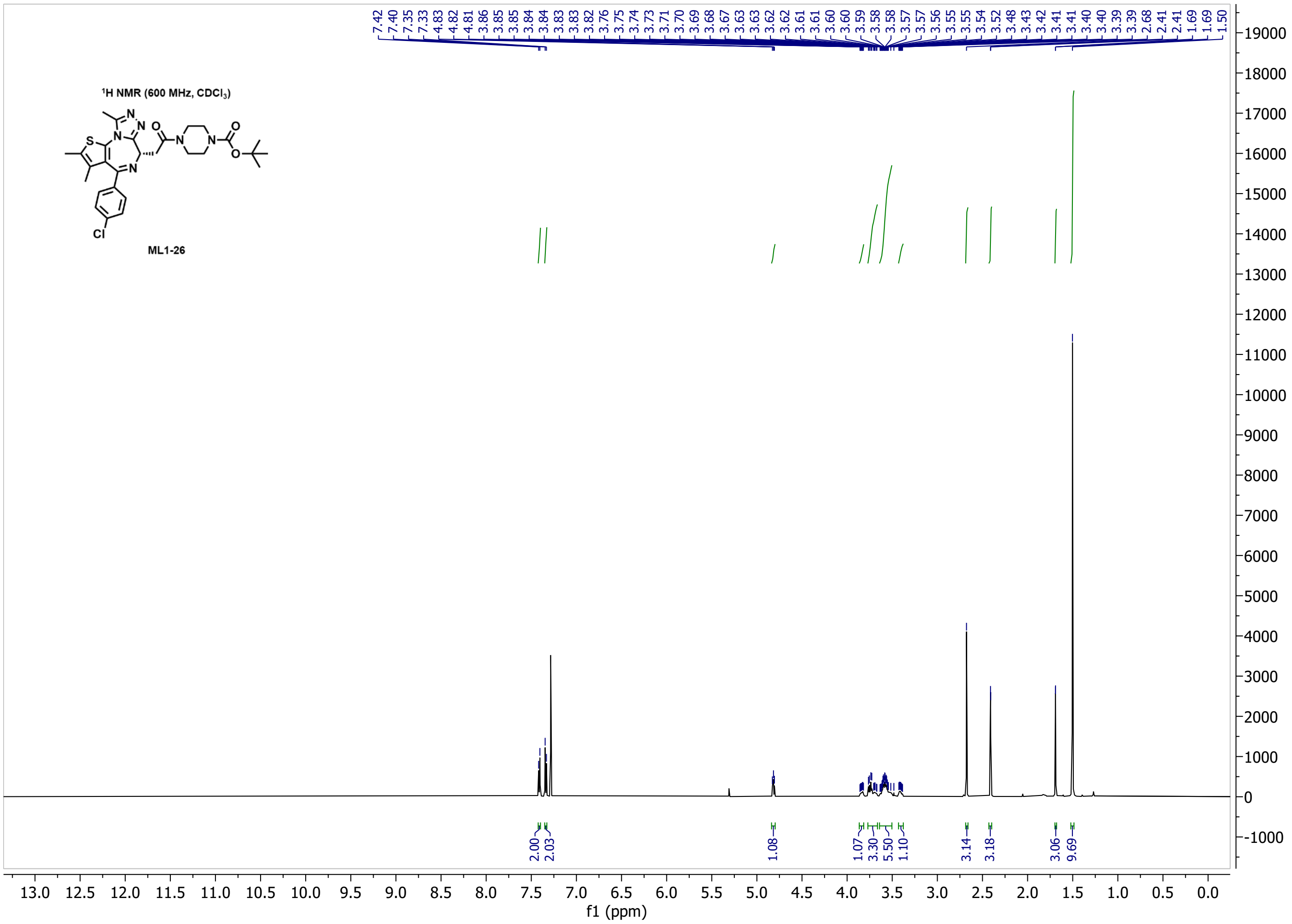
1H NMR (600 MHz, CDCl3)
ML1-26
7.42 7.40 7.35 7.33 4.83 4.82 4.81 3.86 3.85 3.85 3.84 3.84 3.83 3.83 3.82 3.76 3.75 3.74 3.73 3.71 3.70 3.69 3.68 3.67 3.63 3.63 3.62 3.62 3.61 3.61 3.60 3.60 3.59 3.58 3.58 3.57 3.57 3.56 3.55 3.55 3.54 3.52 3.48 3.43 3.42 3.41 3.41 3.40 3.40 3.39 3.39 2.68 2.41 2.41 1.69 1.69 1.50
2.00 2.03 1.08 1.07 3.30 5.50 1.10 3.14 3.18 3.06 9.69
f1 (ppm)

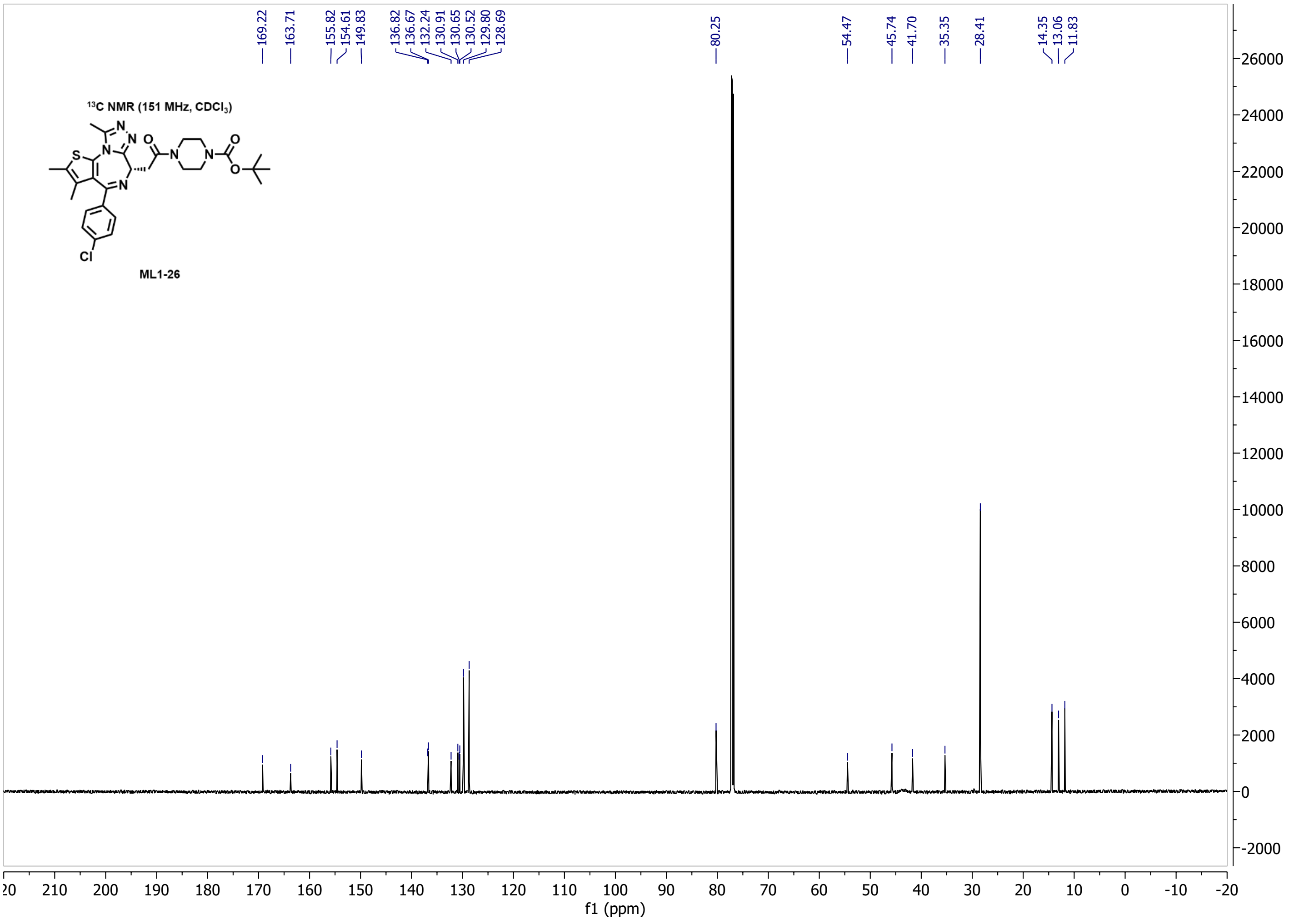
13C NMR (151 MHz, CDCl3)
ML1-26
169.22 163.71 155.82 154.61 149.83 136.82 136.67 132.24 130.91 130.65 130.52 129.80 128.69 80.25 54.47 45.74 41.70 35.35 28.41 14.35 13.06 11.83
f1 (ppm)

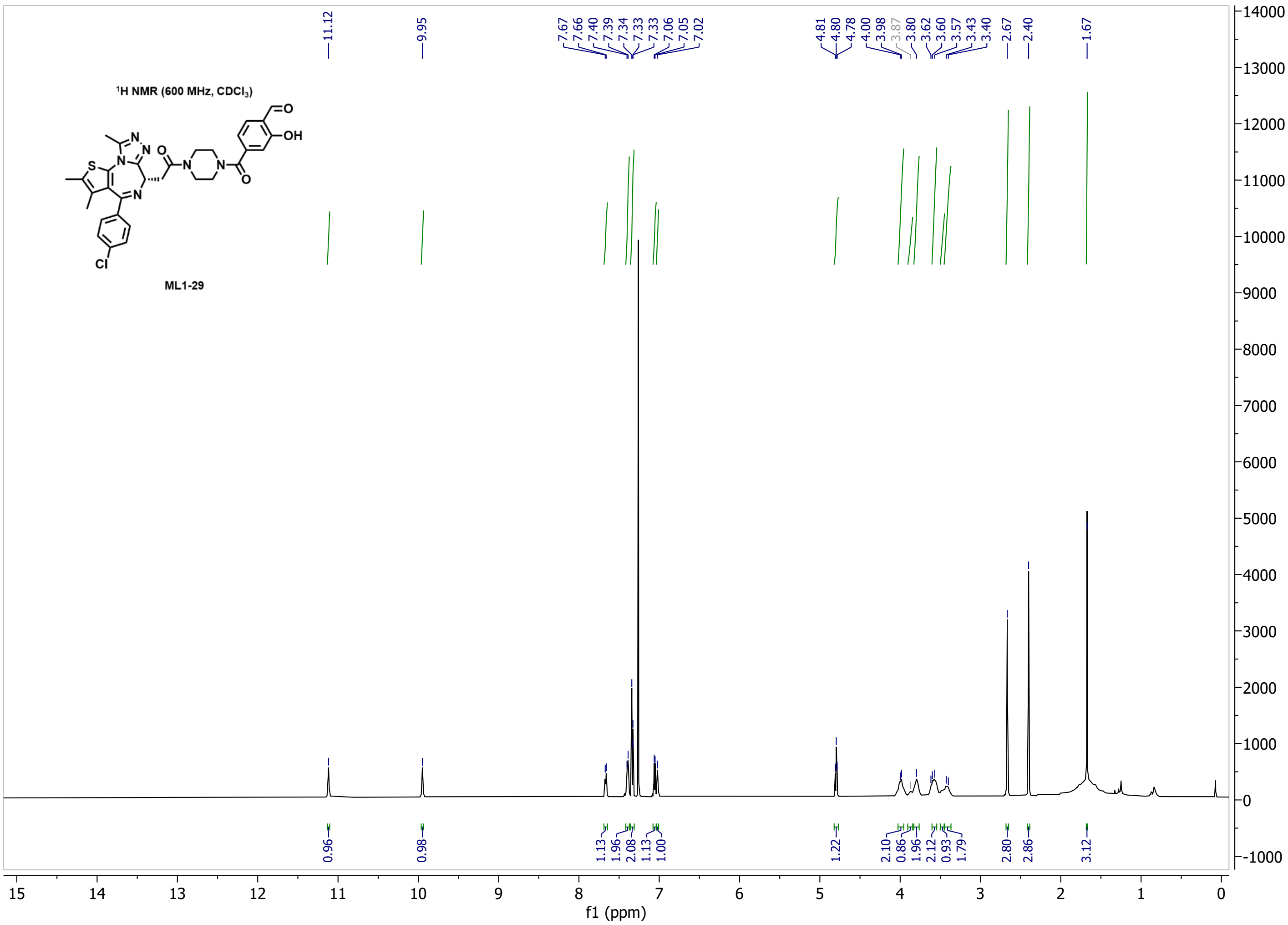

1H NMR (600 MHz, CDCl3)
ML1-29
11.12
9.95
7.67 7.66 7.40 7.39 7.34 7.33 7.33 7.06 7.05 7.02
4.81 4.80 4.78 4.00 3.98 3.87 3.80 3.62 3.60 3.57 3.43 3.40 2.67 2.40
1.67
0.96 0.98 1.13 1.96 2.08 1.13 1.00 1.22 2.10 0.86 1.96 2.12 0.93 1.79 2.80 2.86 3.12
f1 (ppm)


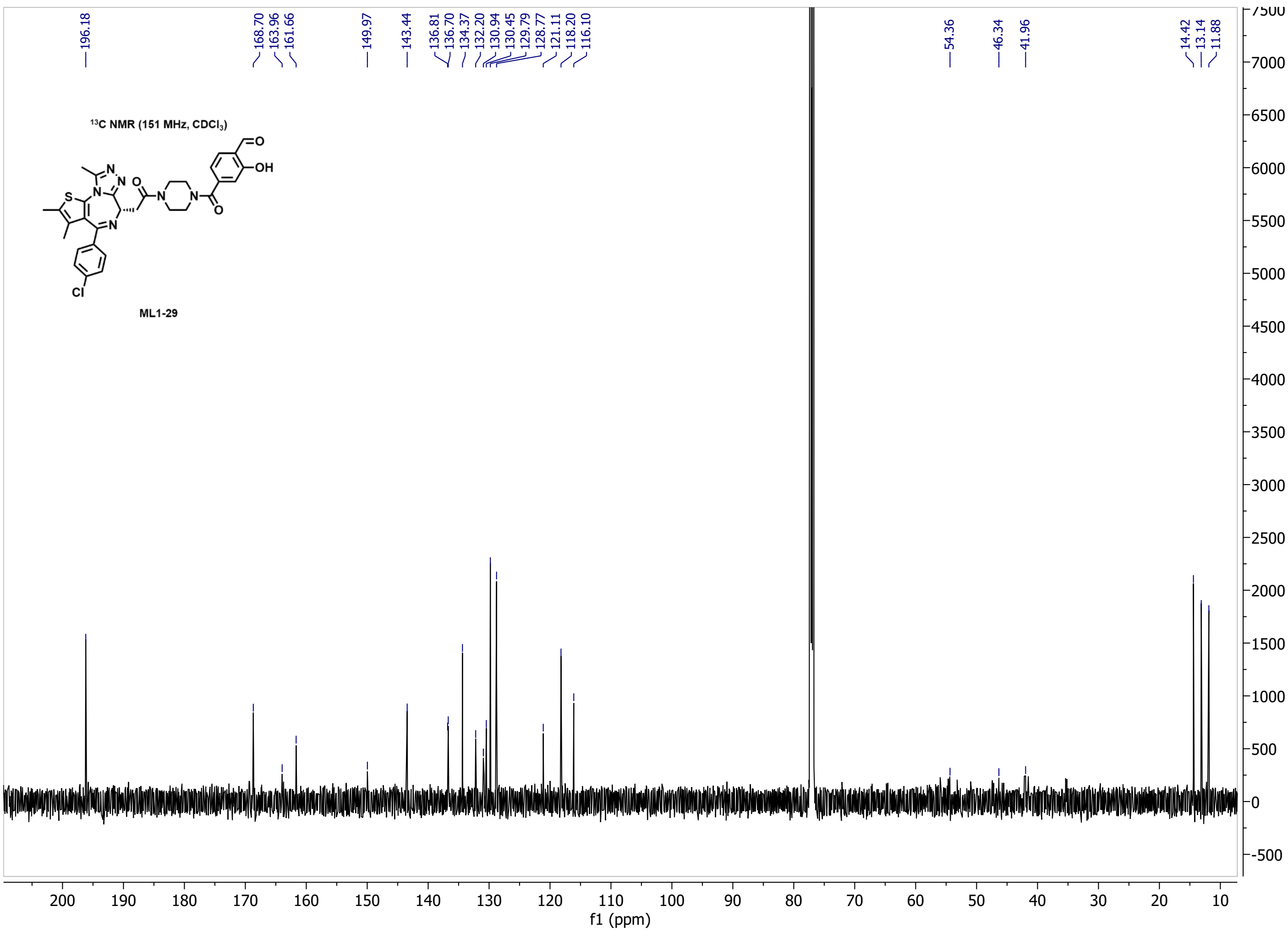

13C NMR (151 MHz, CDCl3)
ML1-29
196.18
168.70 163.96 161.66
149.97
143.44
136.81 136.70 134.37 132.20 130.94 130.45 129.79 128.77 121.11 118.20 116.10
54.36
46.34
41.96
14.42 13.14 11.88
f1 (ppm)

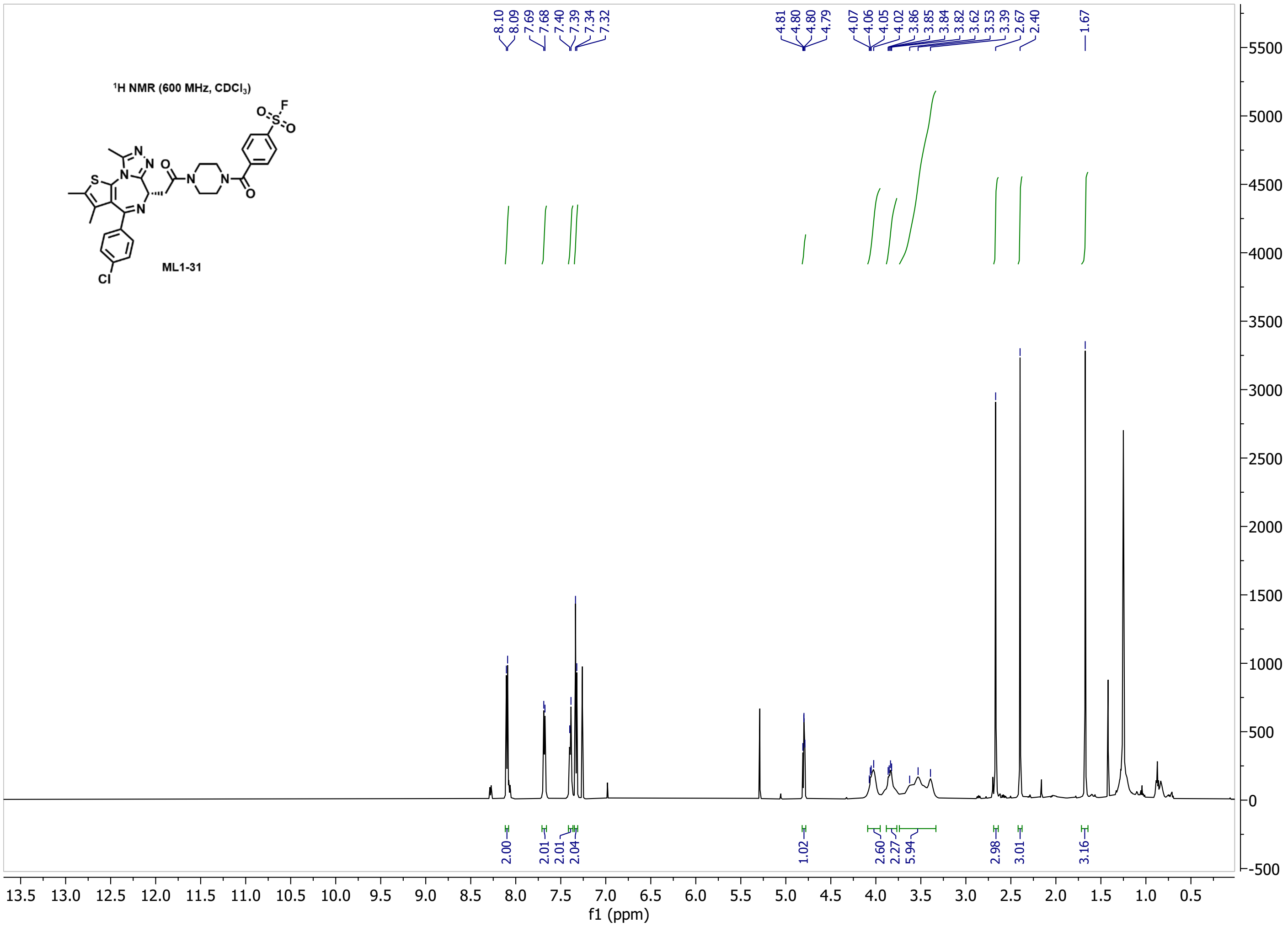

¹H NMR (600 MHz, CDCl₃)
ML1-31
8.10
8.09
7.69
7.68
7.40
7.39
7.34
7.32
4.81
4.80
4.80
4.79
4.07
4.06
4.05
4.02
3.86
3.85
3.84
3.82
3.62
3.53
3.39
2.67
2.40
1.67
2.00
2.01
2.01
2.04
1.02
2.60
2.27
5.94
2.98
3.01
3.16
f1 (ppm)


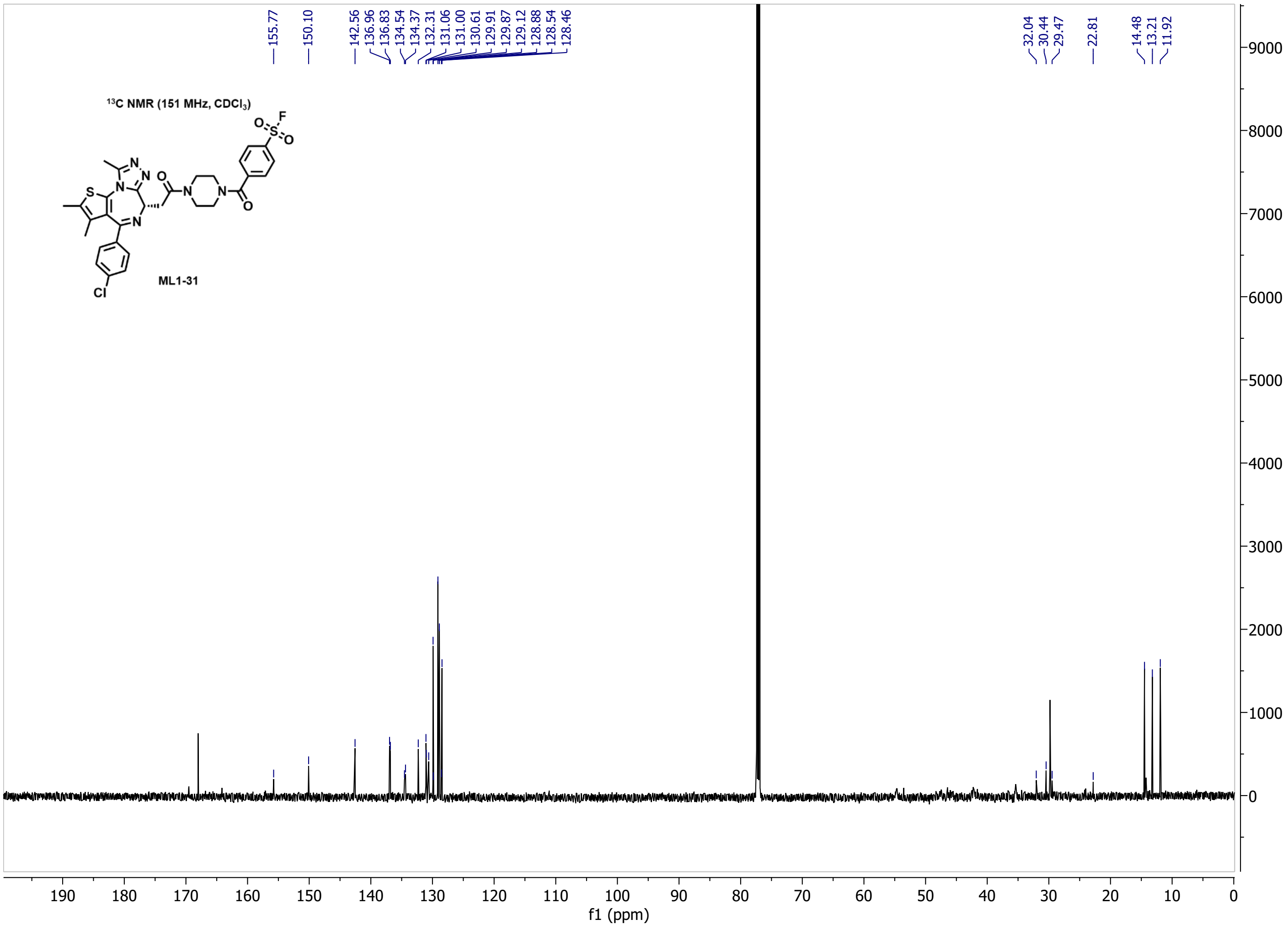

¹³C NMR (151 MHz, CDCl₃)
ML1-31
155.77
150.10
142.56
136.96
136.83
134.54
134.37
132.31
131.06
131.00
130.61
129.91
129.87
129.12
128.88
128.54
128.46
32.04
30.44
29.47
22.81
14.48
13.21
11.92
f1 (ppm)

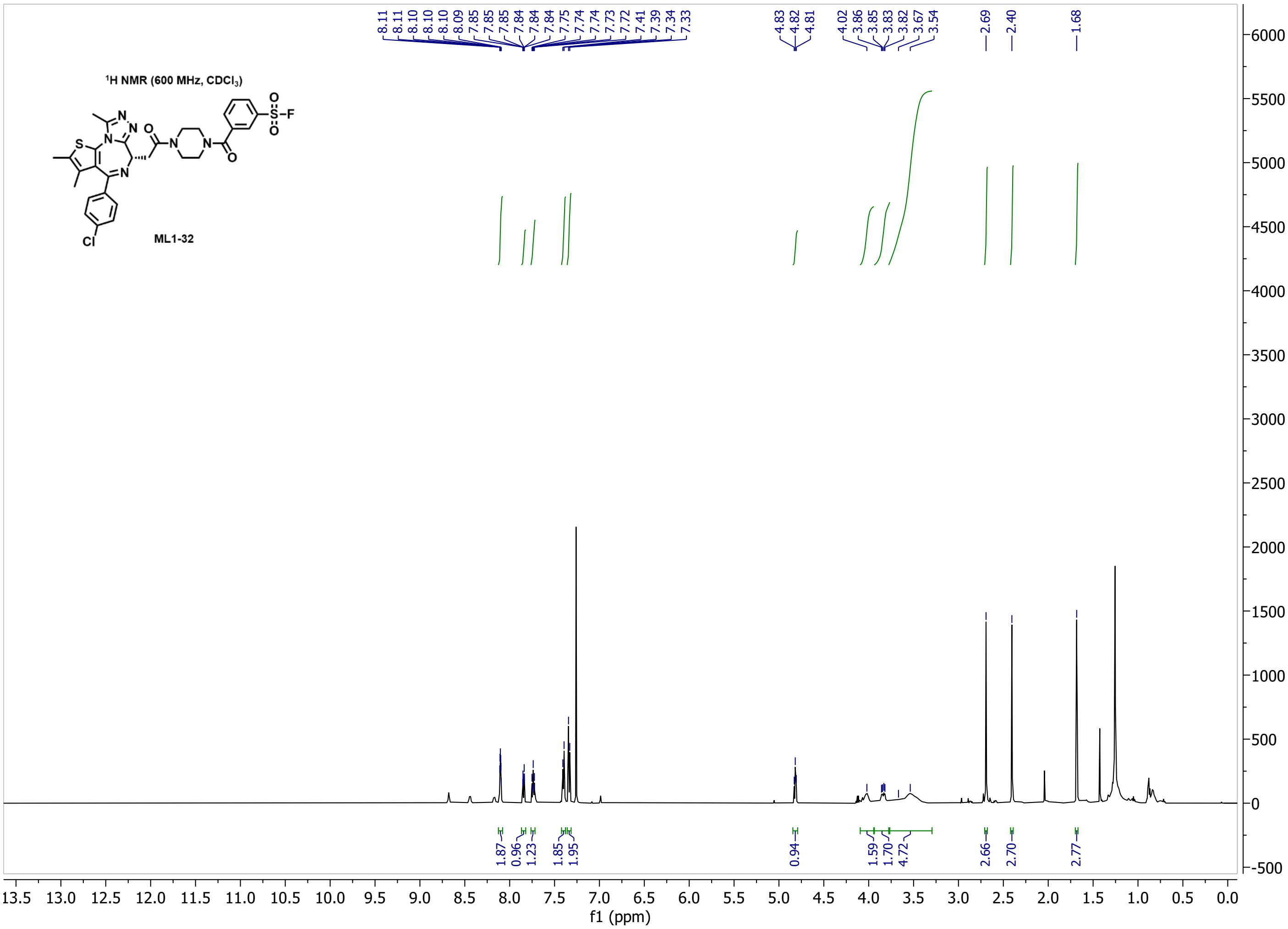

1H NMR (600 MHz, CDCl3)
ML1-32
8.11 8.11 8.10 8.10 8.10 8.09 7.85 7.85 7.85 7.84 7.84 7.84 7.75 7.74 7.74 7.73 7.72 7.41 7.39 7.34 7.33
4.83 4.82 4.81
4.02 3.86 3.85 3.83 3.82 3.67 3.54
2.69 2.40 1.68
1.87 0.96 1.23 1.85 1.95 0.94 1.59 1.70 4.72 2.66 2.70 2.77
f1 (ppm)


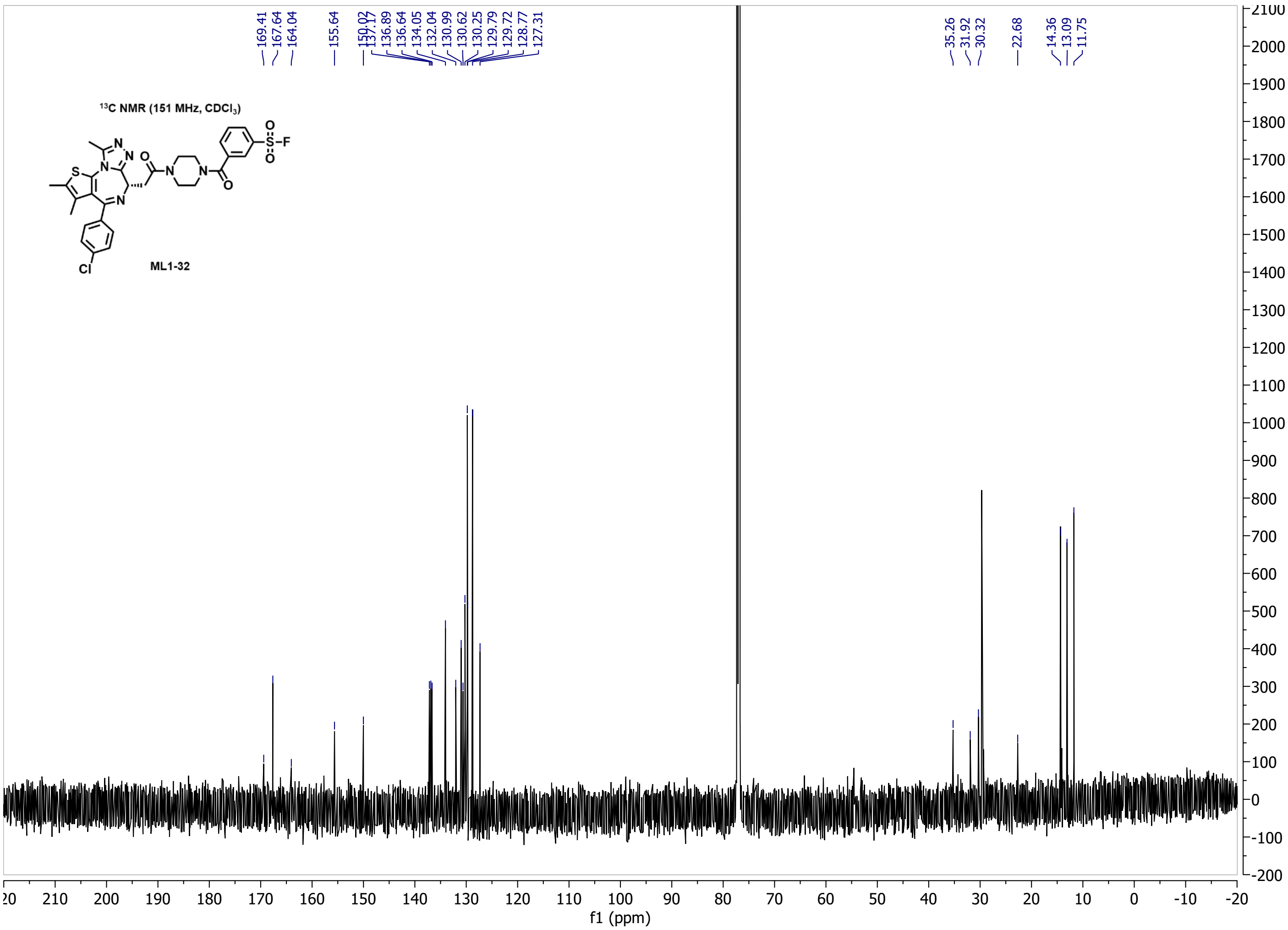

13C NMR (151 MHz, CDCl3)
ML1-32
169.41 167.64 164.04 155.64 150.02 137.17 136.89 136.64 134.05 132.04 130.99 130.62 130.25 129.79 129.72 128.77 127.31
35.26 31.92 30.32 22.68 14.36 13.09 11.75
f1 (ppm)

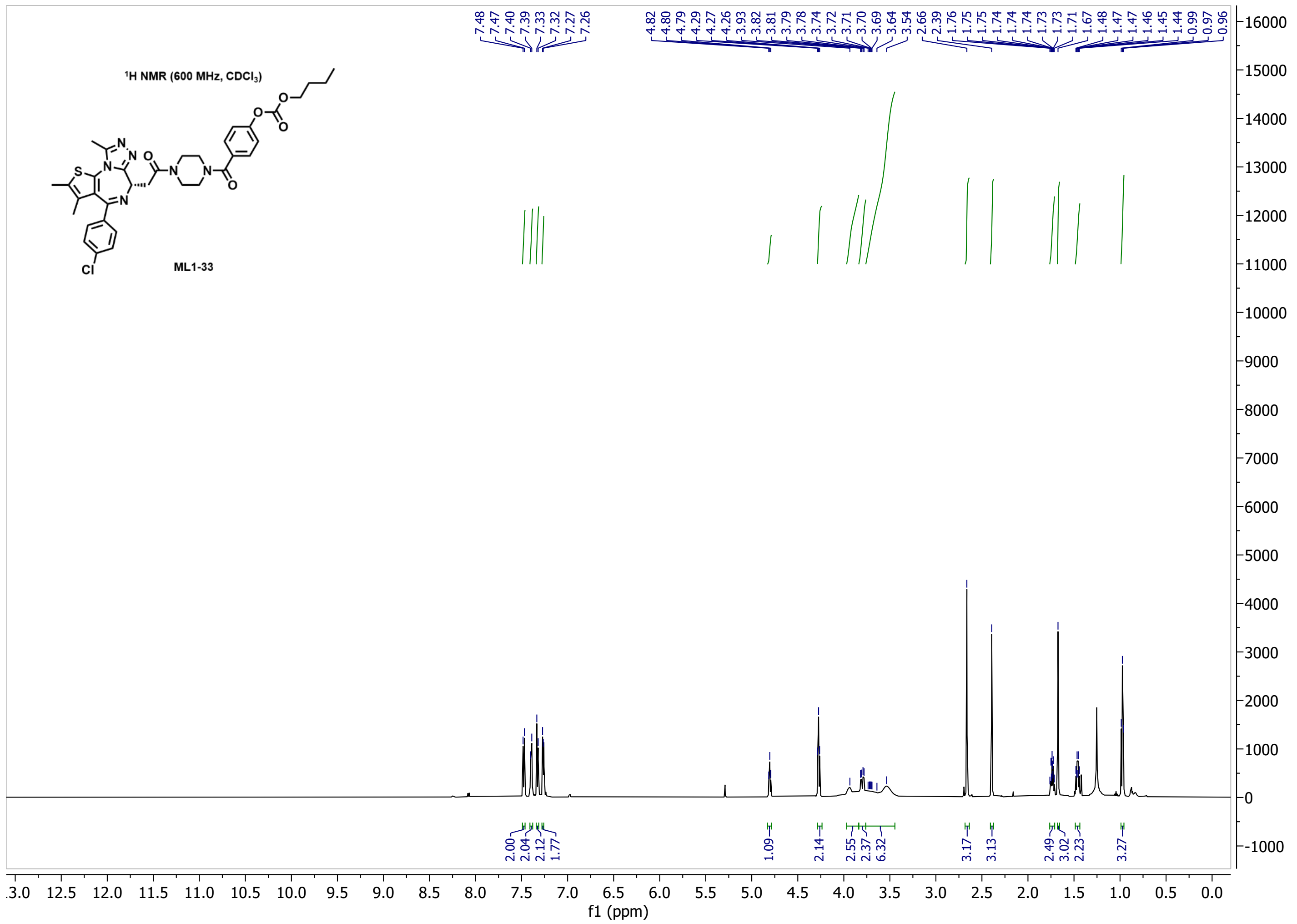

¹H NMR (600 MHz, CDCl₃)
ML1-33
f1 (ppm)


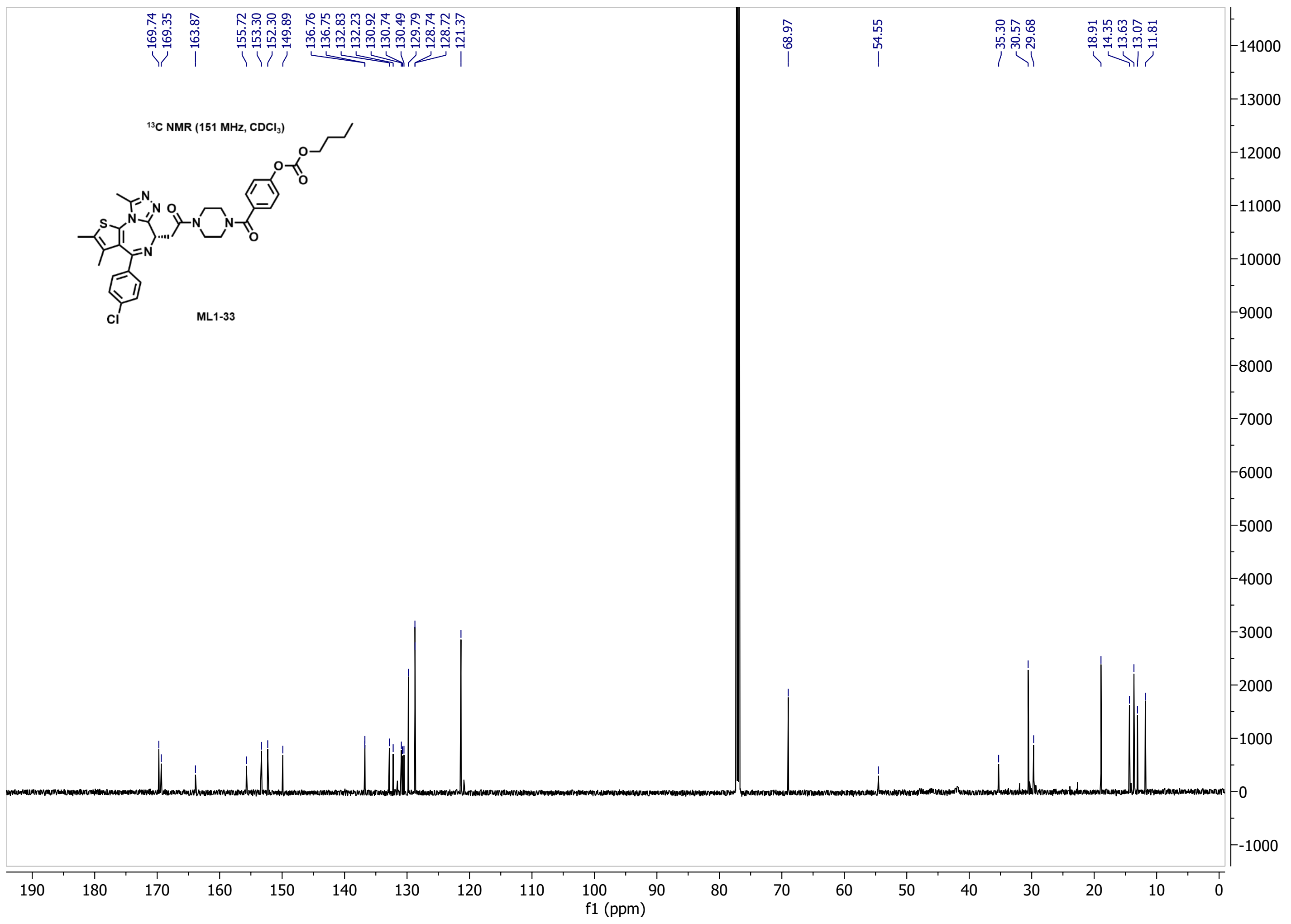

¹³C NMR (151 MHz, CDCl₃)
ML1-33
f1 (ppm)

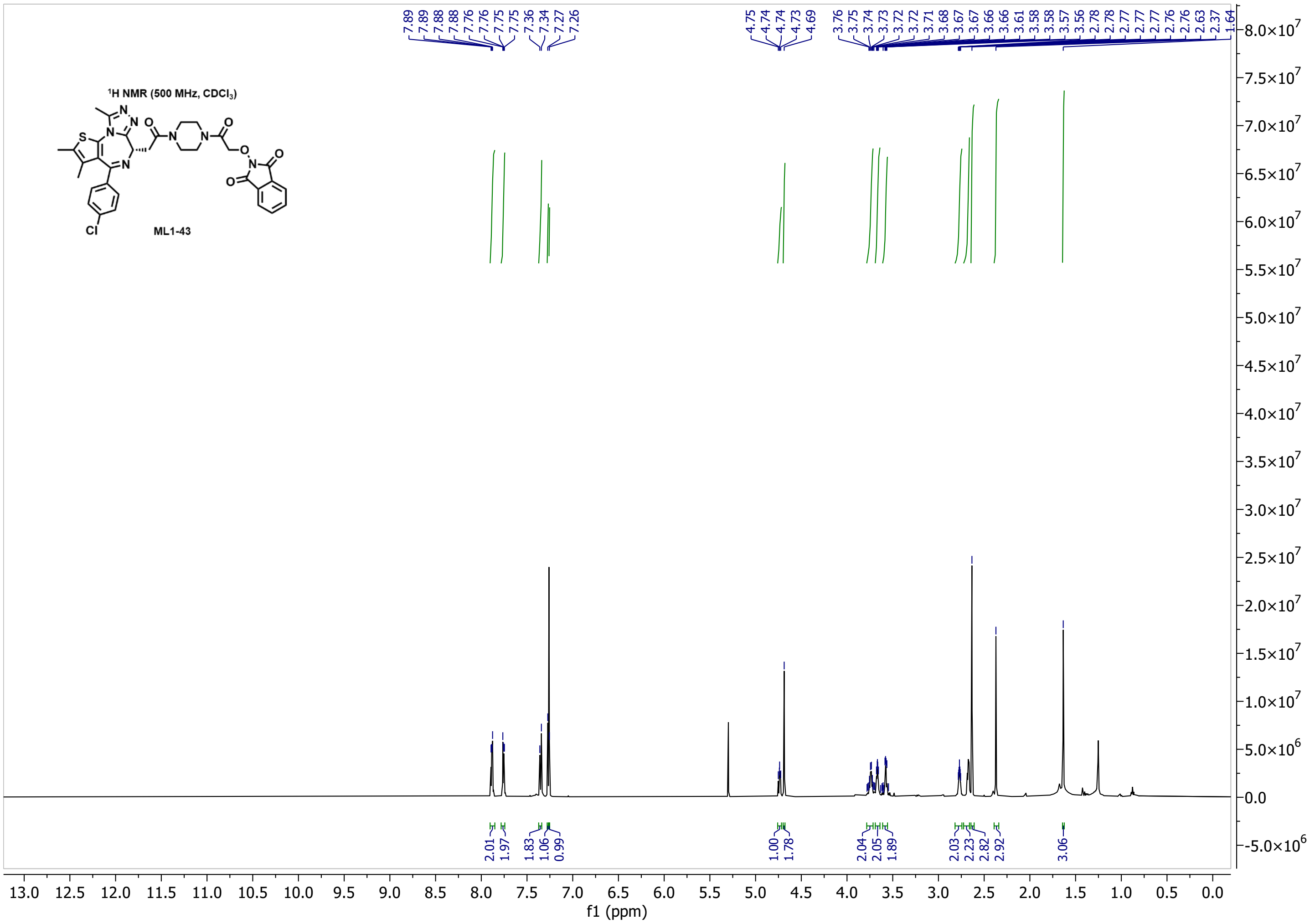

1H NMR (500 MHz, CDCl3)
ML1-43
7.89 7.89 7.88 7.76 7.76 7.75 7.75 7.36 7.34 7.27 7.26
4.75 4.74 4.74 4.73 4.69
3.76 3.75 3.74 3.73 3.72 3.72 3.71 3.68 3.67 3.67 3.66 3.66 3.61 3.58 3.58 3.57 3.56 2.78 2.78 2.77 2.77 2.77 2.76 2.76 2.63 2.37 1.64
2.01 1.97 1.83 1.06 0.99 1.00 1.78 2.04 2.05 1.89 2.03 2.23 2.82 2.92 3.06
f1 (ppm)


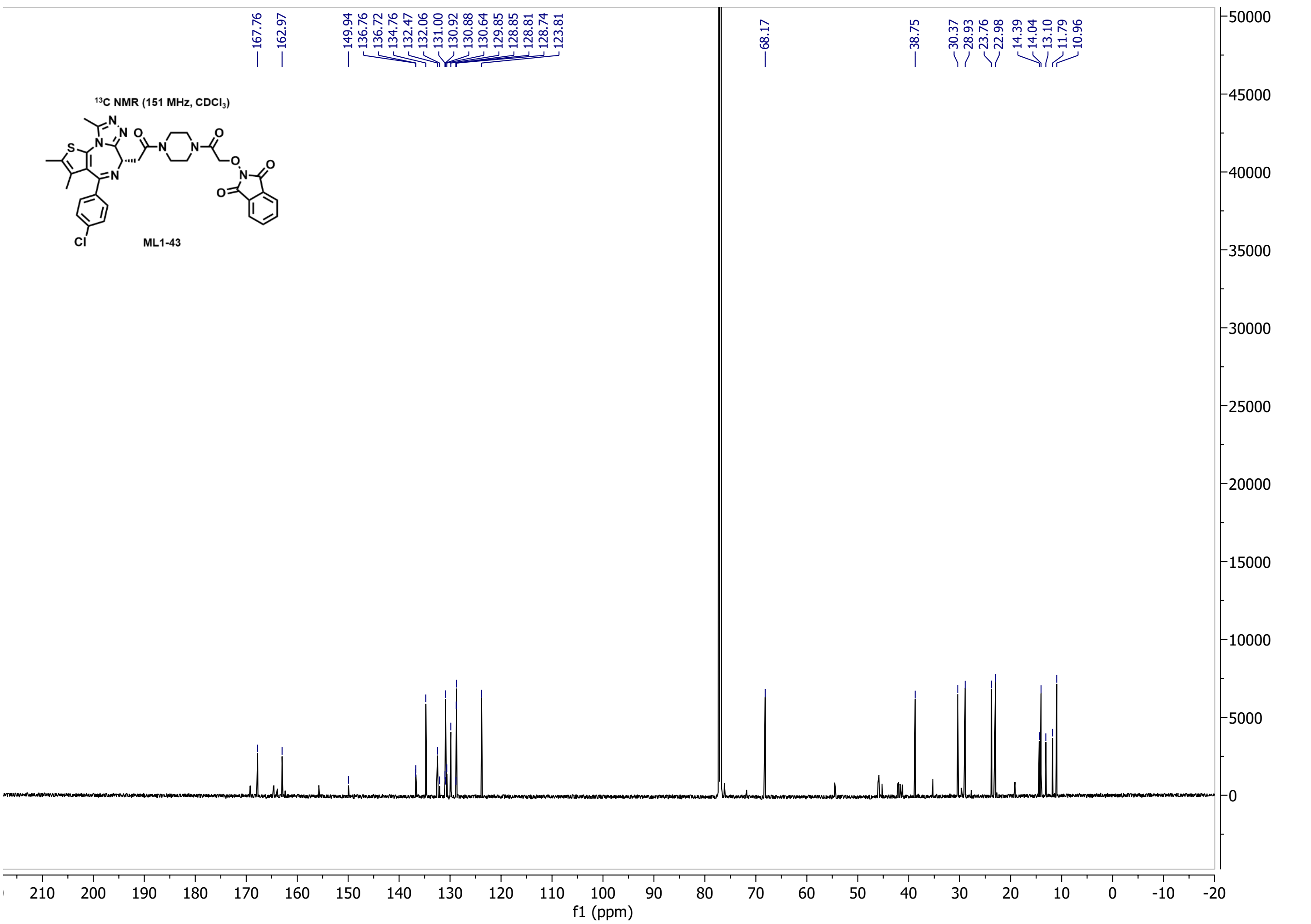

13C NMR (151 MHz, CDCl3)
ML1-43
167.76 162.97 149.94 136.76 136.72 134.76 132.47 132.06 131.00 130.92 130.88 130.64 129.85 128.85 128.81 128.74 123.81
68.17 38.75 30.37 28.93 23.76 22.98 14.39 14.04 13.10 11.79 10.96
f1 (ppm)

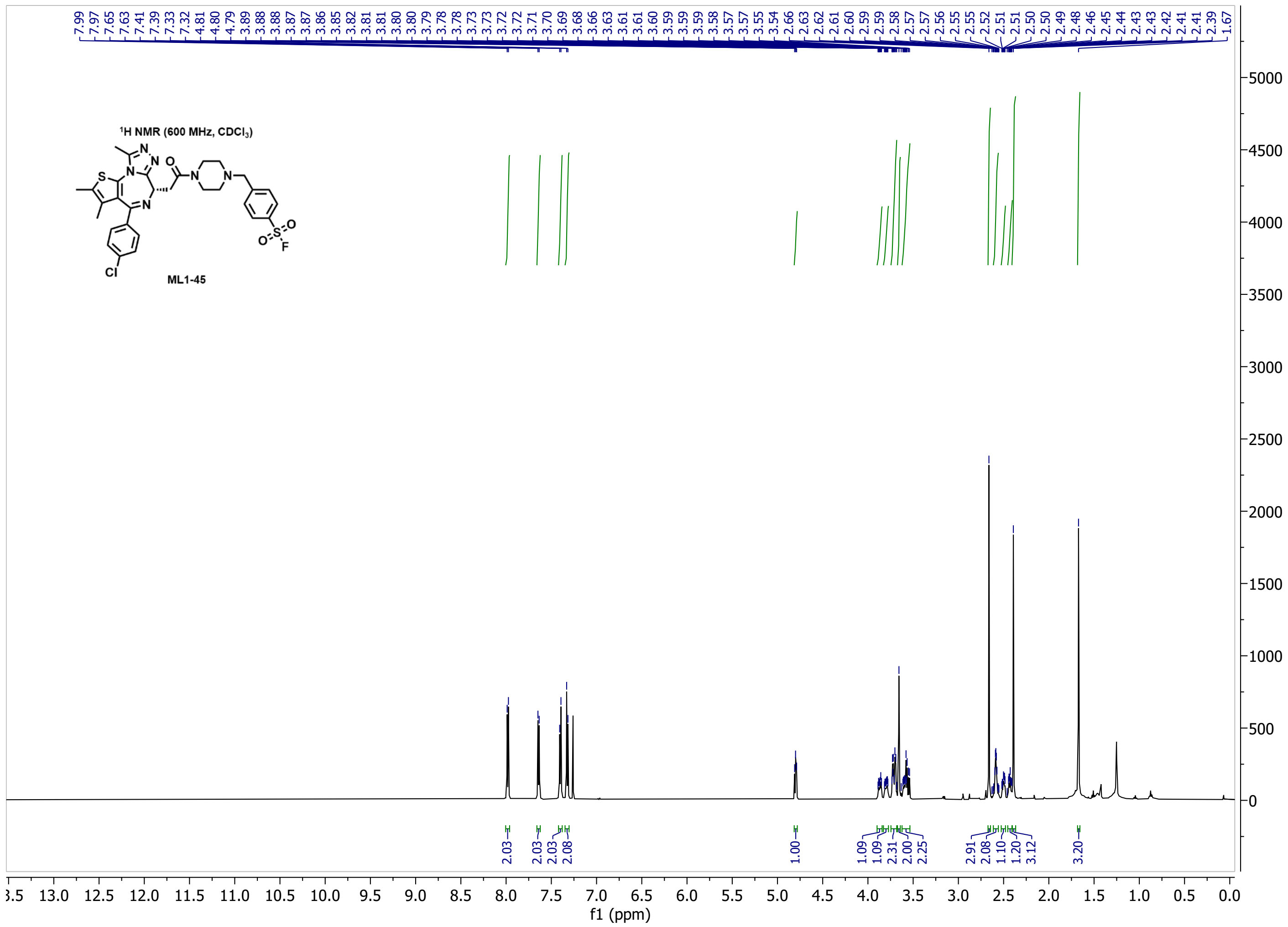

7.99 7.97 7.65 7.63 7.41 7.39 7.33 7.32 4.81 4.80 4.79 3.89 3.88 3.88 3.87 3.87 3.86 3.85 3.82 3.81 3.81 3.80 3.80 3.79 3.78 3.78 3.73 3.73 3.72 3.72 3.71 3.70 3.69 3.68 3.66 3.63 3.61 3.61 3.60 3.59 3.59 3.59 3.58 3.57 3.57 3.55 3.54 2.66 2.63 2.62 2.61 2.60 2.59 2.59 2.58 2.57 2.57 2.56 2.55 2.55 2.52 2.51 2.51 2.50 2.50 2.49 2.48 2.46 2.45 2.44 2.43 2.43 2.42 2.41 2.41 2.39 1.67
¹H NMR (600 MHz, CDCl3)
ML1-45
2.03 2.03 2.03 2.08 1.00 1.09 1.09 2.31 2.00 2.25 2.91 2.08 1.10 1.20 3.12 3.20
f1 (ppm)


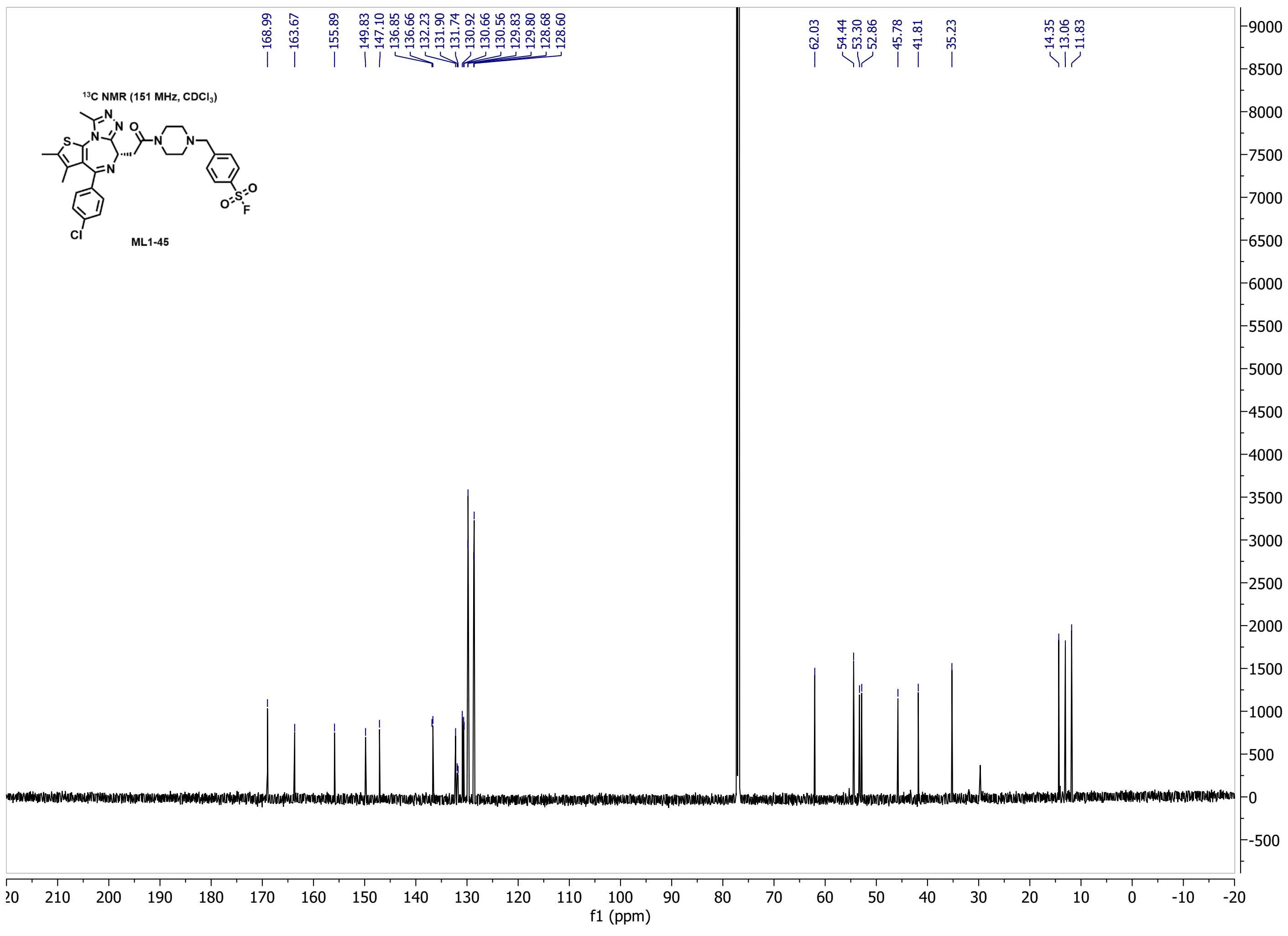

168.99 163.67 155.89 149.83 147.10 136.85 136.66 132.23 131.90 131.74 130.92 130.66 130.56 129.83 129.80 128.68 128.60 62.03 54.44 53.30 52.86 45.78 41.81 35.23 14.35 13.06 11.83
¹³C NMR (151 MHz, CDCl3)
ML1-45
f1 (ppm)

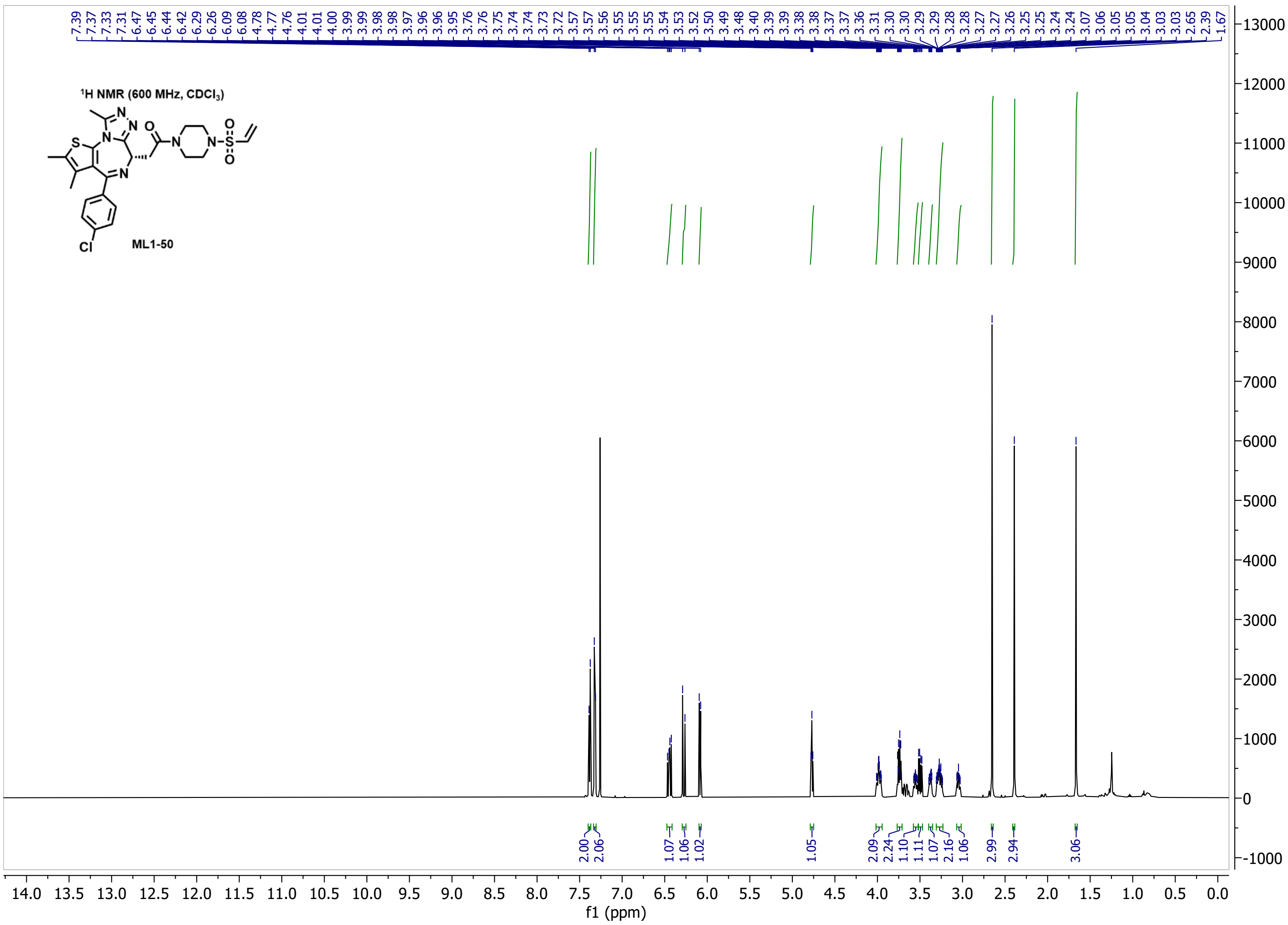

$^1$H NMR (600 MHz, $CDCl_3$)
ML1-50


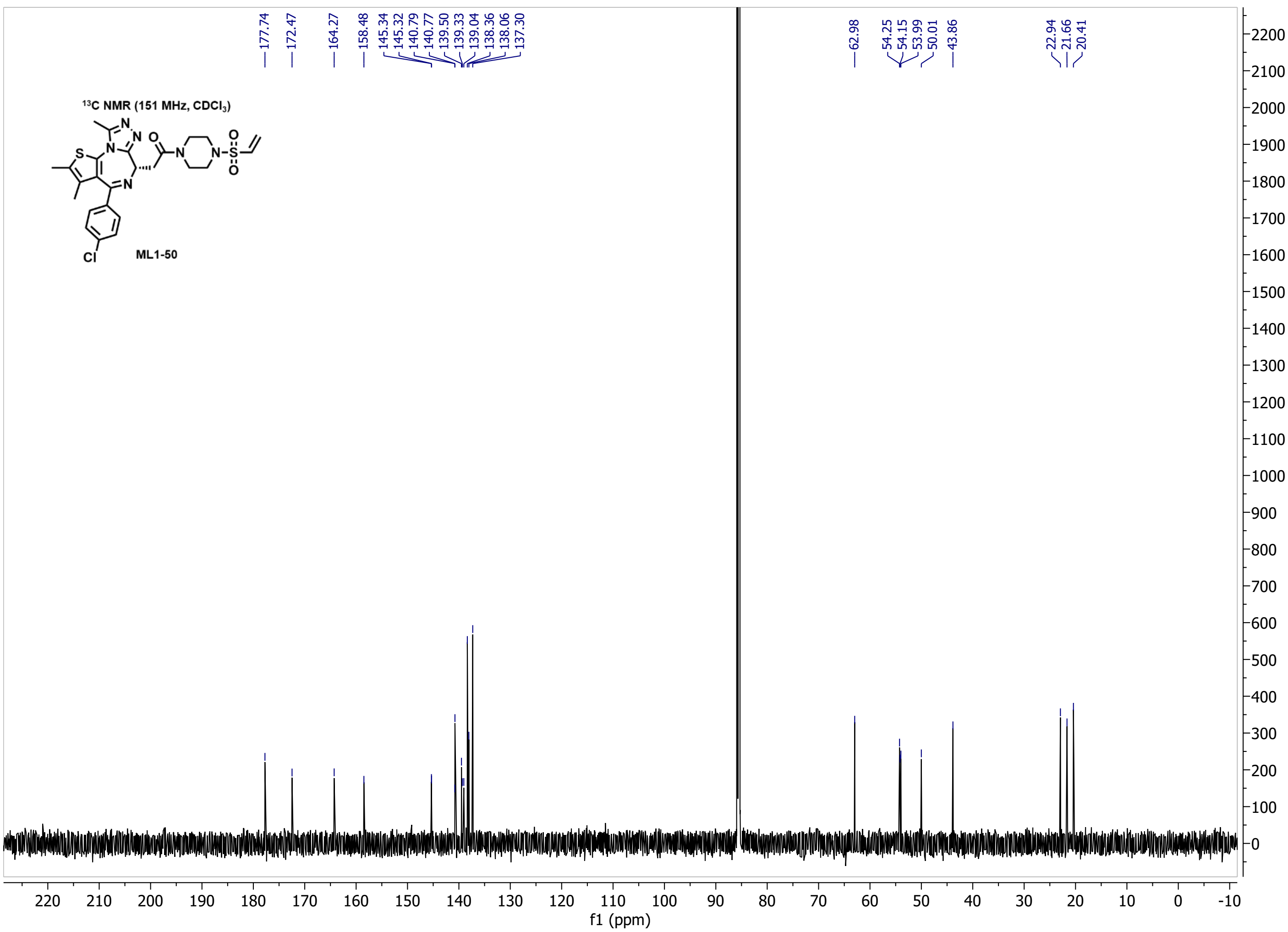

$^{13}$C NMR (151 MHz, $CDCl_3$)
ML1-50

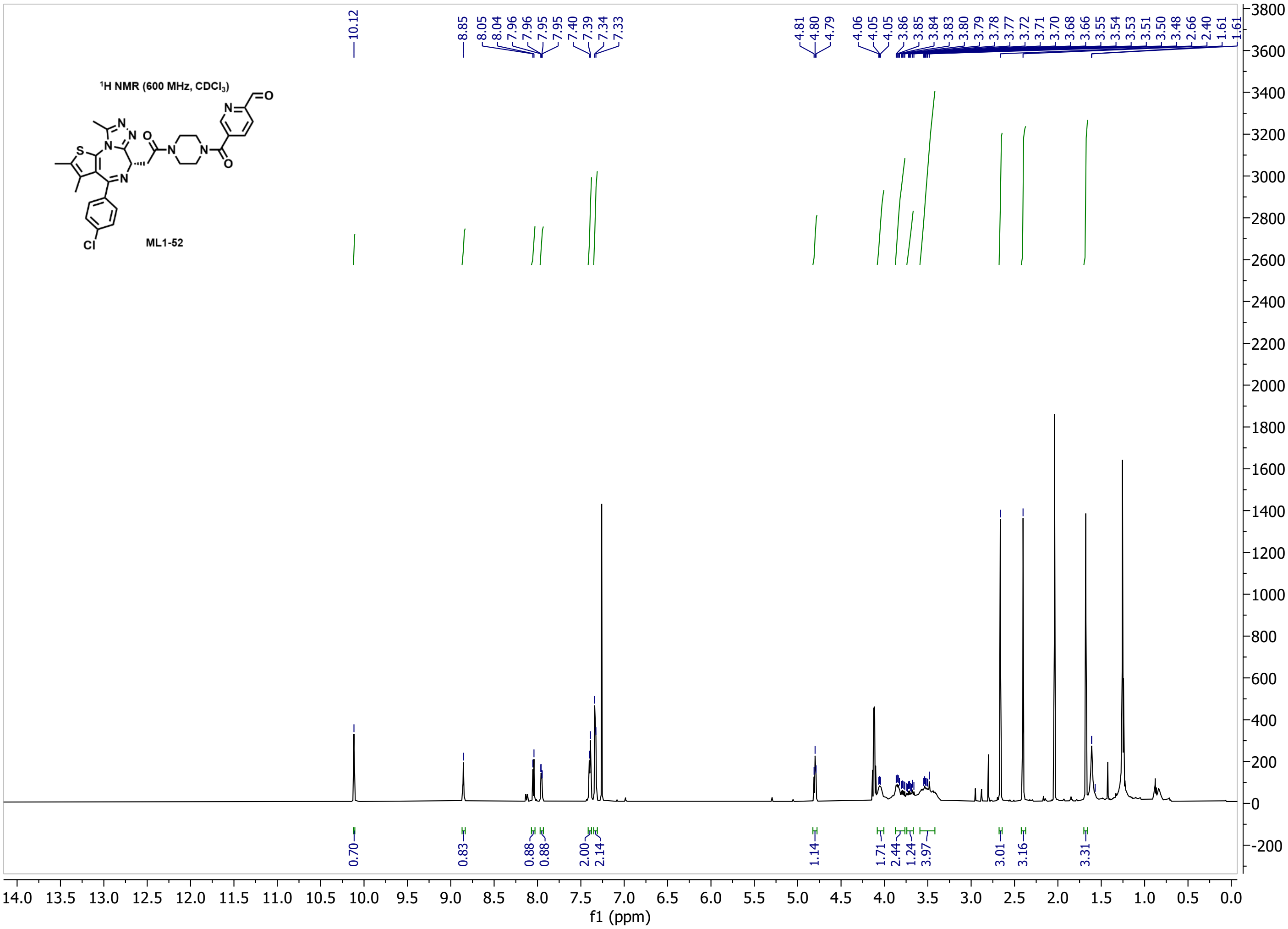

1H NMR (600 MHz, CDCl3)
ML1-52
10.12
8.85 8.05 8.04 7.96 7.95 7.95 7.40 7.39 7.34 7.33
4.81 4.80 4.79 4.06 4.05 4.05 3.86 3.85 3.84 3.83 3.80 3.79 3.78 3.77 3.72 3.71 3.70 3.68 3.66 3.55 3.54 3.53 3.51 3.50 3.48 2.66 2.40 1.61 1.61
0.70 0.83 0.88 0.88 2.00 2.14 1.14 1.71 2.44 1.24 3.97 3.01 3.16 3.31
f1 (ppm)


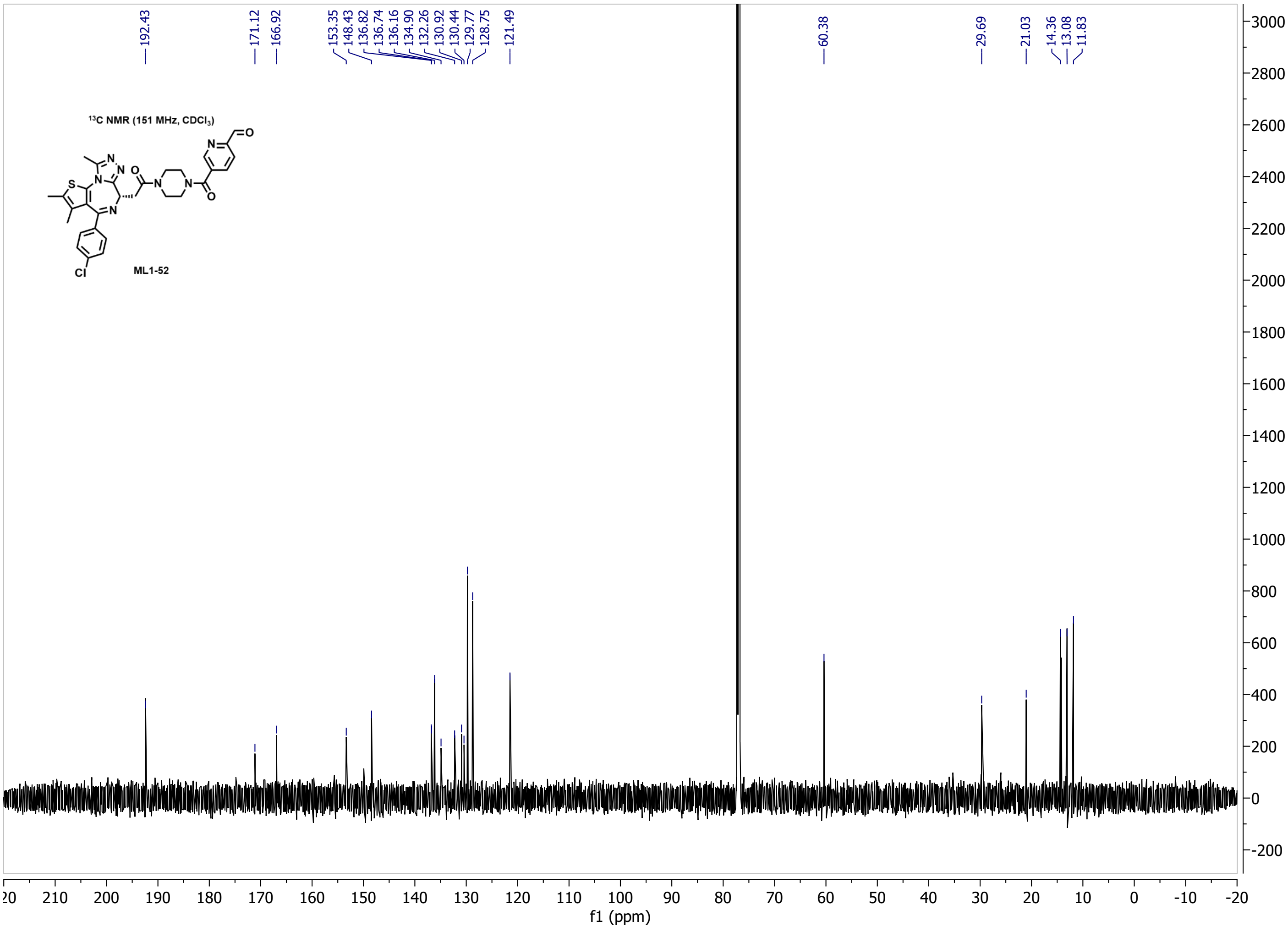

13C NMR (151 MHz, CDCl3)
ML1-52
192.43
171.12 166.92
153.35 148.43 136.82 136.74 136.16 134.90 132.26 130.92 130.44 129.77 128.75 121.49
60.38
29.69 21.03 14.36 13.08 11.83
f1 (ppm)

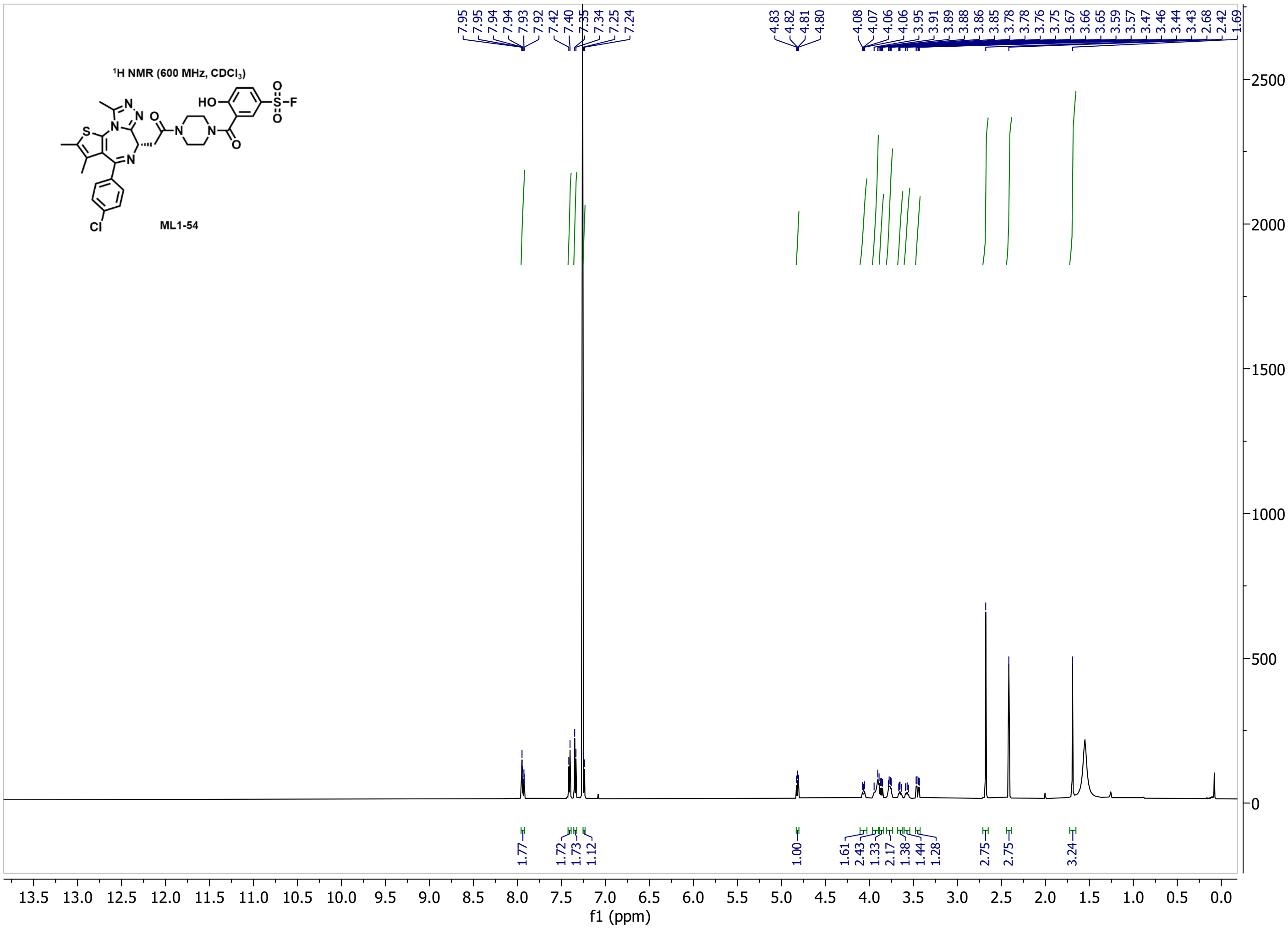

¹H NMR (600 MHz, CDCl₃)
ML1-54
7.95 7.95 7.94 7.94 7.93 7.92 7.42 7.40 7.35 7.34 7.25 7.24
4.83 4.82 4.81 4.80
4.08 4.07 4.06 4.06 3.95 3.91 3.89 3.88 3.86 3.85 3.78 3.78 3.76 3.75 3.67 3.66 3.65 3.59 3.57 3.47 3.46 3.44 3.43 2.68 2.42 1.69
1.77 1.72 1.73 1.12 1.00 1.61 2.43 1.33 2.17 1.38 1.44 1.28 2.75 2.75 3.24
f1 (ppm)


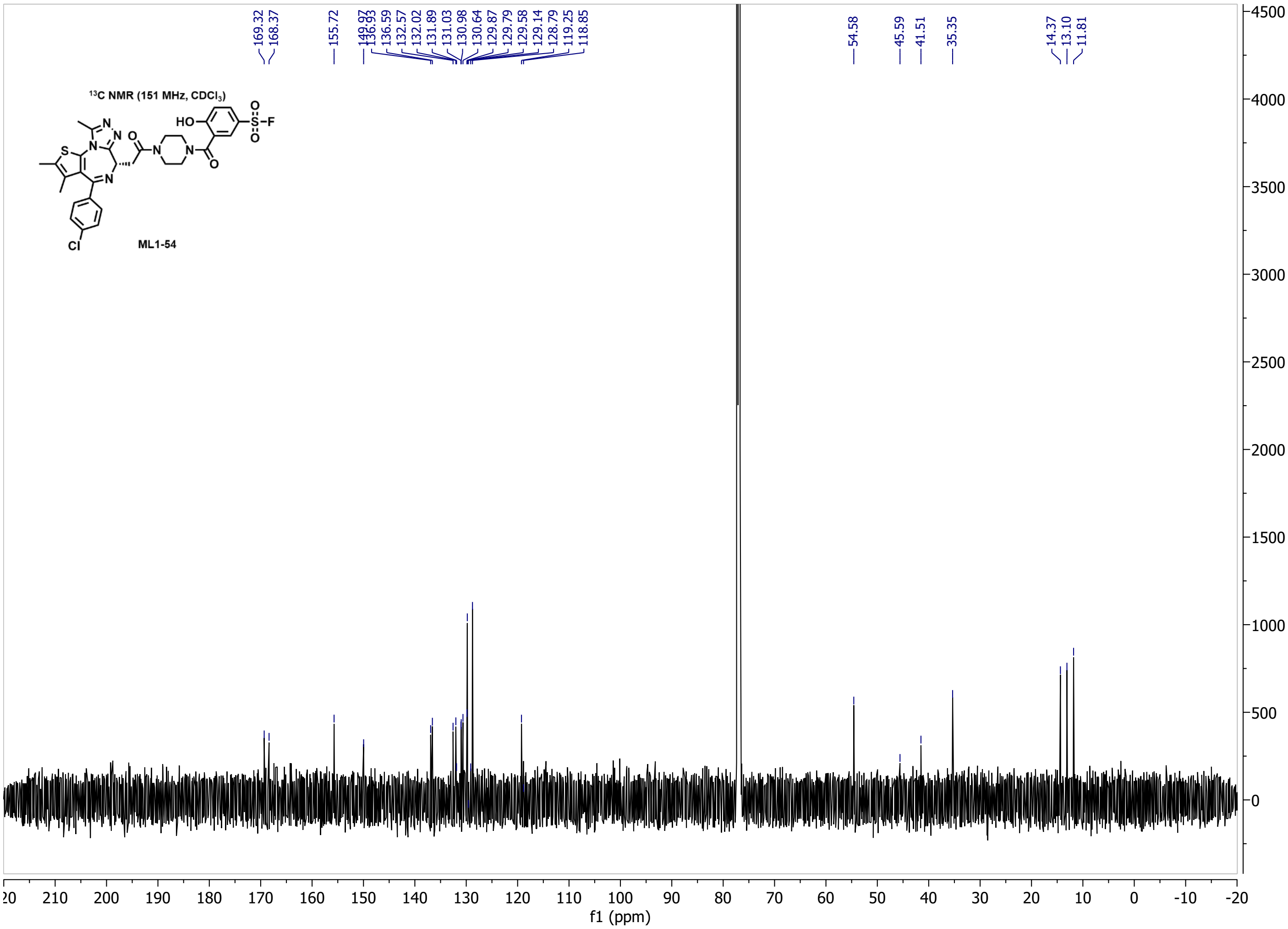

¹³C NMR (151 MHz, CDCl₃)
ML1-54
169.32 168.37 155.72 149.97 136.93 136.59 132.57 132.02 131.89 131.03 130.98 130.64 129.87 129.79 129.58 129.14 128.79 119.25 118.85
54.58 45.59 41.51 35.35 14.37 13.10 11.81
f1 (ppm)

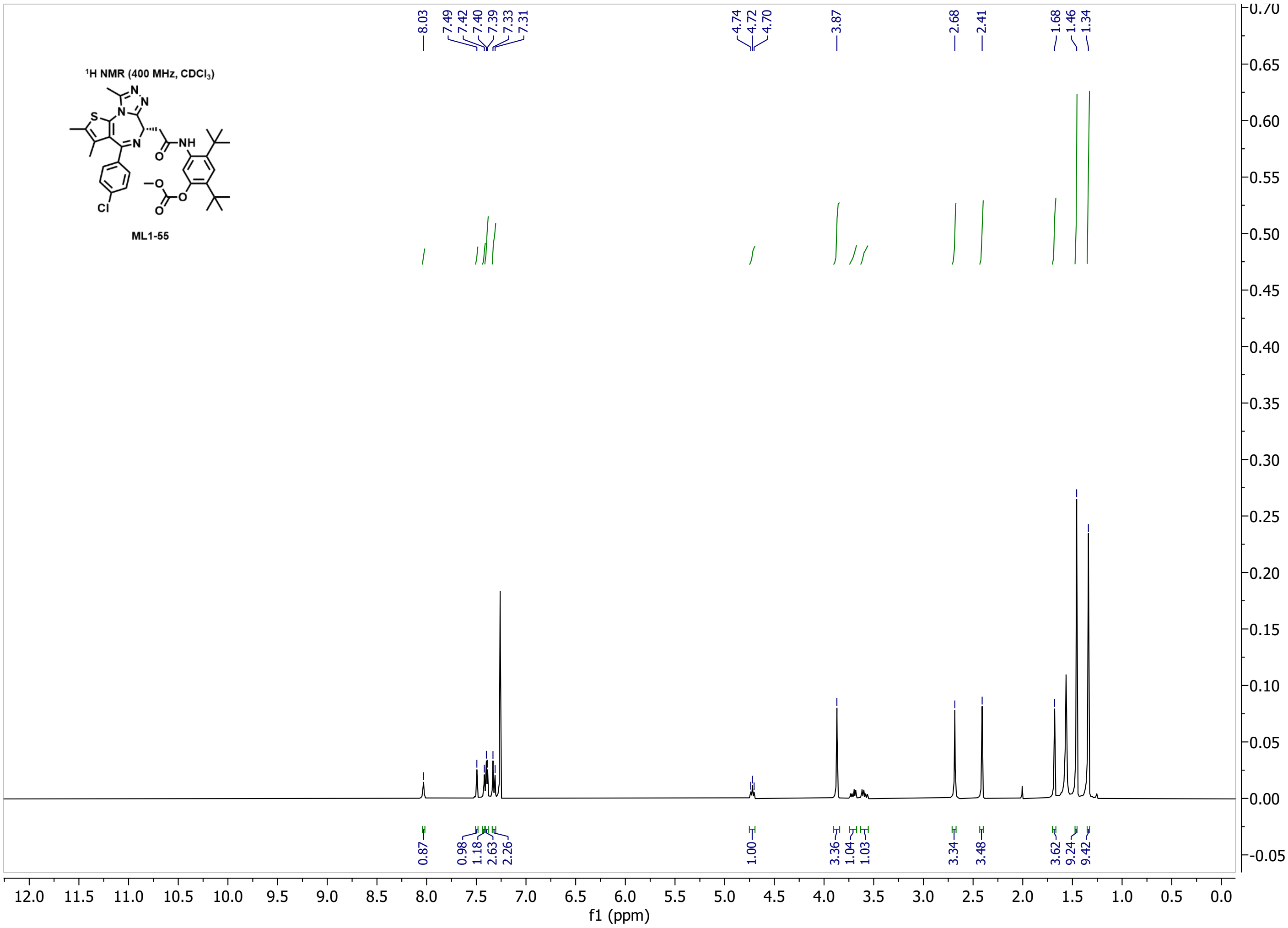

1H NMR (400 MHz, CDCl3)
ML1-55
8.03
7.49
7.42
7.40
7.39
7.33
7.31
4.74
4.72
4.70
3.87
2.68
2.41
1.68
1.46
1.34
0.87
0.98
1.18
2.63
2.26
1.00
3.36
1.04
1.03
3.34
3.48
3.62
9.24
9.42
f1 (ppm)


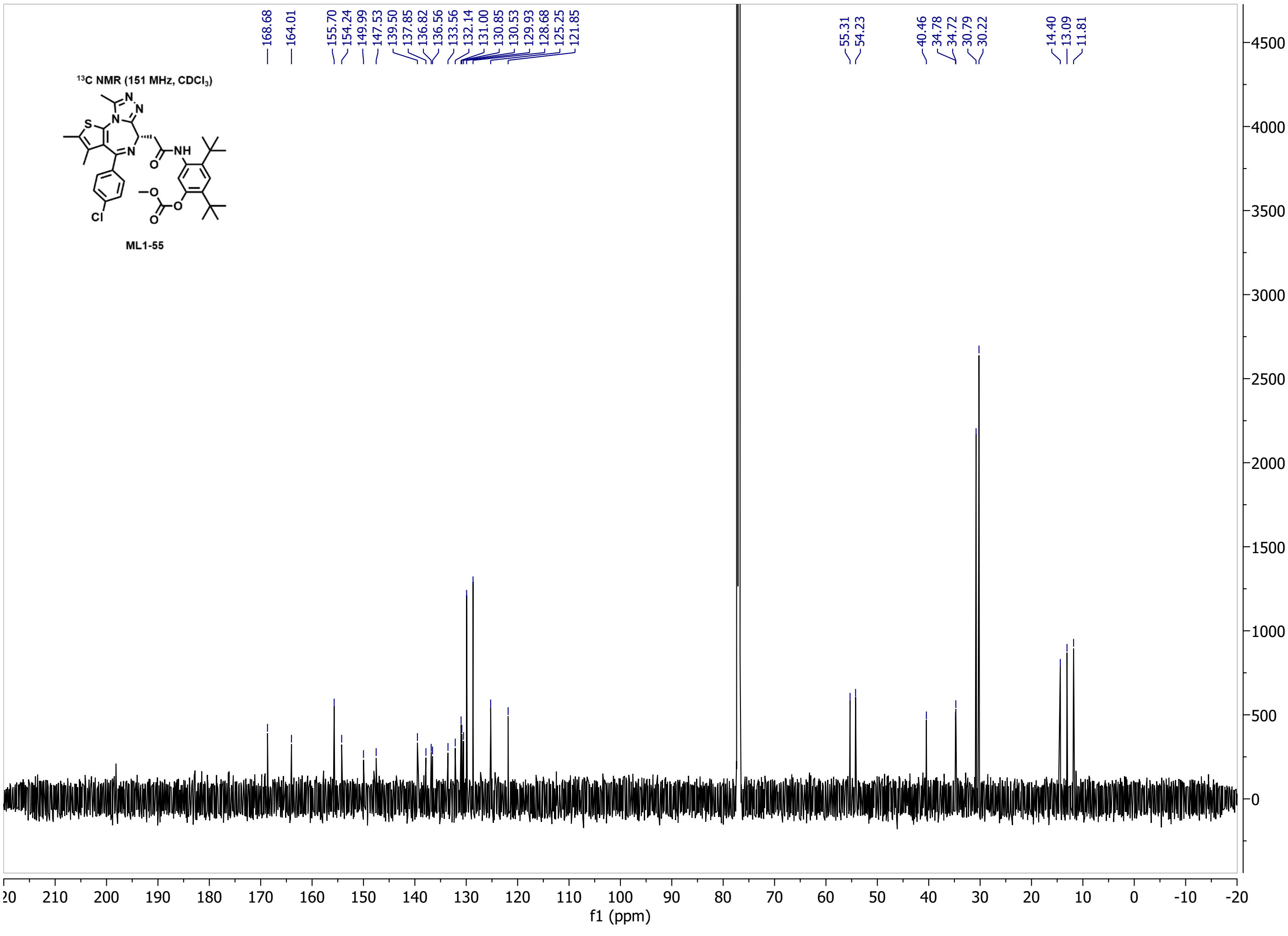

13C NMR (151 MHz, CDCl3)
ML1-55
168.68
164.01
155.70
154.24
149.99
147.53
139.50
137.85
136.82
136.56
133.56
132.14
131.00
130.85
130.53
129.93
128.68
125.25
121.85
55.31
54.23
40.46
34.78
34.72
30.79
30.22
14.40
13.09
11.81
f1 (ppm)

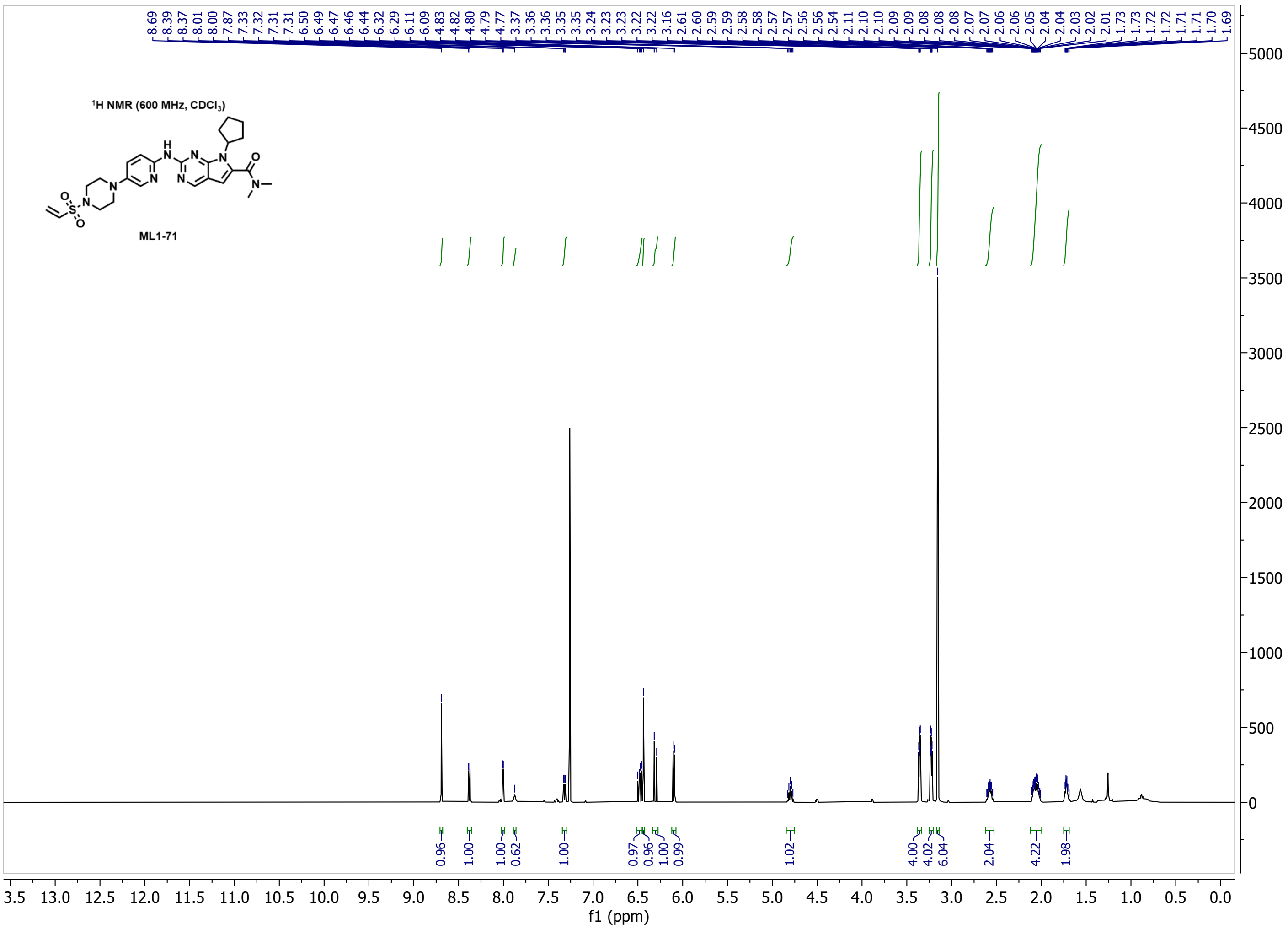

¹H NMR (600 MHz, CDCl₃)
ML1-71
8.69 8.39 8.37 8.01 8.00 7.87 7.33 7.32 7.31 7.31 6.50 6.49 6.47 6.46 6.44 6.32 6.29 6.11 6.09 4.83 4.82 4.80 4.79 4.77 3.37 3.36 3.36 3.35 3.35 3.24 3.23 3.23 3.22 3.22 3.16 2.61 2.60 2.59 2.59 2.58 2.58 2.57 2.57 2.56 2.56 2.54 2.11 2.10 2.10 2.09 2.09 2.08 2.08 2.08 2.07 2.07 2.06 2.06 2.05 2.04 2.04 2.03 2.02 2.01 1.73 1.73 1.72 1.71 1.71 1.70 1.69
0.96 1.00 1.00 0.62 1.00 0.97 0.96 1.00 0.99 1.02 4.00 4.02 6.04 2.04 4.22 1.98
f1 (ppm)


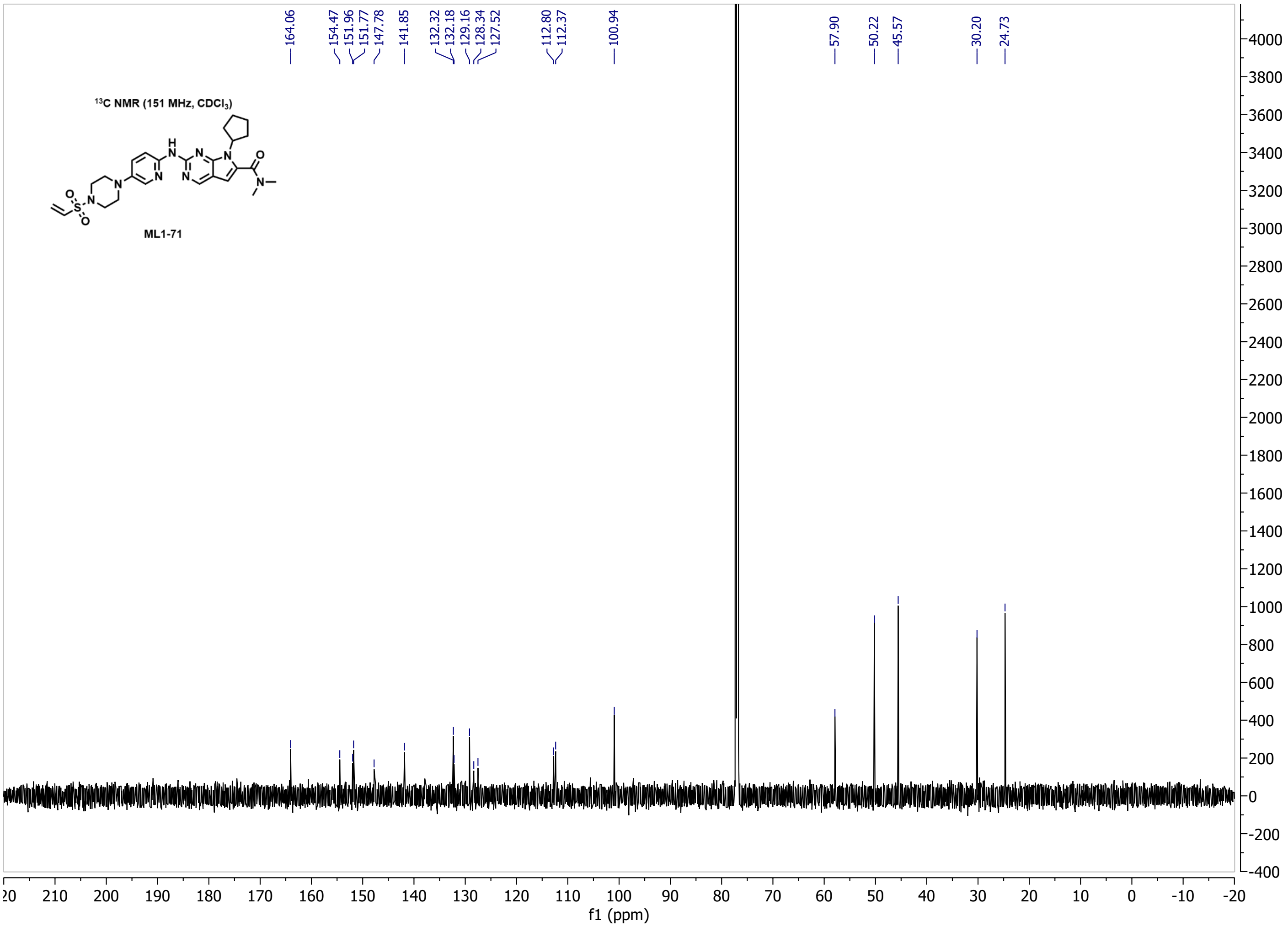

¹³C NMR (151 MHz, CDCl₃)
ML1-71
164.06 154.47 151.96 151.77 147.78 141.85 132.32 132.18 129.16 128.34 127.52 112.80 112.37 100.94 57.90 50.22 45.57 30.20 24.73
f1 (ppm)

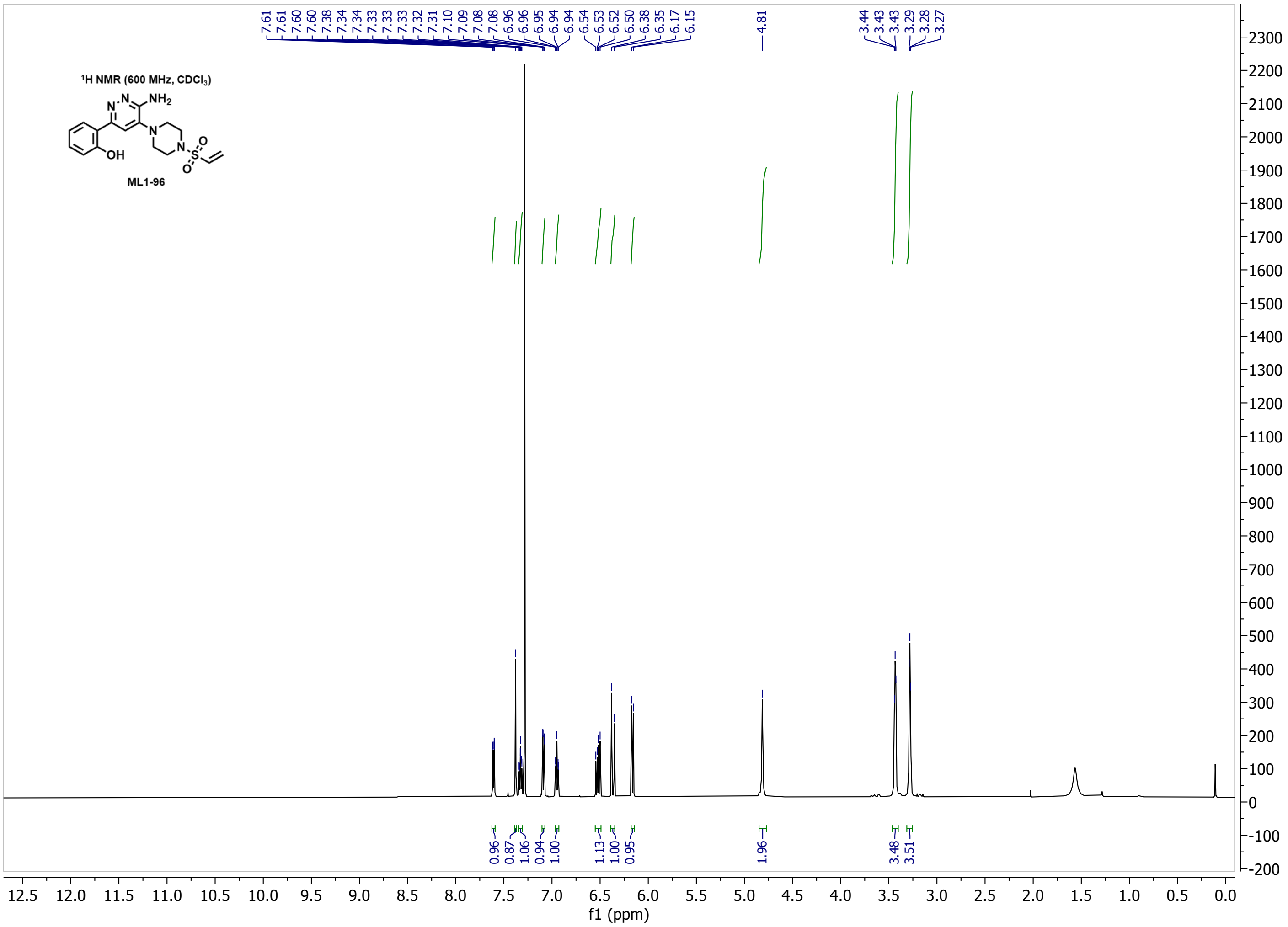
1H NMR (600 MHz, CDCl3)
NH2
OH
ML1-96
7.61 7.61 7.60 7.38 7.34 7.34 7.33 7.33 7.33 7.32 7.31 7.10 7.09 7.08 7.08 6.96 6.96 6.95 6.94 6.94 6.54 6.53 6.52 6.50 6.38 6.35 6.17 6.15
4.81
3.44 3.43 3.43 3.29 3.28 3.27
0.96 0.87 1.06 0.94 1.00 1.13 1.00 0.95 1.96 3.48 3.51
f1 (ppm)

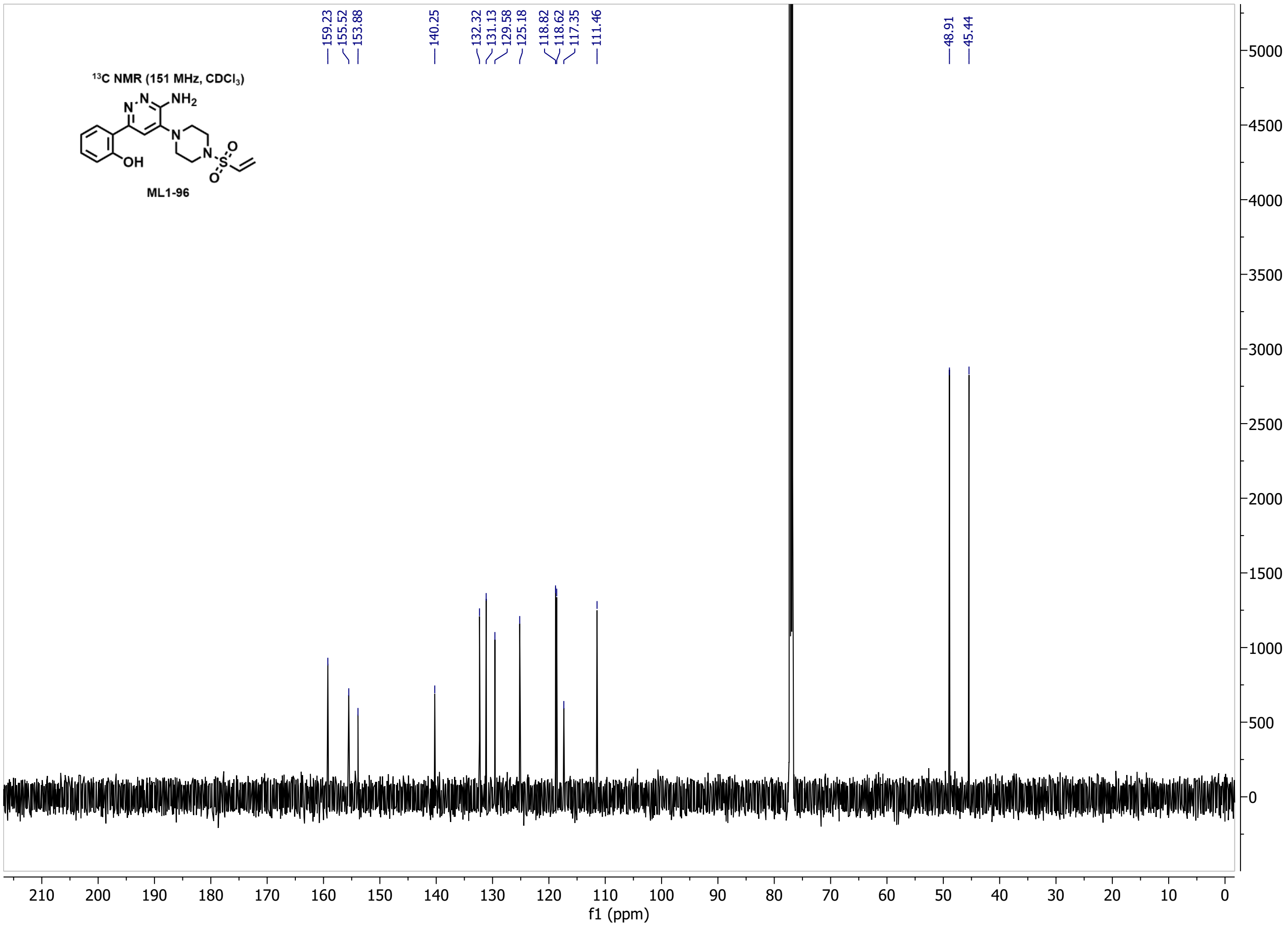
13C NMR (151 MHz, CDCl3)
NH2
OH
ML1-96
159.23 155.52 153.88
140.25
132.32 131.13 129.58 125.18
118.82 118.62 117.35 111.46
48.91 45.44
f1 (ppm)

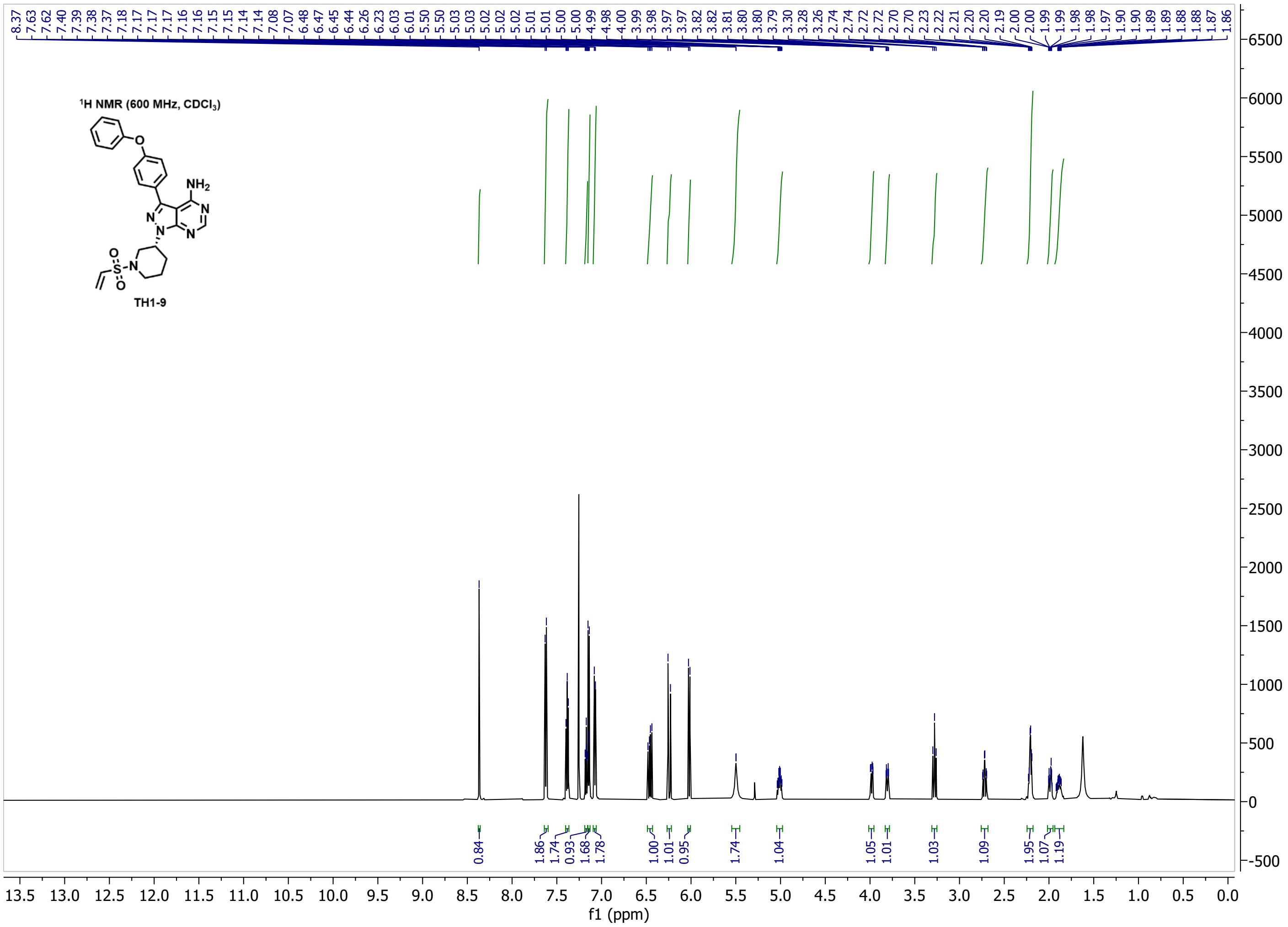

$^1$H NMR (600 MHz, CDCl$_3$)
TH1-9
f1 (ppm)


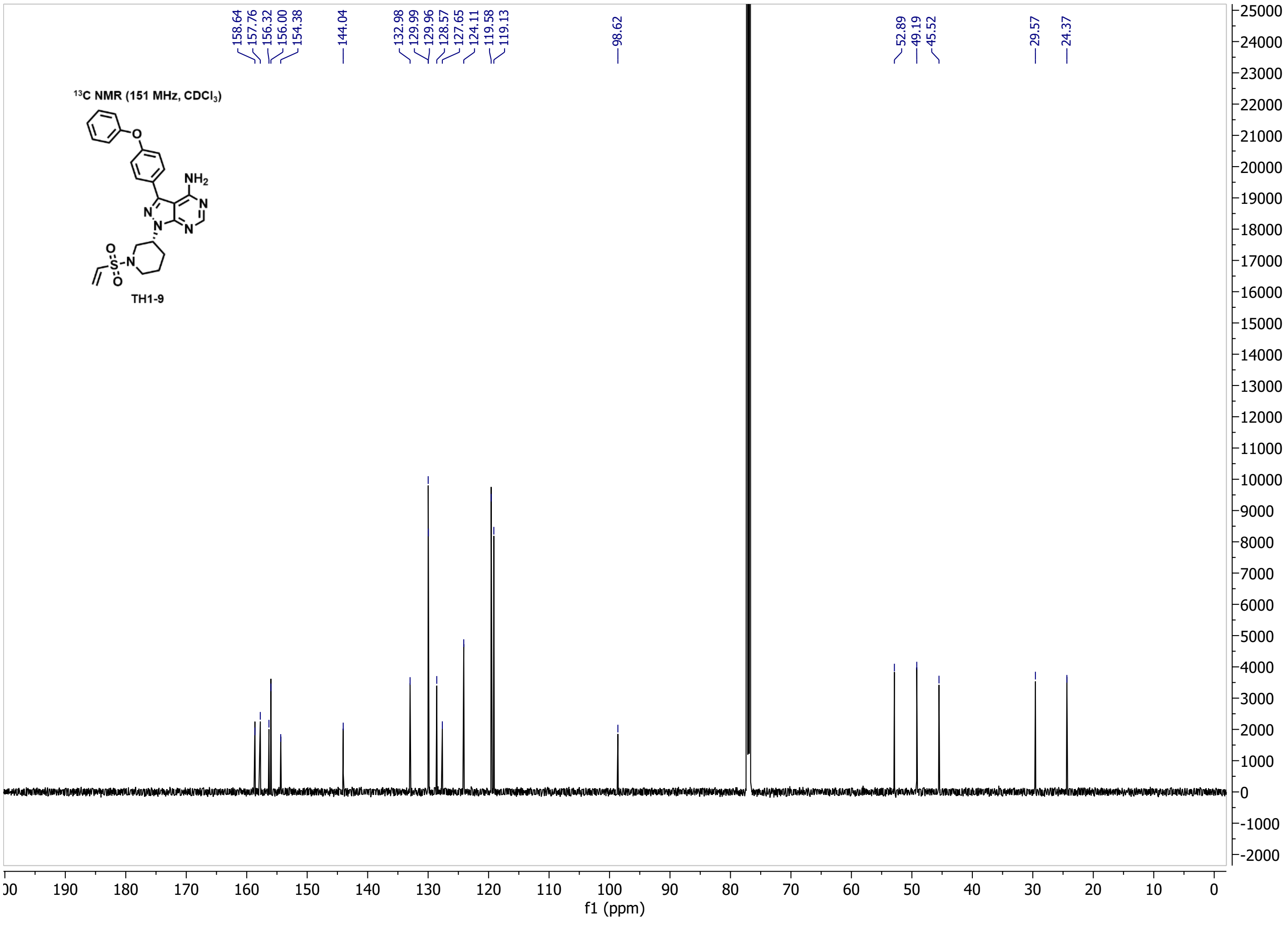

$^{13}$C NMR (151 MHz, CDCl$_3$)
TH1-9
f1 (ppm)

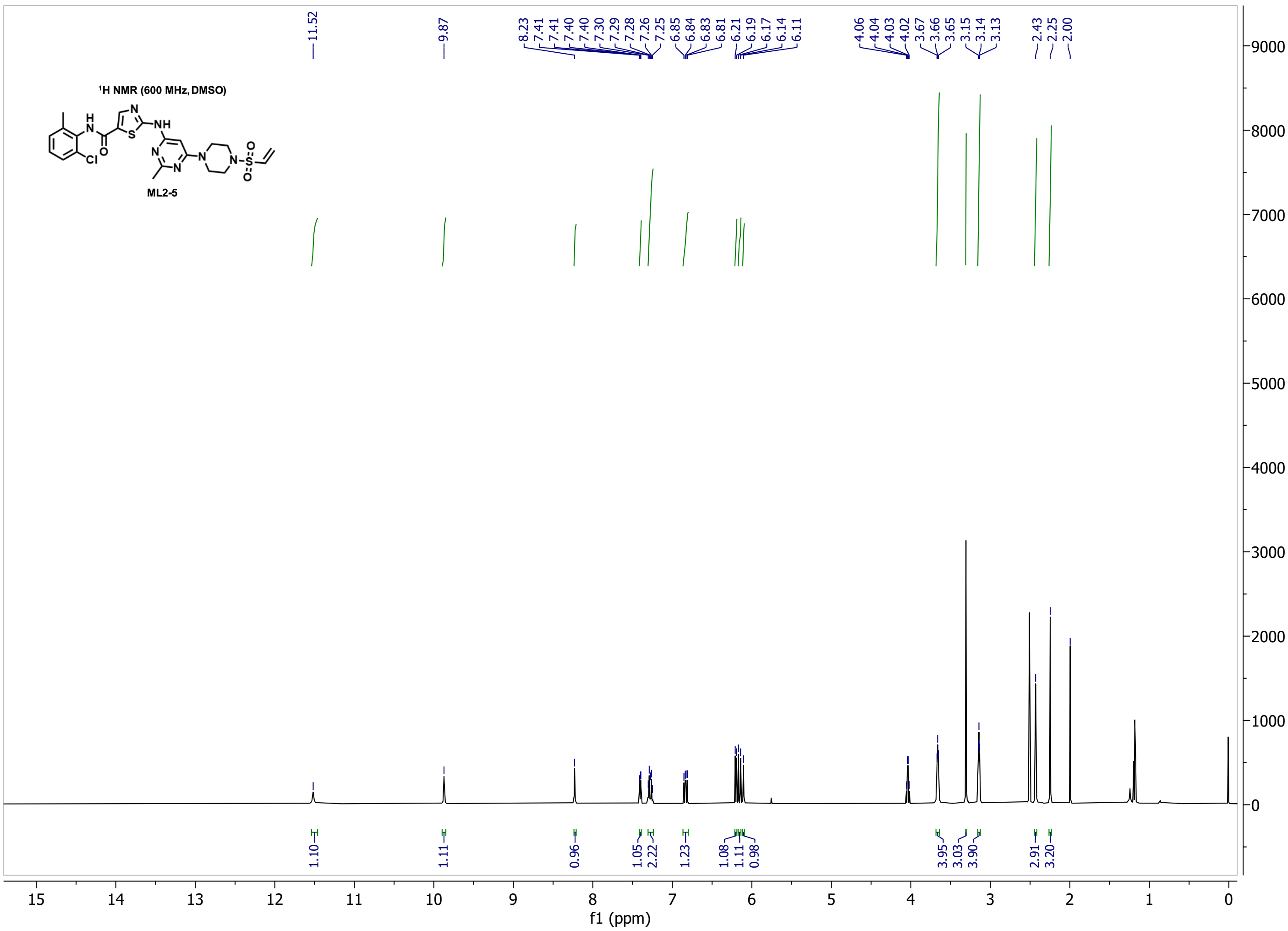

1H NMR (600 MHz, DMSO)
ML2-5
11.52
9.87
8.23 7.41 7.41 7.40 7.30 7.29 7.28 7.26 7.25 6.85 6.84 6.83 6.81 6.21 6.19 6.17 6.14 6.11
4.06 4.04 4.03 4.02 3.67 3.66 3.65 3.15 3.14 3.13
2.43 2.25 2.00
1.10 1.11 0.96 1.05 2.22 1.23 1.08 1.11 0.98 3.95 3.03 3.90 2.91 3.20
f1 (ppm)


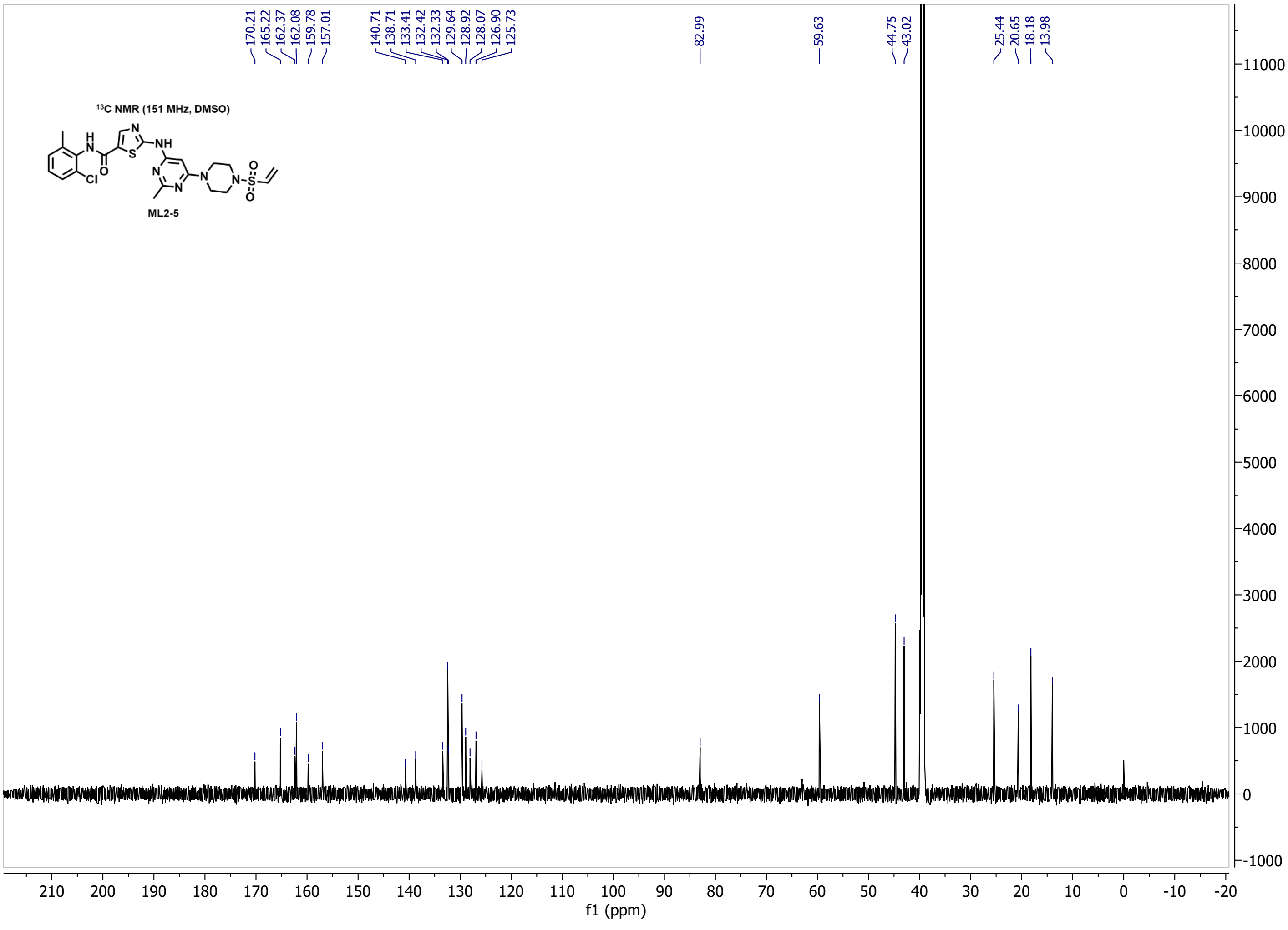

13C NMR (151 MHz, DMSO)
ML2-5
170.21 165.22 162.37 162.08 159.78 157.01
140.71 138.71 133.41 132.42 132.33 129.64 128.92 128.07 126.90 125.73
82.99
59.63
44.75 43.02
25.44 20.65 18.18 13.98
f1 (ppm)

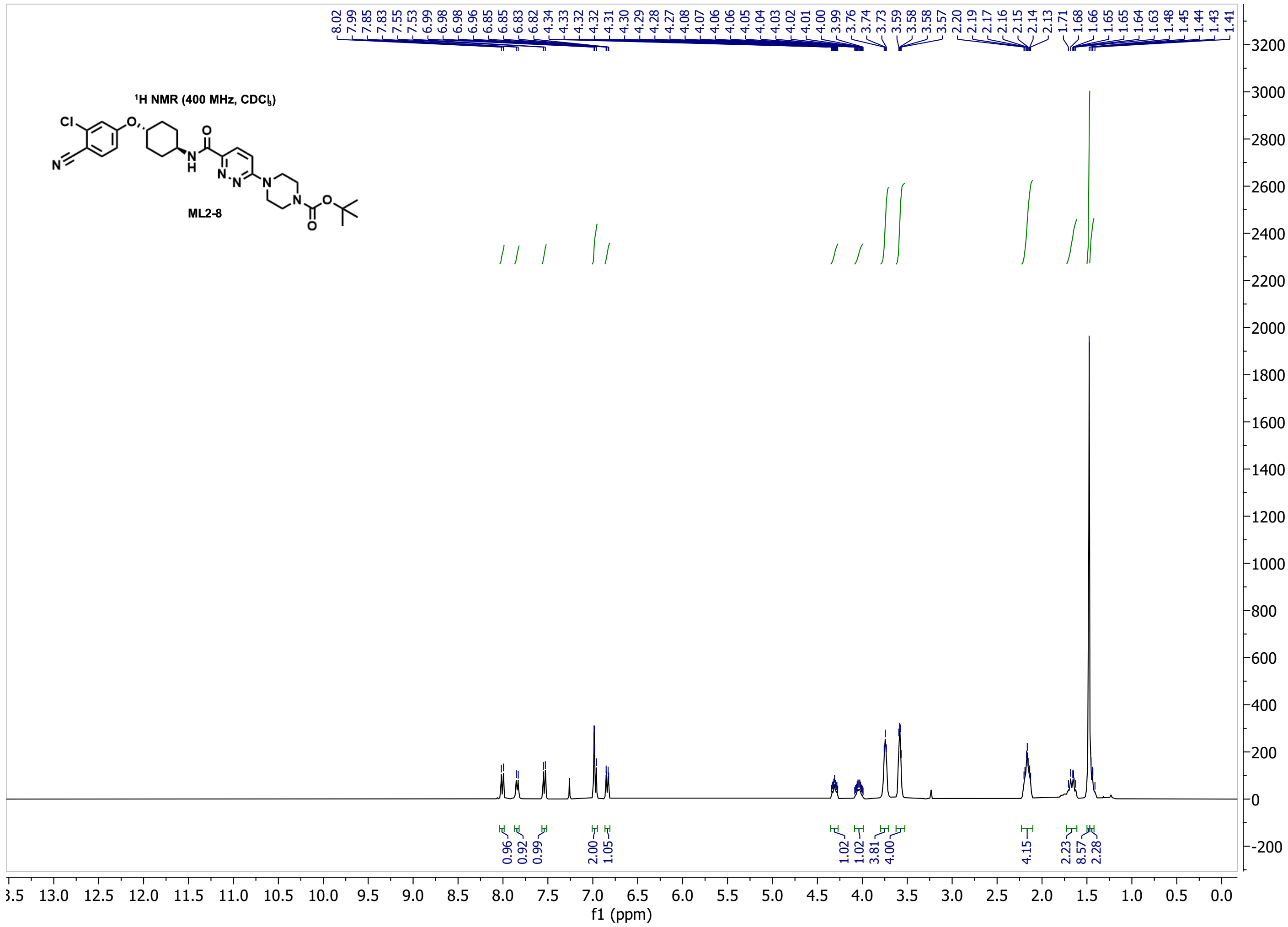

8.02 7.99 7.85 7.83 7.55 7.53 6.99 6.98 6.98 6.96 6.85 6.85 6.83 6.82 4.34 4.33 4.32 4.32 4.31 4.30 4.29 4.28 4.27 4.08 4.07 4.06 4.06 4.05 4.04 4.03 4.02 4.01 4.00 3.99 3.76 3.74 3.73 3.59 3.58 3.58 3.57 2.20 2.19 2.17 2.16 2.15 2.14 2.13 1.71 1.68 1.66 1.65 1.64 1.63 1.48 1.45 1.44 1.43 1.41
¹H NMR (400 MHz, CDCl₃)
Cl
O
N
H
N
N
N
N
O
O
O
ML2-8
0.96 0.92 0.99 2.00 1.05 1.02 1.02 3.81 4.00 4.15 2.23 8.57 2.28
f1 (ppm)


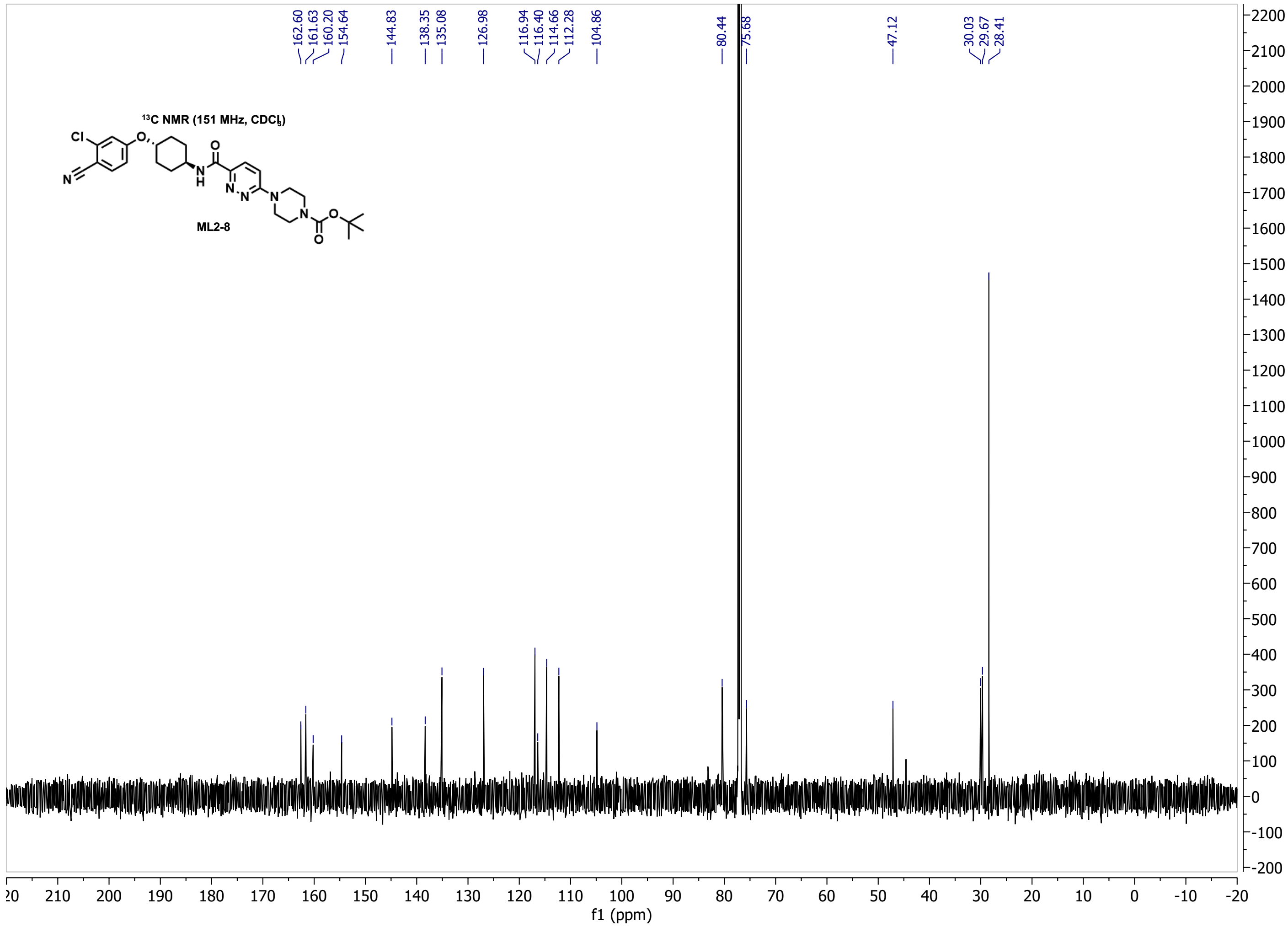

162.60 161.63 160.20 154.64 144.83 138.35 135.08 126.98 116.94 116.40 114.66 112.28 104.86 80.44 75.68 47.12 30.03 29.67 28.41
¹³C NMR (151 MHz, CDCl₃)
ML2-8
f1 (ppm)

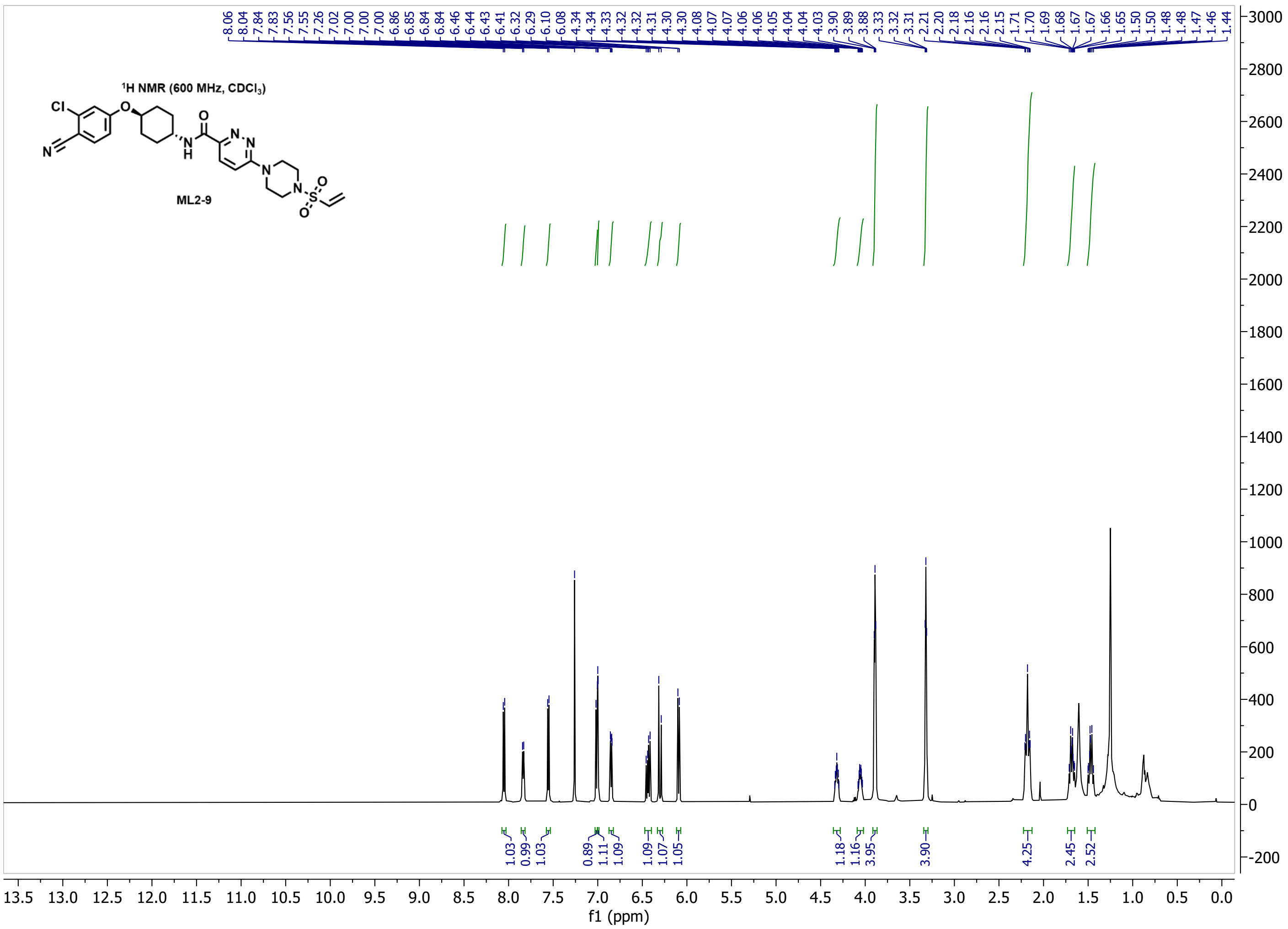
1H NMR (600 MHz, CDCl3)
ML2-9
Cl
O
N
H
N
N
N
S
O
O
8.06 8.04 7.84 7.83 7.56 7.55 7.26 7.02 7.00 7.00 7.00 6.86 6.85 6.84 6.84 6.46 6.44 6.43 6.41 6.32 6.29 6.10 6.08 4.34 4.34 4.33 4.32 4.32 4.31 4.30 4.30 4.08 4.07 4.06 4.06 4.05 4.04 4.04 4.03 3.90 3.89 3.88 3.33 3.32 3.31 2.21 2.20 2.18 2.16 2.16 2.15 1.71 1.70 1.69 1.68 1.67 1.67 1.66 1.65 1.50 1.50 1.48 1.48 1.47 1.46 1.44
1.03 0.99 1.03 0.89 1.11 1.09 1.09 1.07 1.05 1.18 1.16 3.95 3.90 4.25 2.45 2.52
f1 (ppm)

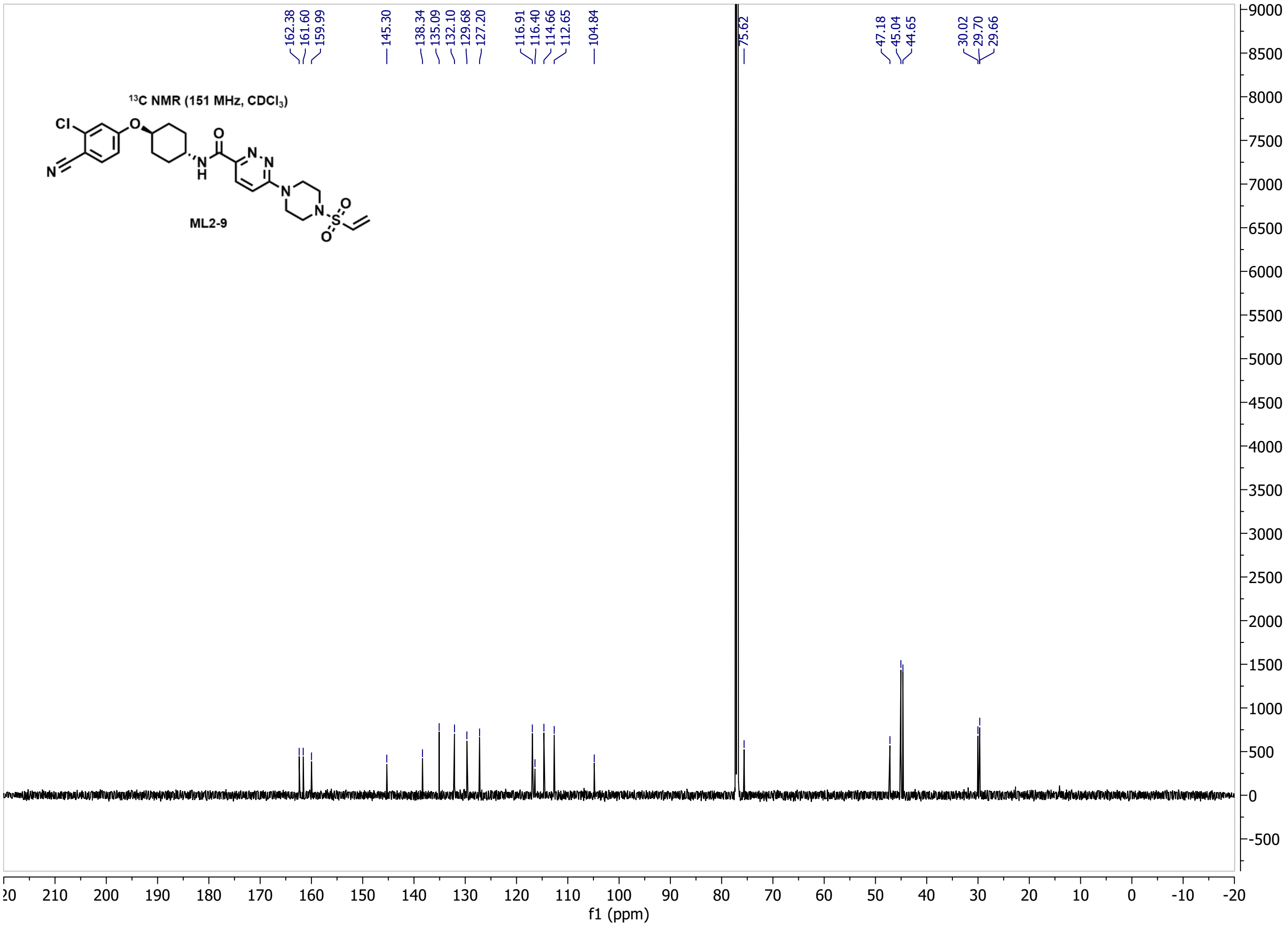
13C NMR (151 MHz, CDCl3)
ML2-9
Cl
O
N
H
N
N
N
S
O
O
162.38 161.60 159.99 145.30 138.34 135.09 132.10 129.68 127.20 116.91 116.40 114.66 112.65 104.84 75.62 47.18 45.04 44.65 30.02 29.70 29.66
f1 (ppm)

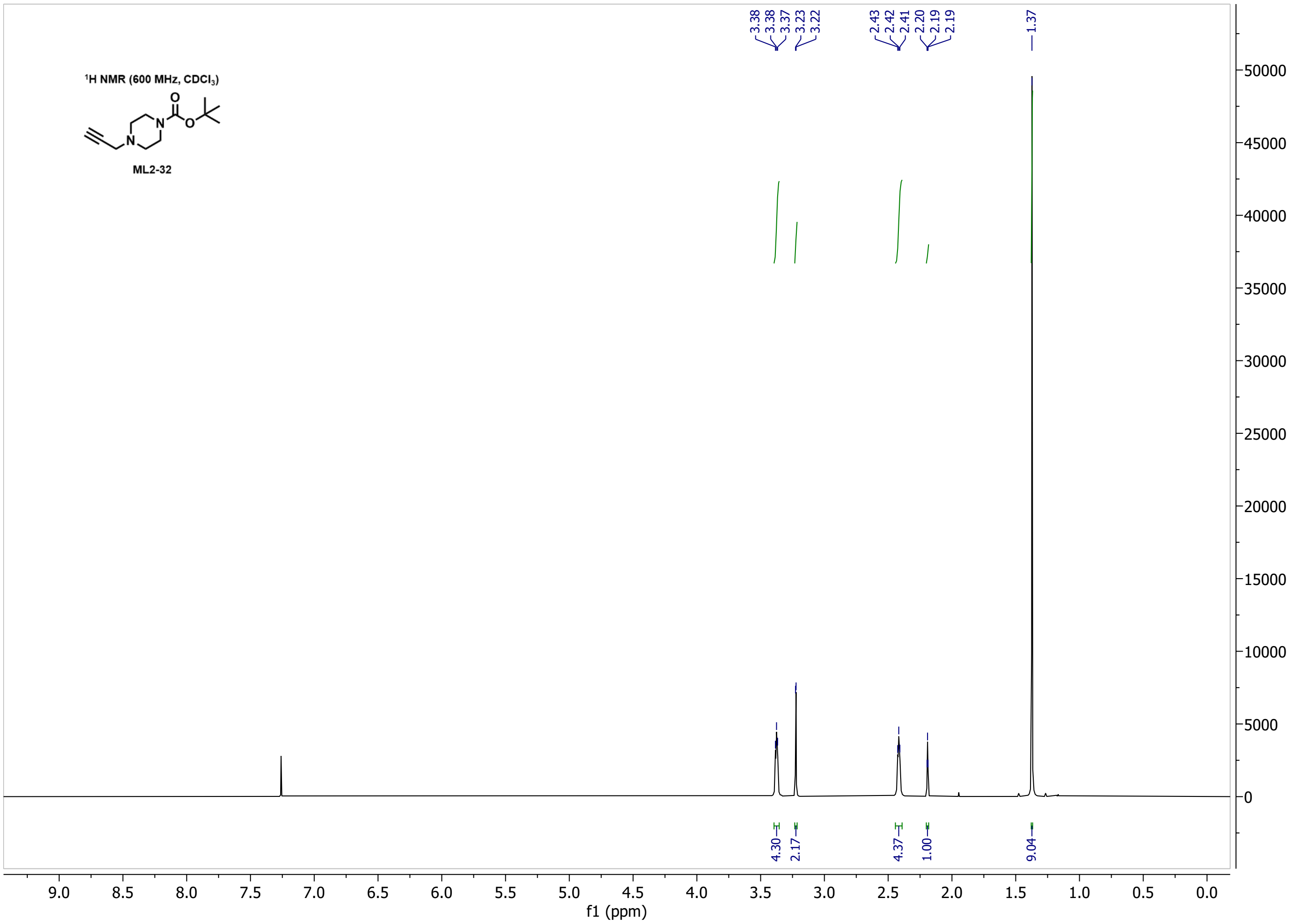

1H NMR (600 MHz, CDCl3)
ML2-32
3.38
3.38
3.37
3.23
3.22
2.43
2.42
2.41
2.20
2.19
2.19
1.37
4.30
2.17
4.37
1.00
9.04
f1 (ppm)


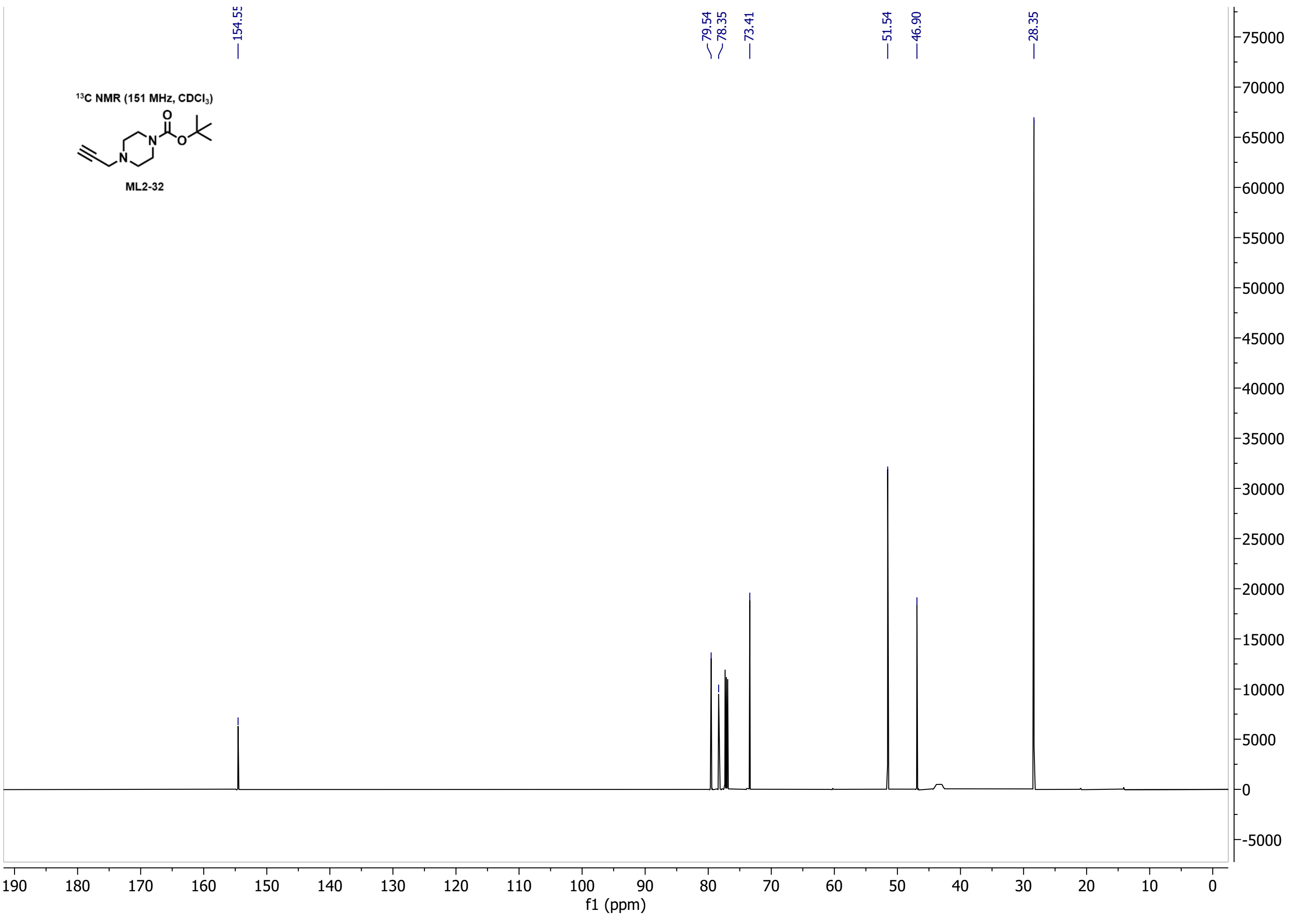

13C NMR (151 MHz, CDCl3)
ML2-32
154.55
79.54
78.35
73.41
51.54
46.90
28.35
f1 (ppm)

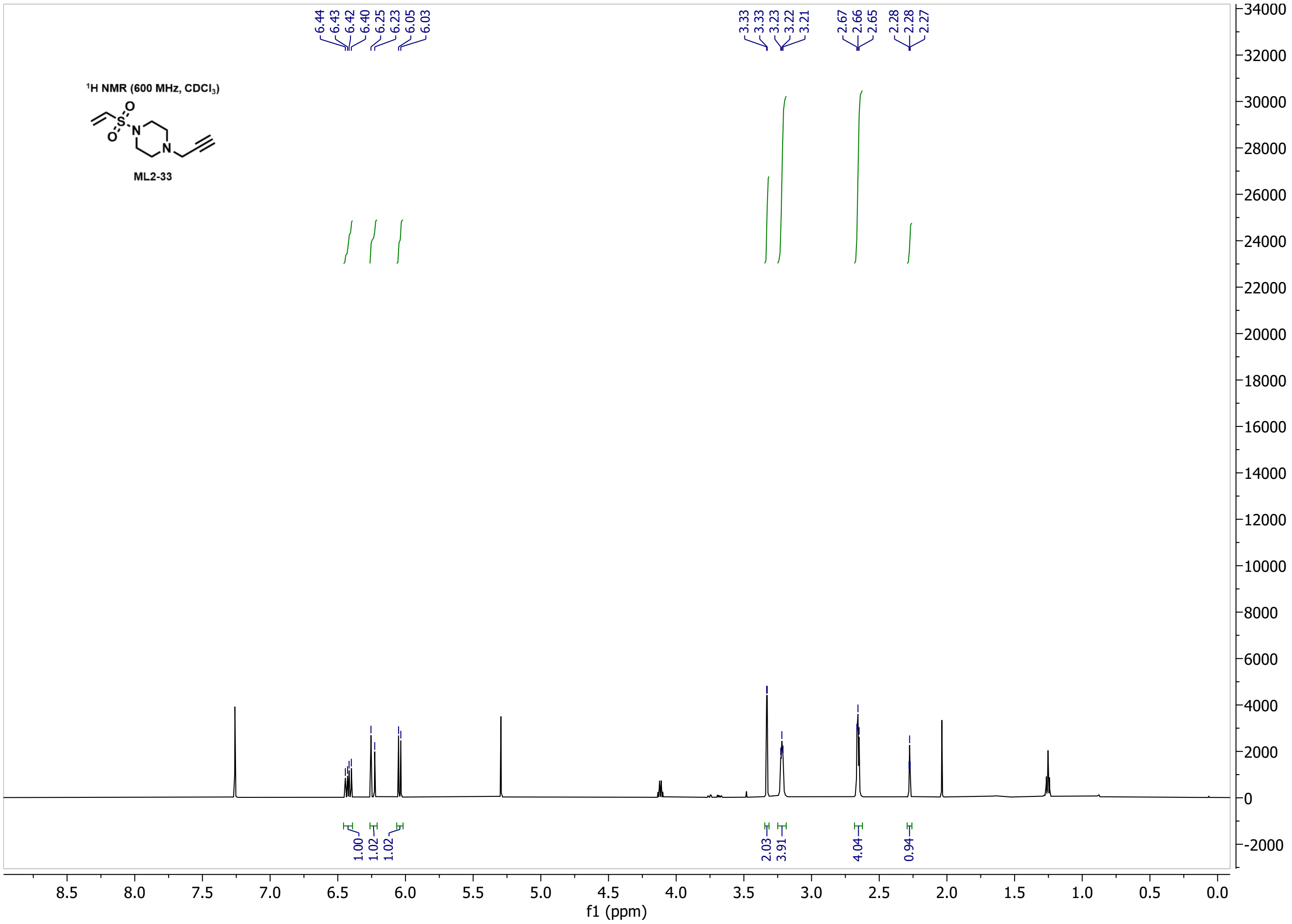

6.44
6.43
6.42
6.40
6.25
6.23
6.05
6.03
3.33
3.33
3.23
3.22
3.21
2.67
2.66
2.65
2.28
2.28
2.27
¹H NMR (600 MHz, CDCl3)
ML2-33
1.00
1.02
1.02
2.03
3.91
4.04
0.94
f1 (ppm)


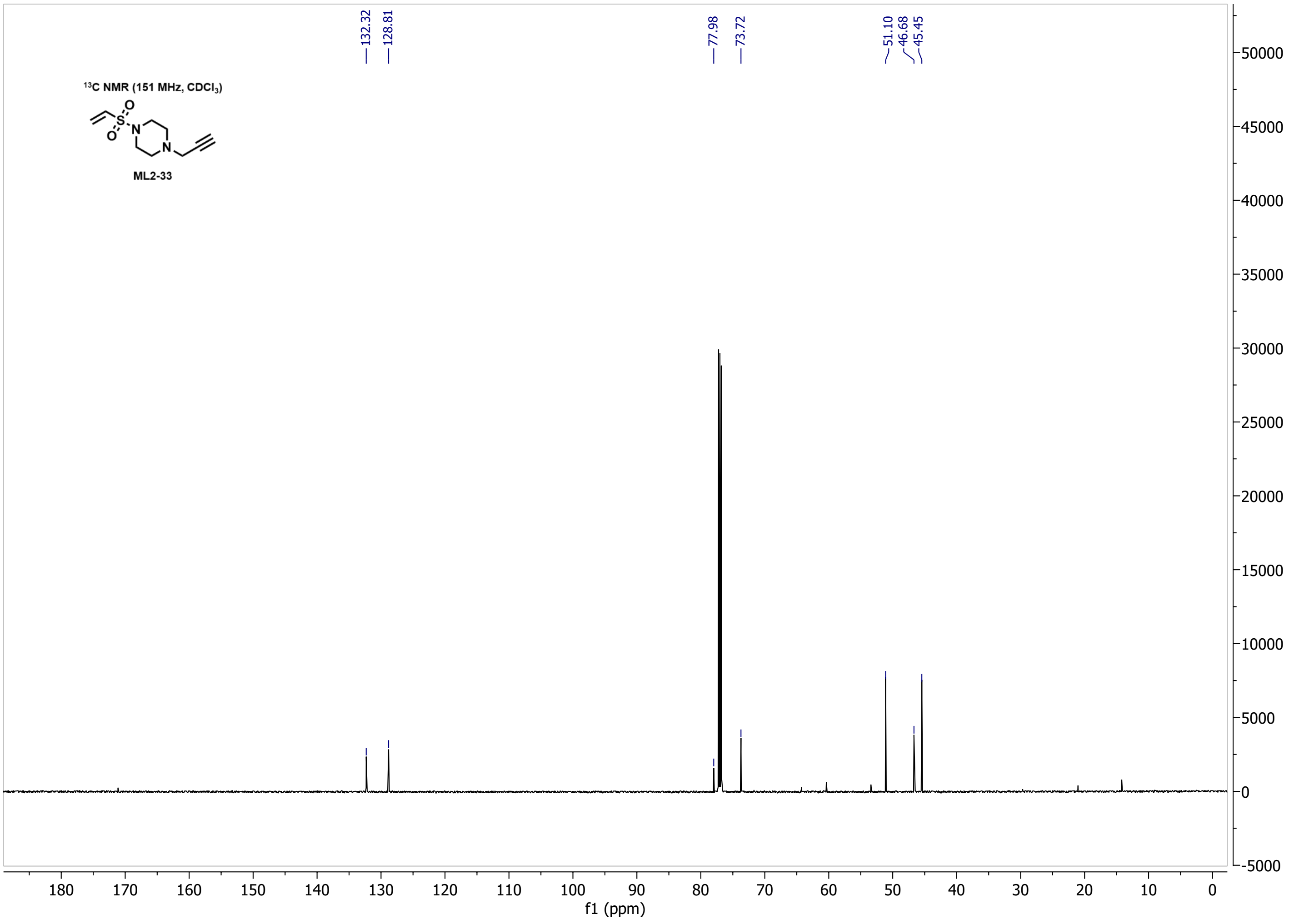

132.32
128.81
77.98
73.72
51.10
46.68
45.45
¹³C NMR (151 MHz, CDCl3)
ML2-33
f1 (ppm)
